# Plasticity and environment-specific relationships between gene expression and fitness in *Saccharomyces cerevisiae*

**DOI:** 10.1101/2024.04.12.589130

**Authors:** Mohammad A. Siddiq, Fabien Duveau, Patricia J. Wittkopp

## Abstract

Phenotypic evolution is shaped by interactions between organisms and their environments. The environment influences how an organism’s genotype determines its phenotype and how this phenotype affects its fitness. To better understand this dual role of the environment in the production and selection of phenotypic variation, we empirically determined and compared the genotype-phenotype-fitness relationship for mutant strains of the budding yeast *Saccharomyces cerevisiae* in four environments. Specifically, we measured how mutations in the promoter of the metabolic gene *TDH3* modified its expression level and affected its growth on media with four different carbon sources. In each environment, we observed a clear relationship between *TDH3* expression level and fitness, but this relationship differed among environments. Genetic variants with similar effects on *TDH3* expression in different environments often had different effects on fitness and vice versa. Such environment-specific relationships between phenotype and fitness can shape the evolution of phenotypic plasticity. The set of mutants we examined also allowed us to compare the effects of mutations disrupting binding sites for key transcriptional regulators and the TATA box, which is part of the core promoter sequence. Mutations disrupting the binding sites for the transcription factors had more variable effects on expression among environments than mutations disrupting the TATA box, yet mutations with the most environmentally variable effects on fitness were located in the TATA box. This observation suggests that mutations affecting different molecular mechanisms are likely to contribute unequally to regulatory sequence evolution in changing environments.

**Significance Statement:** Environments can affect the phenotypic traits an organism produces as well as the adaptive value of these traits (i.e. whether those traits will allow the organism to better survive and pass their genes on to the next generation). This study shows how the environment impacts both the production and selection of traits using the expression of a metabolic gene in the baker’s yeast as a model system. This study further shows that some types of genetic changes make gene expression traits more responsive to environmental changes than others, suggesting that genetic changes affecting different molecular mechanisms of gene regulation may contribute differently to genetic evolution.

## Introduction

An organism’s environment affects how its genotype determines its phenotype during the short-term process of development and how selection acts on that phenotype during the longer-term process of evolution (Fig. 1*A*). Often, a single genotype can produce different phenotypes in different environments, which is known as phenotypic plasticity (1, 2). Phenotypic plasticity is itself a variable and evolvable trait, and the same environmental change can induce different phenotypic changes among genetically distinct organisms through gene-by-environment (GxE) interactions (1, 3–8). The differences in reproductive success conferred by diverse phenotypes - known as relative fitness - can also differ among environments because some trait values can be better suited to one environment than another (9–17). This relationship between phenotypes and fitness, which can be described as a 2-dimensional fitness function when considering variation in one particular trait (18, 19), determines which individuals are most likely to reproduce and influence evolution (Fig. 1*A*, gray arrow).

**Figure 1.**
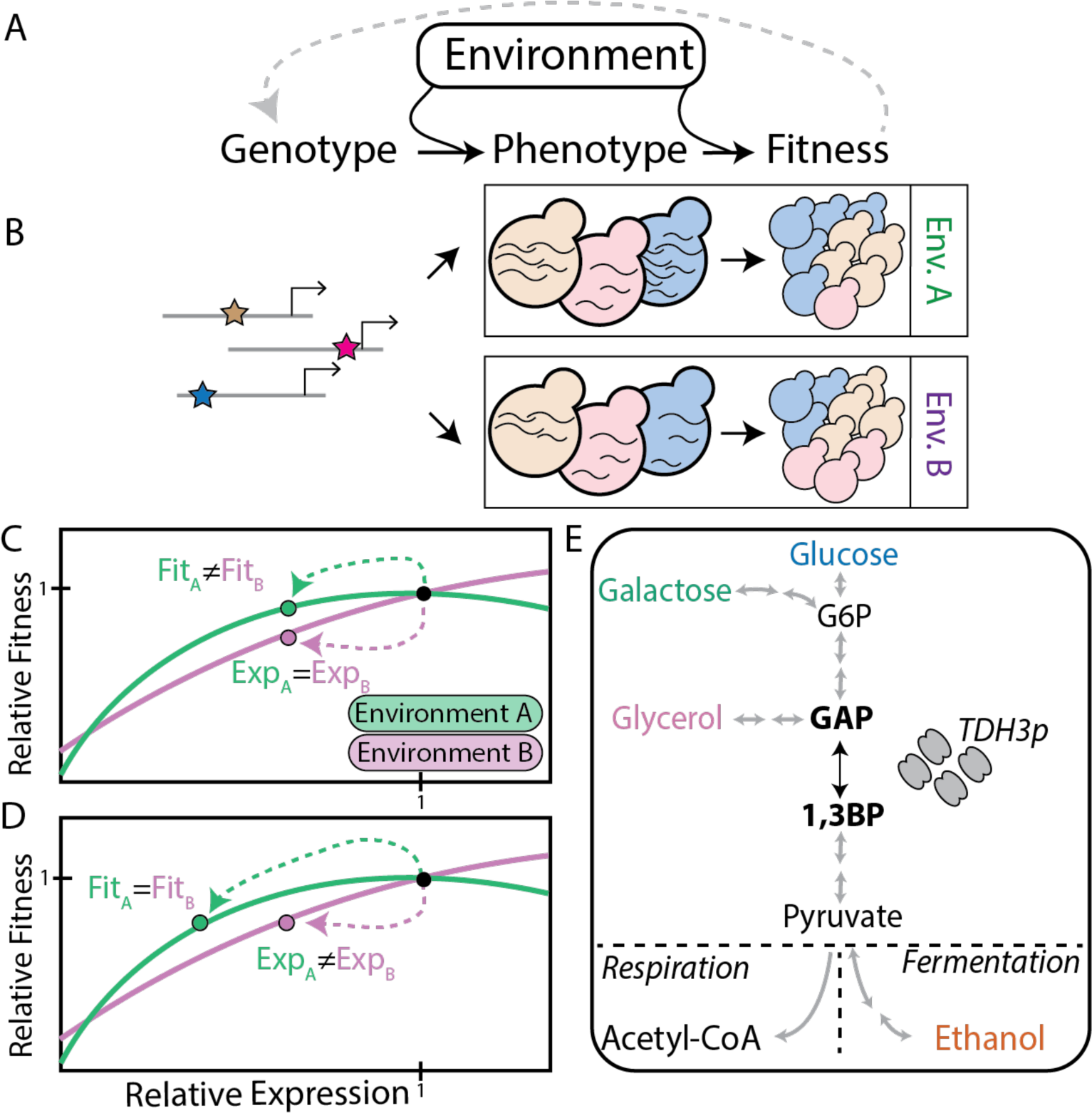
Environmental impacts on genotype-phenotype-fitness relationships. (A) The schematic shows that Genotype and Environment both influence the development of a Phenotype and that Phenotype and Environment together determine an organism’s relative Fitness. The grey dotted line connecting Fitness to Genotype reflects reproduction and potential evolutionary changes in allele frequencies from one generation to the next. (B) Allelic variation for a gene’s promoter (with arrowheads indicating the gene’s transcription start site) is shown under Genotype, leading to variation in the gene’s expression (shown with curvy lines representing differences in RNA abundance in budding yeast cells) shown under Phenotype, and variation in the abundance of different yeast genotypes resulting from their differences in gene expression shown under Fitness. Hypothetical differences in RNA abundance and relative fitness between two environments (Env. A and Env. B) are also shown. (C, D) Solid lines show the relationship between a focal gene’s expression and fitness in two different environments (green = environment A; purple = environment B). The black dot shows the expression and fitness of a “wild-type” promoter allele, with both values defined as 1. The green and purple dots show the expression and fitness of the same alternate promoter allele in environment A (green) and environment B (purple) with expression and fitness defined relative to the wild-type allele. The green and purple dotted arrows highlight the effects of genetic differences between the wild-type and alternate promoter alleles on gene expression and fitness in each environment. Panel C shows a case where the genetic difference between the wild-type and alternate allele (e.g., a new mutation) has the same effect on the focal gene’s expression in both environments (Exp_A_ = Exp_B;_ no phenotypic plasticity) but different effects on fitness (Fit_A_ ≠ Fit_B_) due to differences in the shape of the fitness function between environments. Panel D shows a case where the genetic difference between the wild-type and alternate allele (e.g., a new mutation) has different effects on the focal gene’s expression in the two environments (Exp_A_ ≠ Exp_B;_ phenotypic plasticity) but the same effect on fitness (Fit_A_ ≠ Fit_B_) due to differences in the shape of the fitness function between environments. Note that if the wild-type and alternate alleles are considered together, the case shown in C exhibits a gene-by-environment interaction for fitness because the alternate allele shows differences in fitness between the two environments but the wild-type allele does not; this case does not show a gene-by-environment interaction for gene expression because expression is the same for both alleles in the two environments (SI Appendix Fig. S1*A,B*). The opposite is true for the case shown in panel D: a gene-by-environment interaction is present for gene expression but not fitness (SI Appendix Fig. S1C,D). (E) The abbreviated metabolic pathway shown illustrates how the carbon sources used in this study (glucose, galactose, glycerol, and ethanol) relate to the glycolysis/gluconeogenesis pathway. The enzyme encoded by the *TDH3* gene (*TDH3p*) catalyzes the interconversion of glyceraldehyde-3-phoshate (GAP) to 1,3-biphosphoglycerate (1,3BP), as shown by the double-headed black arrow. Gray arrows highlight other metabolic steps; multiple gray arrows indicate reactions catalyzed by two or more enzymes.

The importance of the dual role of the environment in the production and selection of phenotypic variation is widely recognized (1, 20). For example, empirical studies have demonstrated that phenotypic plasticity can be adaptive (9, 11, 14, 16, 17, 21, 22), such as when dorsal head spikes that increase fitness develop in *Daphnia* water fleas in response to predator-associated chemicals (9, 17). However, phenotypic plasticity can also be maladaptive (12, 23–25): for example, mice adapted to low altitude conditions have a physiological response to low oxygen conditions that also causes overproduction of red blood cells and potential pulmonary hypertension in high altitude conditions (25). Theoretical studies have articulated evolutionary scenarios in which plasticity may be selectively maintained or removed (26, 27) and how genotype-by-environment interactions may allow the accumulation of genetic variants that only impact phenotype under certain conditions (28, 29). However, the genetic and molecular underpinnings of phenotypic plasticity–and how variable environments shape evolution of those genetic and molecular mechanisms–remain poorly understood (8, 14, 30). For example, are mutations disrupting certain types of molecular processes more prone to producing environment-dependent phenotypes and thereby more likely to be conditionally beneficial or deleterious? And, how do environmental effects of mutations on phenotype and fitness compare, given that the environment can change both the phenotypes produced and the nature of selection acting on those phenotypes?

To understand how changes in the environment can affect phenotypic evolution, we need to know how specific genetic changes impact a focal trait in different environments and how the fitness of the resulting phenotypes differs among those same environments. Such experiments are now possible in microbial systems (31, 32), and gene expression in yeast is a powerful focal phenotype for this work for several reasons (31, 33, 34). First, changes in gene expression can impact cellular function (and thus fitness), and such changes in gene expression are primarily responsible for plasticity in downstream phenotypes. Second, genetic changes that affect expression of a gene can be easily generated via site-directed mutagenesis of the promoter sequence located upstream the gene’s coding sequence. Third, the quantitative effects of promoter mutations on the focal gene’s expression can be measured with high precision and throughput using fluorescent reporters or RNA-seq. Fourth, the impact of mutations on fitness in the unicellular eukaryote *Saccharomyces cerevisiae* can be directly and precisely measured by comparing the relative growth rate of strains differing only by the mutations of interest. Finally, the environment of yeast cells can be easily controlled and changed in the laboratory.

By characterizing the expression and fitness of a set of yeast strains that differ only by mutations in the promoter of a focal gene in multiple environmental conditions (Fig. 1*B*), we can tease apart the effects of the environment on each mutation’s expression level and relative fitness. We can then interpret the environment-specific relationship between expression level and fitness using empirically determined fitness functions for the different environments. For example, if the fitness function (i.e., the relationship between the focal gene’s expression and fitness) differs between two environments, a mutation could have the same effect on expression in both environments but different effects on fitness (Fig. 1*C*; *SI Appendix* Fig. 1*A*-*B*), different effects on expression in the two environments but the same effect on fitness (Fig. 1*D*; *SI Appendix* Fig. 1*C*-*D*), or some other combination of effects on expression and fitness.

Here, we use *TDH3* in *S. cerevisiae* as a focal gene to examine the effects of promoter mutations on gene expression and fitness in four environments: media containing glucose, galactose, glycerol, or ethanol as a carbon source. Glucose and galactose are fermentable carbon sources that can be metabolized by *S. cerevisiae* in aerobic and anaerobic conditions via glycolysis and alcoholic fermentation, with galactose first requiring degradation via the Leloir pathway. Glycerol and ethanol are non-fermentable carbon sources that can only be metabolized by *S. cerevisiae* cells in aerobic conditions. The *TDH3* gene encodes a *GAPDH* protein that catalyzes the interconversion of glyceraldehyde-3-phosphate and 1,3-biphosphoglycerate, and this biological role makes the gene’s activity important for the metabolism of different carbon sources via glycolysis and gluconeogenesis (Fig. 1*E*). Past studies have demonstrated that some mutations in this promoter have effects on expression levels that vary among different carbon environments (35), and studies in rich glucose media have shown that variation in *TDH3* promoter activity affects organismal fitness (36, 37), but it has remained unknown how the relationship between expression level and fitness changes with the environment and, in turn, how the effects of mutations on expression and fitness vary among environments.

To address this knowledge gap, we use 51 strains of *S. cerevisiae* carrying different alleles of the *TDH3* promoter to empirically characterize the concurrent effects of the environment on expression level and fitness in four environments. We first use these data to construct environment-specific fitness functions relating *TDH3* expression levels to relative growth rate, our proxy for fitness. Using these fitness functions, we then assess the relationship between environment-dependent effects on expression and relative fitness for promoter variants. We find that both plasticity and fitness functions vary among environments, with plasticity being beneficial in some contexts and detrimental in others. We also ask whether plasticity differs for mutations that impact transcription through different molecular mechanisms by comparing the effects of mutations in specific transcription factor binding sites and mutations in the TATA box on which the core transcriptional preinitiation complex assembles. We found that mutations in specific transcription factor binding sites generally had more variable effects on expression among environments than mutations in the TATA box; however, the mutations with the most variable effects on fitness among environments occurred in the TATA box. Together, these data show how the environment jointly impacts the production and selection of phenotypic variation, reveal different environment-dependent effects of mutations that regulate gene expression through distinct molecular mechanisms, and suggest a mechanistic explanation for propensities in regulatory sequence evolution as organisms navigate life in different environments.

## Results

### Environmental impacts on *TDH3* expression and its contribution to growth rate

Different environments pose different demands for growth on cells. *S. cerevisiae* can grow on a wide range of carbon sources but it does so at different rates. To determine how the environments used in this study impact the growth of *S. cerevisiae*, we measured the growth rate of a haploid reference strain using batch cultures in four types of rich media with each containing a different fermentable (glucose or galactose) or non-fermentable (glycerol or ethanol) carbon source. As expected given that adaptation to glucose is a hallmark of *S. cerevisiae*, this strain grew more rapidly on fermentable than non-fermentable carbon sources and had the highest growth rate on glucose (Fig. 2*A*).

**Figure 2:**
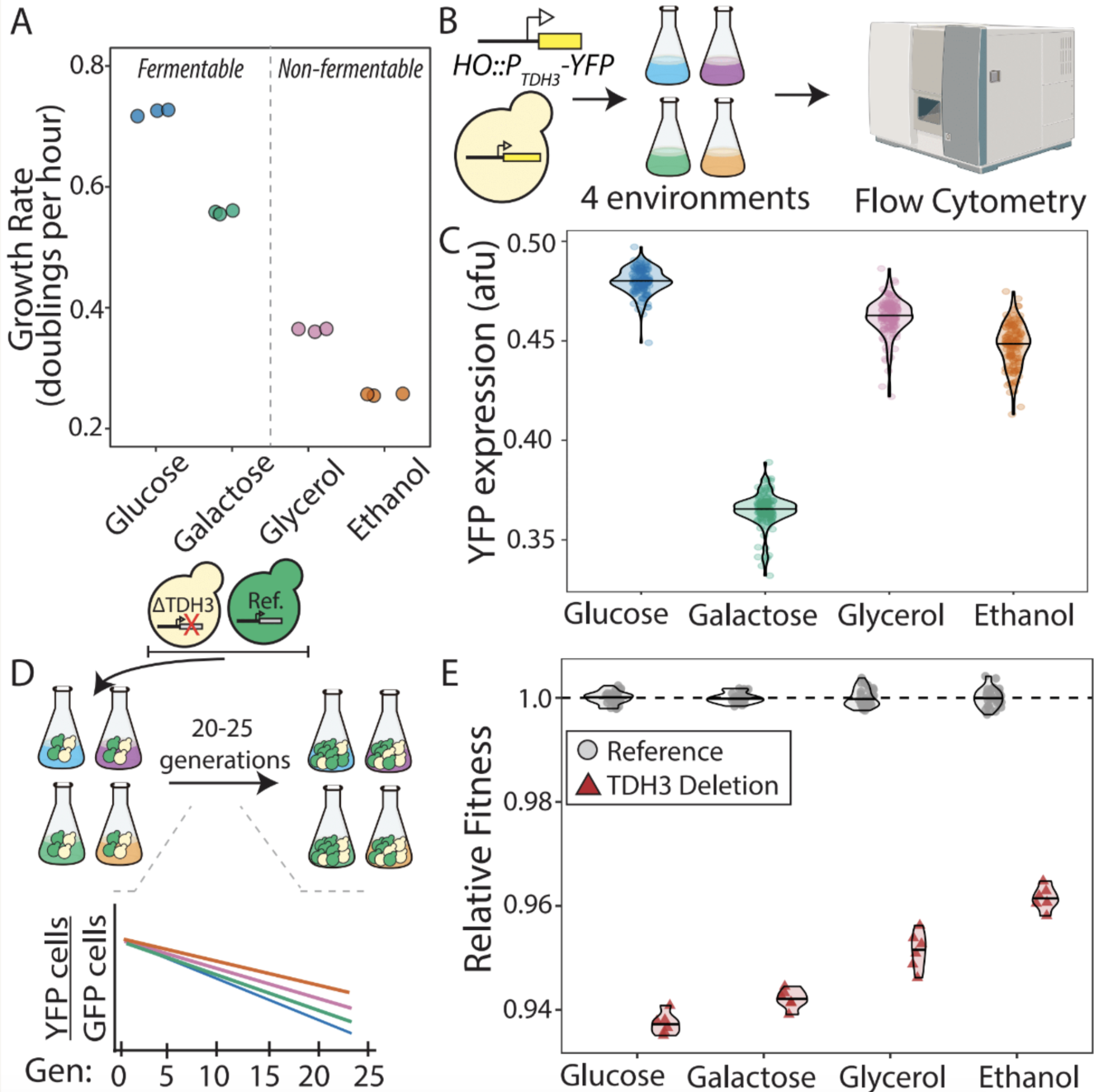
Environmental impacts on growth rate, *TDH3* expression and fitness effects of *TDH3*. (A) The average number of cell divisions per hour (growth rate) is shown for a wildtype strain of *S. cerevisiae* grown in media containing glucose, galactose, glycerol, or ethanol as a carbon source. Growth rate was calculated from optical density measurements of batch cultures over time. As indicated, glucose and galactose are fermentable carbon sources; glycerol and ethanol are non-fermentable carbon sources. (B) Schematic shows that to measure activity of the *TDH3* promoter (*P_TDH3_*) in different environments, a *P_TDH3_-YFP* reporter gene was integrated into the HO locus, cells were grown in each environment (media containing glucose, galactose, glycerol, or ethanol as a carbon source), and YFP fluorescence was measured using flow cytometry. (C) Violin plots show arbitrary fluorescence units (afu) for YFP (corrected for cell size) driven by the *TDH3* promoter in the four different environments. In each case, the horizontal line shows the median expression in each environment and the points show individual replicates. (D) To quantify relative fitness of a wildtype reference strain (not shown) or a *TDH3* deletion strain (ΔTDH3), each marked with YFP, relative to a reference competitor genotype (Ref.) marked with GFP, we mixed cells equally from the two competing genotypes and tracked the relative abundance of each genotype over 20-25 generations of log-phase growth (maintained by diluting the culture with new media every 12 hours) using flow cytometry every ∼6-8 generations. (E) Violin plots show fitness of the reference strain (grey circles) and *TDH3* deletion (red triangles) relative to the GFP-positive competitor strain in the different environments. For each environment, points show the relative fitness of individual replicates (4-6 replicates per condition and genotype) and horizontal lines show the median relative fitness for each genotype.

To determine how these different environments impacted expression of the *TDH3* gene and its contribution to the rate of cell division, we measured the activity of the *TDH3* promoter allele in the same genetic background in each environment and then determined how the loss of *TDH3* impacted the rate of cell division. Activity of the *TDH3* promoter was quantified by using flow cytometry to measure the fluorescence of individual cells with the reference allele of the *TDH3* promoter (*P_TDH3_*) driving expression of a yellow fluorescent protein (YFP) at the HO locus in each of the four environments (Fig. 2*B*). We found that the activity of the *TDH3* promoter varied among environments, with the highest activity in glucose and the lowest activity in galactose (Fig. 2*C*). We then examined the contribution of *TDH3* to population growth in each environment by using flow cytometry to compare the relative growth rates of strains with and without a functional copy of the *TDH3* gene. The reference and deletion strains were marked with GFP (green fluorescent protein) and YFP reporter genes, respectively (Fig. 2*D*). We found that the contribution of *TDH3* to growth rate varied among environments (Fig 2*E*).

Interestingly, the expression and fitness data suggested a complex relationship between the expression level of *TDH3* in an environment and the importance of *TDH3* for growth in that environment (Fig. 2*C* and 2*E*). For example, *TDH3* had the highest expression and largest fitness impact upon deletion in glucose, but the lowest expression level and the second largest fitness impact upon deletion in galactose. This extreme case scenario–a comparison of a reference and deletion strain–demonstrates that even when a mutation has identical effects on expression in different environments (a complete loss of expression in this case), it can have different effects on relative fitness.

### Environments impact the relationship between TDH3 expression and fitness

To better understand the relationship between *TDH3* expression and its effects on growth in different environments, we identified 47 alleles of the *TDH3* promoter that showed a wide range of effects on *P_TDH3_* activity in glucose (35–37) and measured their effects on expression of the YFP reporter gene inserted at the *HO* locus in each of the three other environments. We then used each of these mutant alleles to drive expression of the native *TDH3* gene and measured its effects on relative growth rate in all four environments via competitive growth assays, as described above. For all *TDH3* promoter alleles, point mutations were located in RAP1 or GCR1 transcription factor binding sites and/or in the TATA box (*SI Appendix* Fig. S2*A*). The promoter alleles examined included 12 alleles with 1 mutation, 11 alleles with 2 mutations, and 11 alleles with more than 2 mutations relative to the reference sequence (*SI Appendix* Fig. S2*B*). The remaining 13 mutant genotypes included a duplication of the *P_TDH3_*-YFP reporter gene (when measuring expression) or of the full *TDH3* gene (when measuring fitness) with or without promoter mutations in both copies of the gene (*SI Appendix* Fig. S3).

Focusing first on *P_TDH3_* activity, we compared YFP expression driven by each of the mutant promoter alleles to expression driven by the unmutated reference allele in the same environment. We found that the overall range of expression levels observed in this set of 47 *P_TDH3_* alleles was similar in all four environments: 0% to 207% in glucose, 0% to 185% in galactose, 1% to 186% in glycerol, and 1% to 182% in ethanol (*SI Appendix* Table S1). As expected, alleles carrying 1 promoter mutation had smaller effects on expression than alleles carrying multiple mutations, and alleles with duplications of the *TDH3* promoter often had expression levels higher than the unmutated reference allele (Fig. 3*A*).

**Figure 3:**
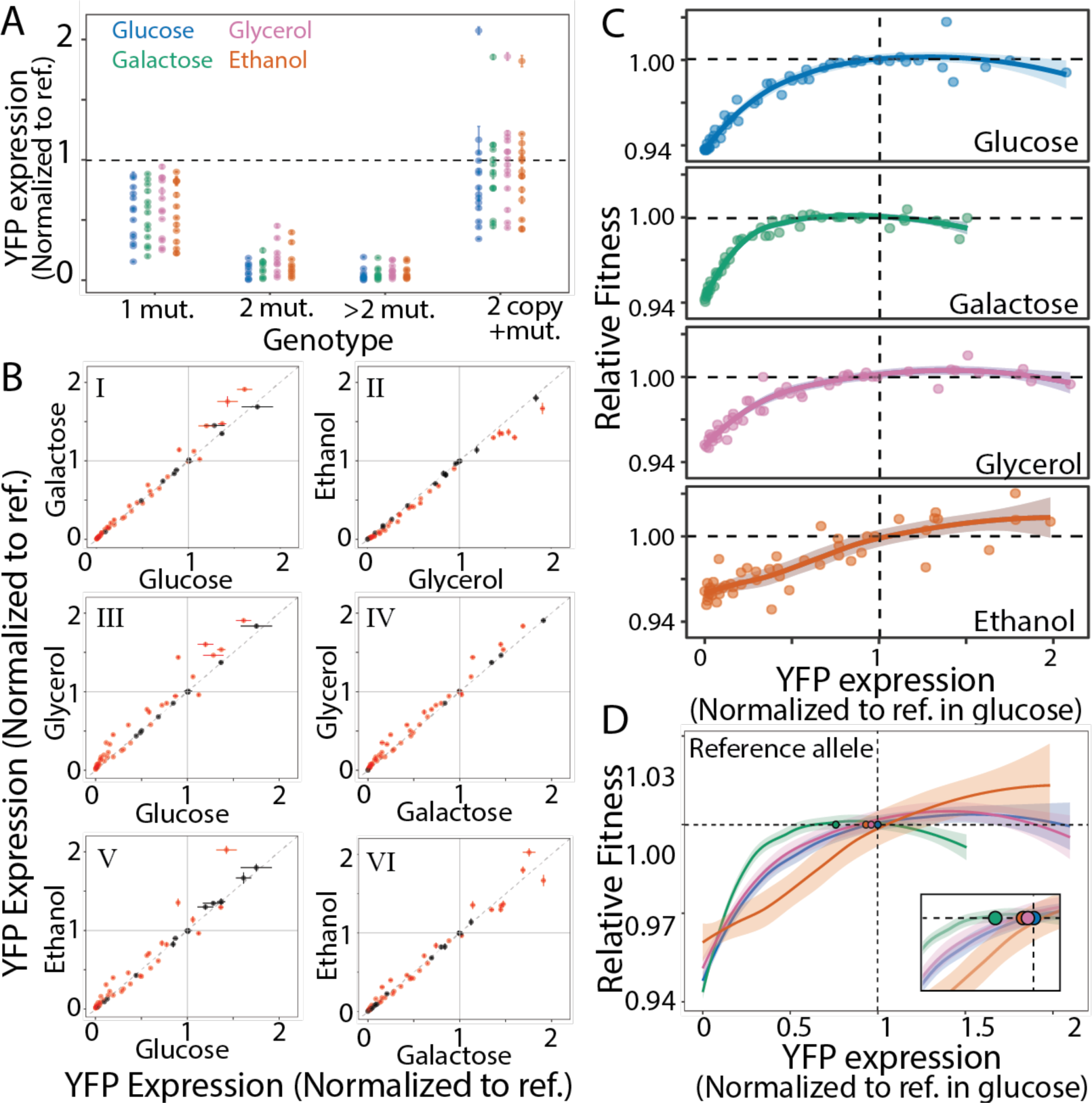
Environment-specific effects of *P_TDH3_* mutations on gene expression and fitness. (A) Mean YFP expression (with error bars representing 95% confidence intervals for 4-6 replicates of each genotype in each environment) is shown for each of the 47 mutant *P_TDH3_* alleles driving expression of YFP in glucose (blue), galactose (green), glycerol (purple), and ethanol (orange), with mutants grouped by whether they have 1 mutation (1 mut.), 2 mutations (2 mut.), more than two mutations (>2 mut.), or two copies of *P_TDH3_* with mutations (2 copy +mut.). YFP expression of each strain was normalized to the expression level of the strain carrying an unmutated *TDH3* promoter grown in glucose. (B) Pairwise comparisons for effects of mutant *P_TDH3_* alleles on gene expression in glucose, galactose, glycerol, and ethanol. The mean expression level of each of the 47 *P_TDH3_*-YFP mutant genotypes relative to the reference strain is shown with error bars indicating the standard deviation among replicates. Panels I and II show comparisons in the two fermentable (glucose and galactose) and two non-fermentable (glycerol and ethanol) environments, respectively. Panels III-VI show comparisons between fermentable and non-fermentable environment Genotypes with significantly different effects between the environments compared are shown in red. (C) Relative activity of the *P_TDH3_* promoter (measured using YFP expression and normalized to expression of the reference in glucose) is plotted against the relative fitness of the strain carrying the same *P_TDH3_* allele at the native locus, with fitness normalized to the fitness of the unmutated reference strain in each environment. These points were used to estimate the shape of the fitness function using LOESS regression. The lighter shaded areas surrounding the fitness function in each environment indicates its 95% confidence interval. (D) The fitness functions shown in C are overlaid to show similarities and differences between them. The colored dots represent relative expression and fitness of the unmutated *P_TDH3_* reference allele in glucose (blue), galactose (green), glycerol (purple), and ethanol (orange). Inset shows a higher magnification of the region of relative expression from approximately 0.5 to 1.1 to better show expression plasticity of the reference allele.

To determine whether individual mutant alleles had environment-specific effects on expression (i.e., gene-by-environment interactions), we used a linear model to separately estimate the effects of the mutant genotype, environment, and genotype-by-environment interactions on gene expression. Despite strong overall correlations between the effects of *P_TDH3_* alleles among environments, we found significant gene-by-environment interactions in many cases (Fig. 3*B*); 45 of the 47 genotypes had effects that were significantly different between at least two environments. Overall, mutations generally had more similar effects on expression in the two fermentable or two non-fermentable carbon sources than between fermentable and non-fermentable carbon sources (Fig. 3*B*). This observation is consistent with a prior study of 235 *P_TDH3_* alleles carrying single point mutations and polymorphisms analyzed in glucose, galactose, and glycerol (35).

To examine whether and how the relationship between *TDH3* expression and fitness varies among environments, we plotted the effects of each mutant allele on gene expression and fitness in each environment. We observed non-linear relationships in all environments with differences in both the optimal expression level (i.e. expression level that maximizes fitness) and the shape of the fitness function among environments (Fig. 3*C*). We expected that the fitness functions for *TDH3* expression might look most similar for cells grown on the same type of carbon source (fermentable or non-fermentable) because they would be using *TDH3* in more similar metabolic processes, but we found that this was not the case. For example, moderate reductions in *TDH3* expression caused significant reductions in fitness in glucose but not galactose, and increasing *TDH3* expression was much more beneficial in ethanol than glycerol (Fig. 3*C*). Surprisingly, the greatest similarity in the shape of the fitness functions was seen for glucose (a fermentable carbon source) and glycerol (a non-fermentable carbon source) (Fig. 3*C*).

Given that the optimal *TDH3* expression level varied among environments (Fig. 3*C*) and that the environment also affected expression driven by the reference *P_TDH3_* allele (Fig. 2*A*), we next asked whether this plasticity served to increase the fitness of the reference allele. Between glucose and galactose, we found that the unmutated reference *P_TDH3_* allele exhibited plasticity that resulted in 24% lower expression in galactose than glucose but fitness remained near the optimum in both environments because of differences in the shape of the fitness functions (Fig. 3*D*). This observation suggests that this plasticity is either neutral or beneficial. By contrast, the unmutated reference allele exhibited 7% lower expression in ethanol than in glucose even though higher expression levels were associated with higher fitness in this environment. The fitness function observed in ethanol was distinct from the fitness functions in the other three environments in that it did not reach a fitness peak or plateau within the range of *TDH3* expression levels sampled (Fig. 3*C*). This unique relationship between *TDH3* expression and fitness is consistent with *S. cerevisiae* using ethanol quite differently from the other three carbon sources (Fig. 1*E*): glucose, galactose, and glycerol are precursors of GAP (the substrate that *TDH3* converts into 1,3 biphosphoglycerate in the second part of the glycolysis metabolic pathway) whereas ethanol is not a precursor of GAP and the role of TDH3 in ethanol metabolism likely involves gluconeogenesis.

### Molecular mechanisms underlying environment-specific effects on expression and fitness

In any environment, the fitness function determines whether a mutation’s effect is visible to selection; selection is agnostic to the molecular mechanisms through which that mutation affects the phenotype. However, mutations affecting some molecular mechanisms may be more likely to be exposed to or hidden from selection because they have environment-specific effects. For example, promoters regulate gene expression by binding to context-specific transcription factors as well as core transcription factors that are part of the basal transcriptional machinery required every time a gene is transcribed. Mutations altering these different types of binding sites are expected to differ in their environmental sensitivity to effects on gene expression and fitness, which could preferentially maintain some promoter mutations in populations when environments are changing. For example, mutations in binding sites for context-specific transcription factors that only function in one environment might not affect expression in other environments and thus be invisible to selection when the organism is in these other environments. By contrast, mutations that alter regulatory sequences used in all environments might always be subject to selection.

The *S. cerevisiae TDH3* promoter contains a binding site for the RAP1 transcription factor, two binding sites for the GCR1 transcription factor, and a TATA-box sequence upon which the basal transcriptional machinery assembles (Fig. 4*A*). Because prior work suggests the RAP1 and GCR1 binding sites play different roles in regulating *TDH3* expression in fermentable and non-fermentable environments (35, 38), we hypothesized that mutations affecting the RAP1 and GCR1 binding sites would be more prone to environment-specific effects than mutations affecting the TATA box, which functions more similarly in all environments. To test this hypothesis, we examined the 7 mutant genotypes in our datasets with a single copy of *P_TDH3_* that carried a single point mutation in one of the RAP1 or GCR1 binding sites and compared their effects on gene expression and fitness to the 7 mutant genotypes in our dataset with one or more mutation(s) in the TATA box (*SI Appendix* Fig. S2*B;* highlighted with asterisk). Environmental variability in expression and fitness was estimated for each of the 14 mutant genotypes by calculating the variance of the relative *P_TDH3_-YFP* expression or relative fitness measures for each mutant allele among the four environments.

**Figure 4.**
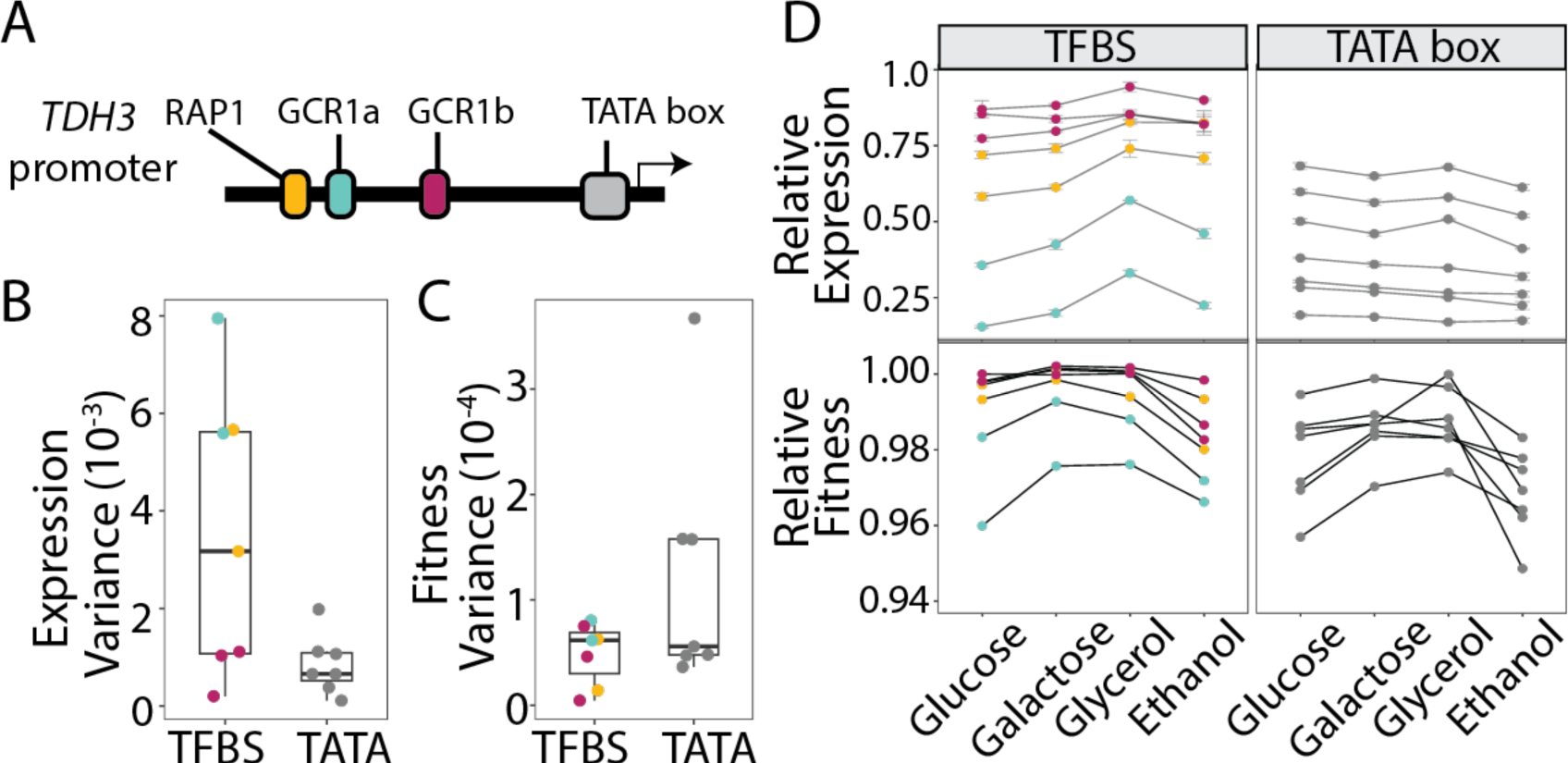
Effects of mutations in RAP1 and GCR1 binding sites have greater environmental variance than TATA box mutations for gene expression, but not fitness. (A) Schematic of the *S. cerevisiae TDH3* promoter shows the location of binding sites for the transcription factors RAP1 (yellow) and GCR1 (blue and red) as well as the location of the TATA box (grey) and transcription start site (arrow) for *TDH3*. (B) Variance in effects of mutations on gene expression in the four environments tested (normalized to the reference allele in each environment) is shown for *P_TDH3_* alleles with a mutation only in either a RAP1 or GCR1 binding site (TFBS) or with a mutation(s) only in the TATA box (TATA). In each box plot, the center line indicates the median effect size for the group. (C) The same information is shown as in panel B but for variance in the effects of the promoter alleles on fitness rather than expression in the four environments. (D) Effects of each of the 14 *P_TDH3_* mutant alleles shown in panels B and C on gene expression and fitness are shown in each of the four environments. Error bars indicate the 95% confidence interval for each genotype in each environment, calculated from 6-9 replicates for gene expression and 4 replicates for fitness. In all panels, the color of each point indicates whether the strain carried a mutation(s) in the RAP1 binding site (yellow), GCR1 binding site (blue or red) or the TATA box (grey).

We observed the hypothesized pattern for the effects of mutations on gene expression, but not fitness. The measures of expression variability between the set of alleles with mutations in either a RAP1 or GCR1 binding site tended to have effects on gene expression that were more variable than the set of alleles with a mutation(s) in the TATA box (directional Mann-Whitney U test, *p = 0.048*, Fig. 4*B*), with mutations that had the most variable effects on expression among environments located in the RAP1 binding site and the adjacent GCR1 binding site (GCR1a in Fig. 4*A*). Despite the more environmentally variable effects of mutations in the RAP1 and GCR1 binding sites on gene expression (Fig. 4*B*), the 3 mutant alleles with the most environmentally variable effects on fitness all carried a mutation(s) located in the TATA box (Fig. 4*C*). Overall, the average variance of the 7 TATA box mutants for fitness was higher than the average variance of the 7 RAP1/GCR1 binding site mutants (Fig. 4*C*), although this difference was not statistically significant (directional Mann-Whitney U test, *p = 0.19*). Looking at the expression and fitness of each of these mutants individually in the four environments showed that the effects of these mutations on expression were generally largest in glucose and galactose and the effects of TATA box mutations on fitness were often largest in ethanol (Fig. 4*D*), which has a steep fitness function (Fig. 3*D*). The differences in effects on expression and fitness we observed for the two classes of mutations are explained by their plasticity effects as well as the different shapes of the fitness functions among environments.

## Discussion

By measuring the effects of promotor mutations on both *TDH3* expression and fitness in four different carbon environments, this study illustrates the dual role of the environment in the production and selection of phenotypic variation. The plasticity in promoter activity, mutations exhibiting environment-dependent effects, and differences in the shape of the fitness functions among environments that we observed are all expected to influence the evolutionary fate of genetic variants in the changing environments yeast experience in the wild. For example, when *S. cerevisiae* grows on a fermentable carbon source such as glucose, the non-fermentable carbon source ethanol accumulates and, when glucose is exhausted, the ethanol produced is metabolized via respiration. The differences in fitness functions we observed among environments with different carbon sources can favor promoter alleles that exhibit plasticity conferring higher fitness in multiple environments. Indeed, a prior study comparing the effects of mutations and polymorphisms on *P_TDH3_* activity suggested that natural selection has favored a particular degree of *TDH3* expression plasticity between glucose and galactose (35), although the differences in fitness functions reported here for these two environments suggest that even greater plasticity (in the same direction) could be even more beneficial.

When considering how mutations in the *TDH3* promoter impacted expression and fitness, we found that mutations in the RAP1 and GCR1 binding sites caused greater variability in gene expression among the four environments tested. Yet, these mutations were no more variable in their effects on fitness than mutations in the TATA box. In fact, the three strains with the most variable effects on fitness had a mutation(s) in the TATA box. This observation suggests that genetic variation in the RAP1 and GCR1 transcription factor binding sites is less likely to be deleterious when environments are changing than genetic variation in the TATA box, which could explain why genetic variants causing differences in gene expression between strains of *S. cerevisiae* are often found in transcription factor binding sites (34) and show gene-by-environment interactions (32). For example, genetic variation in the RPN4 binding site of the *ERG4* promoter and the GAL4 binding site of the *PGM1* promoter have both been shown to have adaptive, environment-specific effects in *S. cerevisiae* and related species (32, 44–46).

Arguments that condition-specific effects of *cis*-regulatory mutations in transcription factor binding sites are less deleterious than mutations affecting a gene’s expression constitutively have been made most often for tissue-specific enhancers found in metazoans (39–43) but the same logic applies to environment-specific regulatory sequences in yeast promoters.

The different shapes of fitness functions we observed for *TDH3* expression among environments also suggest a way for populations of *S. cerevisiae* to accumulate genetic variation that is inconsequential for fitness in one environment but affects fitness in another. Such genetic variation is often referred to as “cryptic” because its effects are only seen in some environments (28,47). For example, our data suggest that mutations with small to moderate effects on *TDH3* expression (e.g., up to a 50% decrease in expression relative to wildtype) could accumulate neutrally in a population growing on galactose as a carbon source but would then be selected against if the population shifted to growing on glucose, glycerol, or ethanol (Fig. 3*D*). If the effects of these mutations were plastic, however, such that they caused different changes in *TDH3* expression in different environments, they could be neutral in multiple environments, as we saw for the reference allele in glucose, galactose, and glycerol (Fig. 3*D*). With limited data available showing the joint effects of the environment on the production of phenotypic variation and the shape of fitness functions, it remains to be seen how widespread these phenomena might be.

Understanding how the environment influences phenotypes and fitness is a shared goal of evolutionary and molecular genetics, yet studies documenting phenotypic plasticity and gene-by-environment interactions are too rarely integrated with studies investigating environmental impacts on cellular processes. By considering the regulatory and fitness effects of individual mutations on *TDH3* expression in the context of what is known about the molecular mechanism controlling *TDH3* promoter activity, this study provides one such bridge connecting these fields and suggests that historical and ongoing changes in environments might result in selection for phenotypic plasticity that favors particular types of evolutionary changes in regulatory sequences. This type of functional synthesis between molecular and evolutionary biology is critical for addressing fundamental questions in both fields (50).

## Materials and Methods

### Yeast strains

All *S. cerevisiae* strains used in this work to measure the effects of *P_TDH3_* mutations on gene expression and fitness (*SI Appendix* Figure *2* and Table *S1*) were haploids constructed as described in (36, 37, 51). Briefly, to assess the impact of genetic variants in the *TDH3* promoter on its activity, mutant alleles of *P_TDH3_* were cloned upstream of the coding sequence for the venus YFP followed by a *cyc1* terminator, with this reporter gene integrated into the genome at the *HO* locus (*ho::P_TDH3_-YFP-T_CYC_*). To measure the effects of these promoter alleles on fitness, a parallel set of strains was constructed in which the native *P_TDH3_* promoter was replaced by each of the mutant *P_TDH3_* alleles. Prior work (37) has shown that the effects of mutant *P_TDH3_* alleles on reporter gene expression at the *HO* locus is strongly correlated (R^2^ = 0.99) with the effects of the same alleles on *TDH3* expression at the native *TDH3* locus (as measured using a TDH3::YFP fusion protein). For the strains of yeast with two copies of the *P_TDH3_-YFP* reporter gene at the *HO* locus or two copies of *TDH3* at the native locus, a copy of the *URA3* gene was inserted between the two copies to minimize cross-talk between them. Separate reference strains were constructed for the analysis of these strains containing a copy of *URA3* following the *P_TDH3_-YFP* reporter gene at the HO locus or after the native *TDH3* gene (36). The BY4741-drived background used for all strains contained genetic changes that increased sporulation efficiency and decreased petite frequency relative to the alleles of the laboratory S288c strain, as described in (51). The reference *P_TDH3_* allele carried by this strain differs from the S288c allele by an A to G substitution 293 bp upstream of the start codon that has little effect on expression or fitness (*SI Appendix* Table *S1*).

### Environments tested

To compare the growth of *S. cerevisiae* strains on four different carbon sources, cells were cultured in four types of media that were identical except for the carbon source. Each environment contained 10 g/l of yeast extract, 20 g/l of peptone along with either 20 g/l of glucose (YPD), 20 g/l of galactose (YPGal), 30 ml/l of 99% glycerol (YPG) or 50 ml/l of 99% ethanol (YPE).

### Measuring growth rate in four environments

To determine the growth rate of *S. cerevisiae* (strain YPW1160, which was also used as the competitor strain in the fitness assays described below), we measured its doubling time when grown in YPD (glucose), YPGal (galactose), YPG (glycerol) and YPE (ethanol). Three replicate cultures of YPW1160 were started in parallel in 5 ml of YPD, YPGal, YPG or YPE and incubated for 36 hours at 30°C with dilution to 5 × 10^5^ cells/ml every 12 hours. After the last dilution, cell density was quantified every 60 minutes for 10 hours and then after another 800 minutes by measuring optical density at 660 nm. Doubling time was calculated as the inverse of the slope of the linear regression of log(cell density)/log(2) on time during the logarithmic of growth where the relationship between log(cell density) and time is linear. The average doubling time was found to be 80 minutes in YPD, 108 minutes in YPGal, 165 minutes in YPG, and 233 minutes in YPE. These data were converted to the doublings per hour shown in Fig. *2A* by dividing 60 by each of these values.

### Measuring effects of P_TDH3_ alleles on gene expression

Fluorescence attributable to expression of the *P_TDH3_-YFP* reporter gene was measured as a proxy for *P_TDH3_* transcriptional activity using flow cytometry as previously described (51). Briefly, all strains were revived from -80°C glycerol stocks on YPG plates (10 g Yeast extract, 20 g Peptone, 30 ml Glycerol, 20 g agar per liter) and, after 2 days of growth, were arrayed in 96-well plates containing 0.5 ml YPD per well. In addition to the different samples, a reference strain YPW1002 was inoculated in 24 positions, which was used to correct for plate and position effects on fluorescence. Strain YPW978, which does not contain the YFP reporter construct (51), was inoculated in one well per plate and served to correct for autofluorescence. Cells were maintained in suspension by fitting the culture plates on a rotator. After the 22 hours of growth in YPD, samples were diluted to one of four different environments that differed only by the carbon source. Expression assays in glucose and galactose were performed in parallel but on a different day than the experiment in glycerol and ethanol. Samples were acclimated to each environment by dilutions to fresh medium every 12 hours for 36 hours. Prior to each dilution, cell density was measured for all samples using a Sunrise absorbance reader (Tecan), and one dilution factor was calculated for each 96-well plate so that the average cell density would reach 5 × 10^6^ cells/ml after 12 hours of growth. This procedure ensured that all samples were maintained in constant log-phase after the first few hours of growth, because no sample reached a density above 10^7^ cells/ml, while limiting the strength of genetic drift because the smallest number of cells transferred during dilution was ∼10,000. After 36 hours of growth, samples were diluted to 2.5 × 10^6^ cells/ml in PBS, and fluorescence was acquired for 20,000 events per well on the BD Accuri C6 flow cytometer coupled to a HyperCyt autosampler (IntelliCyt Corp). A 488-nm laser was used for excitation and a 530/30 optical filter was used for acquisition of the YFP signal.

Flow cytometry data were analyzed using R packages *flowCore* and *flowClust* as described in (37). Briefly, single cells were separated from all events based on the height and the area of the forward scatter signal. Then, the intensity of the fluorescence signal was normalized by cell size in several steps (37), Extended methods), and the YFP signal was adjusted for autofluorescence based on the signal from strain YPW978, which lacked YFP. The median expression level for each replicate of each genotype was calculated (6 replicates per sample, 24-30 replicates of control strains without promoter mutations), and expression of each genotype was estimated as the mean of the median values. The plasticity of each mutant promoter variant was determined by dividing YFP expression level of each mutant genotype in each environment by the YFP expression level of the unmutated reference strain (YPW1002 for strains with one YFP copy; strain YPW2675 for strains with two YFP copies) in that environment. The main effect of the mutations, environment, and gene-by-environment interactions were estimated using a linear model in R, with terms for genotype, environment, and a gene-by-environment interaction. Scripts used for this analysis are included in *SI Appendix* File 1.

### Fitness assays

We used head-to-head competition assays between strains expressing different levels of *TDH3* protein and a common reference to measure relative growth rate during log-phase and used this as our proxy for fitness as described in (37). Briefly, all strains with promoter mutations at the native *TDH3* locus were engineered to also have an unmutated *P_TDH3_-YFP* reporter gene at the *HO* locus, allowing them to be recognized by their yellow fluorescence. A common competitor strain (YPW1160) with no mutations at the native *TDH3* locus was engineered with a *P_TDH3_-GFP* allele at the *HO* locus that was recognized by its green fluorescence. The 47 YFP-marked mutant strains carrying different variants of the *TDH3* promoter at the native locus (*SI Appendix* Table *S1*) were arrayed on four 96-well plates, with two replicates of each strain on each plate. In parallel, the common competitor YPW1160 expressing GFP was also arrayed on four 96-well plates. After 24 hours of growth in 0.5 ml of YPD at 30°C, 0.1 ml of each culture was mixed with 23 μl of 80% glycerol in 96 well plates and stored at -80°C. From each of the eight culture plates, four plates were frozen to be used for the competition assays performed in the four different environments (YPD, YPGal, YPG and YPE). Competition assays were performed consecutively in the four environments over three weeks, starting with ethanol, then glucose, glycerol, and galactose. The same batches of media were used for the expression and competition experiments, and cells were grown in 96-well plates at 30°C and maintained in suspension on a rotor for all assays. A similar protocol was used for all environments, except for differences in the timing of dilutions mentioned below to adjust for variation in doubling time.

For each competitive growth experiment, YFP and GFP strains were thawed on eight separate omnitrays filled with YPG agar medium. After 48 hours of incubation at 30°C, samples were transferred in 0.5 ml of YPD and grown for 24 hours to saturation. Then, cell densities were measured using the Sunrise absorbance reader. At this point, equal volumes of YFP and GFP cell cultures were mixed in 0.5 ml of either YPD, YPGal, YPG or YPE medium in four 96-well plates. The dilution factor was calculated for each plate based on the doubling time of the GFP strain so that the average cell density would reach ∼5 × 10^6^ cells/ml after 12 hours of growth. This procedure of cell density measurement and dilution followed by 12 hours of growth was repeated three times and constituted the acclimation phase of the experiment, during which the relative frequency of YFP and GFP strains was not recorded. After these first 36 hours of growth, competitive growth was carried on for three additional cycles of dilution and growth during which the ratio of YFP and GFP cells was recorded at four time points (before each cycle of growth and at the end of the last cycle). The duration of these three cycles of competitive growth depended on the environment: samples were diluted every 10 hours in YPD, every 12 hours in YPGal, and every 24 hours in YPG and YPE. Cell density was measured for all samples before each dilution. The relative frequency of YFP and GFP cells was quantified at each time point by flow cytometry. To this end, samples from incubated plates (not from freshly diluted cultures) were diluted in 0.3 ml of PBS to a final density of 1.5 × 10^6^ cells/ml in four 96-well plates and placed on ice to stop growth. These flow cytometry plates were prepared either immediately after samples were diluted to fresh medium for growth (first three time points) or at the end of the last cycle of growth (last time point). Approximately 75,000 events were recorded for each sample on the BD Accuri C6 flow cytometer, using a 488 nm laser for excitation and two different optical filters (510/10 and 585/40) to acquire fluorescence. These filters allowed the separation of the GFP and YFP signals. Therefore, the relative frequency of YFP and GFP cells was measured at four time points during the competition assays.

The number of cells expressing either YFP or GFP was counted for each sample using custom R scripts, as described in Duveau et al (37). Briefly, for each sample, we determined the number of cell generations that occurred during the three dilution cycles, with the median number of generations for all samples grown on the same 96-well plate used as a robust estimator of the number of generations for all samples on that plate. The number of generations over the entire experiment was found to be about 22 in YPD (glucose), 20 in YPGal (galactose), 26 in YPG (glycerol), and 13 in YPE (ethanol). The fitness of the YFP strain relative to the GFP competitor was calculated as the exponential of the slope of the ratio of YFP-positive over GFP-positive cells regressed on the number of generations across the four time points. For each *P_TDH3_* variant, replicates for which fitness departed from the median fitness across all eight replicates by more than four times the median absolute deviation were considered outliers and were excluded from further analysis. For each sample, the fitness relative to the GFP strain was then divided by the mean fitness for all replicates of the reference strain YPW1189 (for single-copy *P_TDH3_* variants) or YPW2682 (for double-copy *P_TDH3_* variants). We then calculated the mean relative fitness and standard deviation over the eight replicates of each variant. This measure of fitness expressed relative to a strain with the reference *P_TDH3_* allele was used in all subsequent analyses.

### Additional data analysis

All data analysis was done using custom R code included as *SI Appendix* File 1. This file includes code for the LOESS regression used to describe the relationship between median *TDH3* expression and fitness from the data collected for all *P_TDH3_* variants, which was performed using the R function *loess* with a span of 0.75 with the reference allele and null alleles of the *TDH3* promoter assigned weights of 100. It also includes code for the statistical tests used to compare the effects of mutations in the RAP1 and GCR1 binding site to the effects of mutations in the TATA box as well as the linear models used to test for gene-by-environment interactions. Code used for plotting data and results from statistical analyses is also included in this file.

## Acknowledgments

We thank Lisa Kim for help determining single-cell division rates, Andrea Hodgins-Davis and Brian PH Metzger for helpful discussions, and all members of the Wittkopp lab for helpful comments on the manuscript. This work was supported by a European Molecular Biology Organization postdoctoral fellowship (EMBO ALTF 1114–2012) to FD, National Institutes of Health National Research Service Award (5F32CA261115) to MAS, and grants from the National Science Foundation (MCB-1021398) and National Institutes of Health (R01GM108826 and 5R35GM118073) to PJW.

---

**SI Appendix Figure S1.**
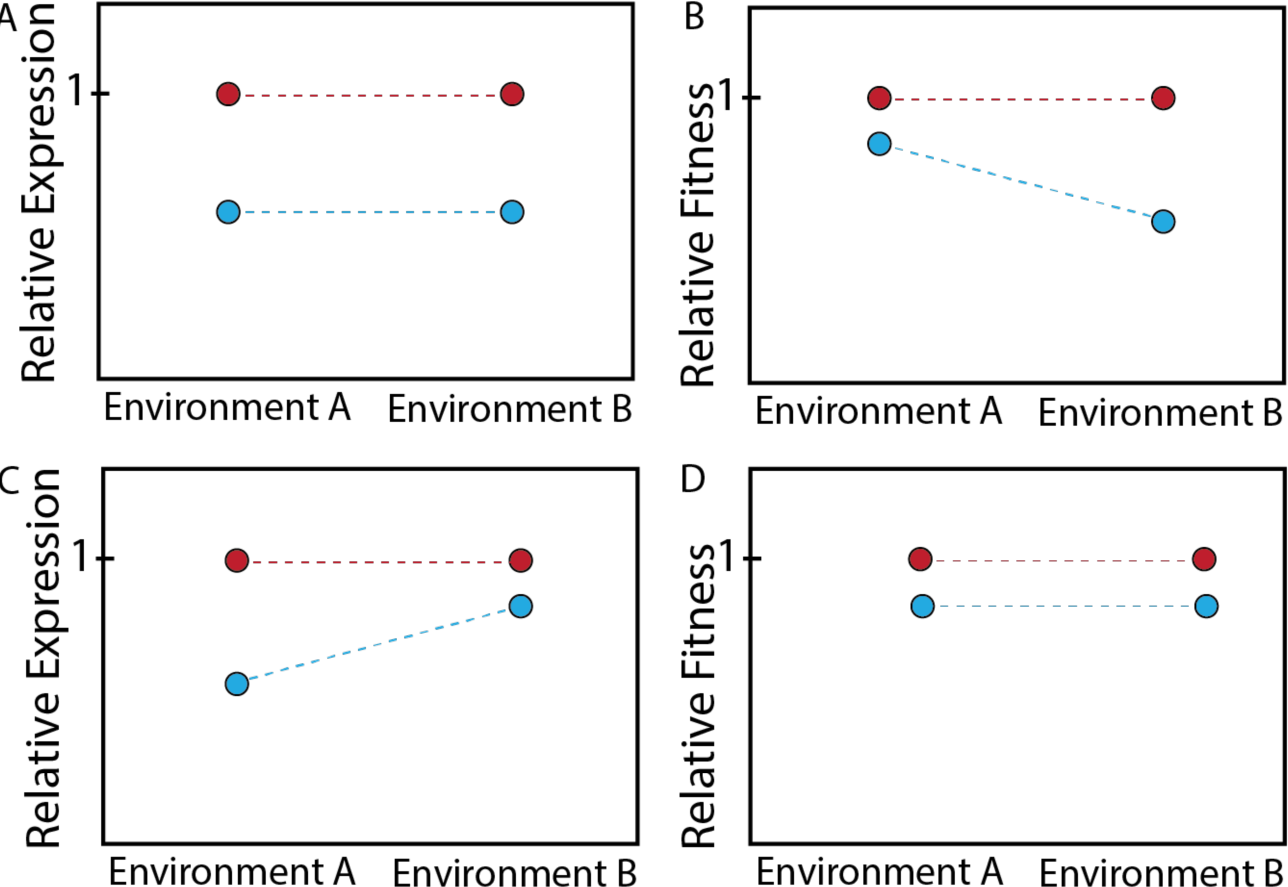
Examples of gene-by-environment interactions for gene expression and fitness. (A, B) Relative gene expression (A) or relative fitness (B) is shown for the scenario depicted in Fig. 1C, with values for the wild-type allele shown in red and values for the alternate allele shown in blue. (C, D) Relative gene expression (C) or relative fitness (D) is shown for the scenario depicted in Fig. 1D, with values for the wild-type allele shown in blue and values for the alternate allele shown in red. In each of the four plots (A-D), lines that are not parallel indicate a gene-by-environment interaction.

**SI Appendix Figure S2.**
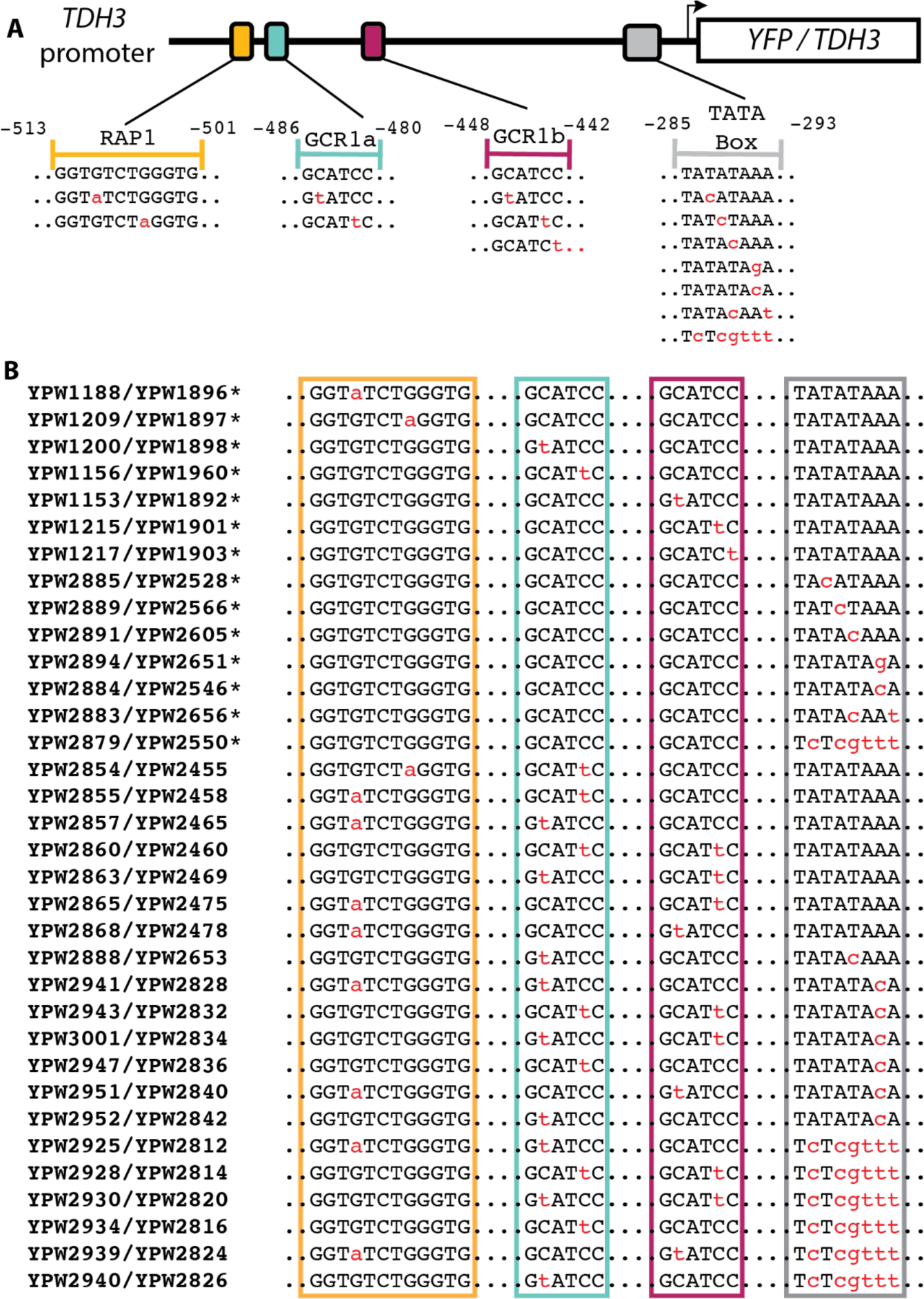
Mutations in strains carrying a single mutant copy of the *TDH3* promoter. (A) Schematic shows the reference sequence and mutations tested in the RAP1 binding site (yellow), two GCR1 binding sites (blue, red), and the TATA box (grey) of the *TDH3* promoter. Coordinates shown are the number of bases upstream of the start codon of the downstream *YFP* or *TDH3* protein. (B) The mutation(s) present in each of the 34 mutant *P_TDH3_* alleles assayed with a single copy of the *P_TDH3_-YFP* reporter gene are shown. In each case, the first strain listed (YPW####) carries the mutant *TDH3* promoter allele cloned upstream of a YFP reporter protein at the *HO* locus and was used to measure the effects of the mutation(s) on gene expression whereas the second strain listed carries the mutant *P_TDH3_* allele at the native *TDH3* locus and was used to measure the effects of the mutation(s) on fitness (*SI Appendix* Table *1*). Asterisks show alleles that were used to compare environment-dependent effects of disrupting TATA box or the TFBS.

**SI Appendix Figure S3.**
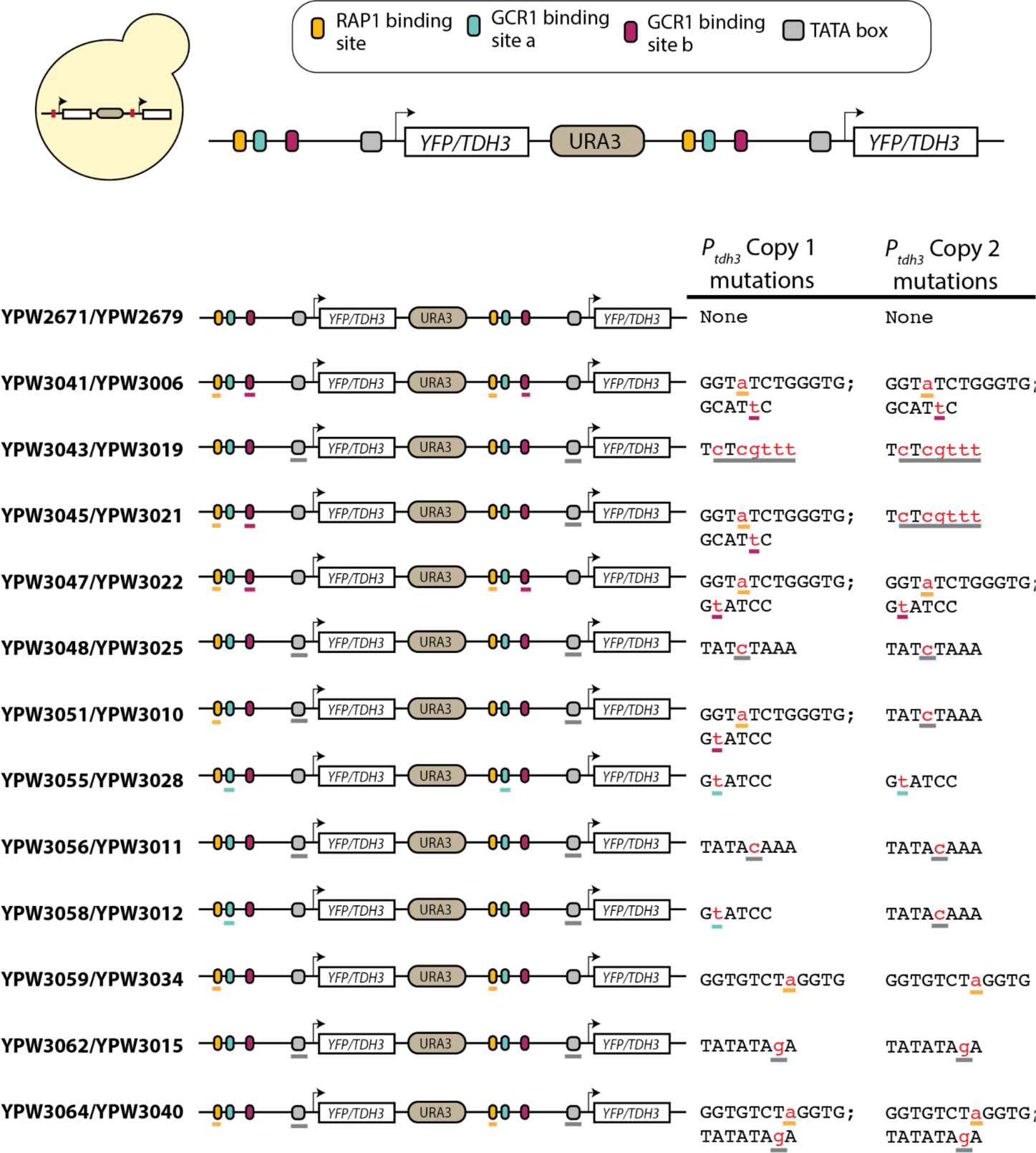
Mutations in strains carrying two mutant copies of the *TDH3* promoter. The schematic at the top of the figure shows the arrangement of two copies of the *TDH3* promoter driving expression of either a YFP reporter protein at the *HO* locus or the native *TDH3* protein at the native locus, with each gene pair separated by a copy of the *URA3* gene. The specific mutations carried in each of the 13 pairs of mutant strains carrying two copies of the *P_TDH3_* allele are then shown with the name (YPW####) of the strain carrying the reporter genes used to measure effects on gene expression listed before the name of the strain with the duplication of *TDH3* used to measure fitness (*SI Appendix* Table *1*). For each pair of strains, the mutated elements are underlined and the sequences of these elements are shown for each copy of the *TDH3* promoter with the mutated sites shown in red. The color of the underline corresponds to the feature mutated: RAP1 binding site = yellow, GCR1a binding site = blue, GCR1b binding site = red, and TATA box = grey.

**SI Appendix Table 1.** Summary of yeast strains, including mutation(s) carried, measures of YFP expression, and measures of relative fitness. (.xls file)

**SI Appendix File 1.** R code used for data processing and analysis.

## References

1. S. M. Scheiner, Genetics and evolution of phenotypic plasticity. Annu. Rev. Ecol. Syst. 24, 35–68 (1993).

2. A. D. Bradshaw, “Evolutionary Significance of Phenotypic Plasticity in Plants” in Advances in Genetics, E. W. Caspari, J. M. Thoday, Eds. (Academic Press, 1965), pp. 115–155.

3. M. J. West-Eberhard, Developmental Plasticity and Evolution (Oxford University Press, 2003).

4. C. K. Ghalambor, J. K. McKAY, S. P. Carroll, D. N. Reznick, Adaptive versus non-adaptive phenotypic plasticity and the potential for contemporary adaptation in new environments. Funct. Ecol. 21, 394–407 (2007).

5. S. J. Simpson, G. A. Sword, N. Lo, Polyphenism in insects. Curr. Biol. 21, R738–49 (2011).

6. E. B. Josephs, Determining the evolutionary forces shaping G × E. New Phytol. 219, 31–36 (2018).

7. A. B. Paaby, N. D. Testa, “Developmental plasticity and evolution” in *Evolutionary Developmental Biology*, (Springer International Publishing, 2018), pp. 1–14.

8. I. Goldstein, I. M. Ehrenreich, Genetic variation in phenotypic plasticity. Phenotypic Plasticity and Evolution 91 (2021).

9. A. A. Agrawal, C. Laforsch, R. Tollrian, Transgenerational induction of defences in animals and plants. Nature 401, 60–63 (1999).

10. D. Reznick, M. J. Butler Iv, H. Rodd, Life-history evolution in guppies. VII. The comparative ecology of high- and low-predation environments. Am. Nat. 157, 126–140 (2001).

11. F. Bashey, Cross-generational environmental effects and the evolution of offspring size in the Trinidadian guppy Poecilia reticulata. Evolution 60, 348–361 (2006).

12. S. C. Campbell-Staton, J. P. Velotta, K. M. Winchell, Selection on adaptive and maladaptive gene expression plasticity during thermal adaptation to urban heat islands. Nat. Commun. 12, 6195 (2021).

13. J. F. Storz, G. R. Scott, Phenotypic plasticity, genetic assimilation, and genetic compensation in hypoxia adaptation of high-altitude vertebrates. Comp. Biochem. Physiol. A Mol. Integr. Physiol. 253, 110865 (2021).

14. M. A. Taylor, et al., Large-effect flowering time mutations reveal conditionally adaptive paths through fitness landscapes in Arabidopsis thaliana. Proc. Natl. Acad. Sci. U. S. A. 116, 17890–17899 (2019).

15. W. Li, et al., A Natural Allele of a Transcription Factor in Rice Confers Broad-Spectrum Blast Resistance. Cell 170, 114–126.e15 (2017).

16. C. D. Kenkel, M. V. Matz, Gene expression plasticity as a mechanism of coral adaptation to a variable environment. Nat Ecol Evol 1, 14 (2016).

17. D. Becker, et al., Adaptive phenotypic plasticity is under stabilizing selection in Daphnia. Nat Ecol Evol 6, 1449–1457 (2022).

18. M. Kimura, J. F. Crow, Effect of overall phenotypic selection on genetic change at individual loci. Proc. Natl. Acad. Sci. U. S. A. 75, 6168–6171 (1978).

19. H. A. Orr, Fitness and its role in evolutionary genetics. Nat. Rev. Genet. 10, 531–539 (2009).

20. R. C. Lewontin, The triple helix: gene, organism and environment. Nat. Med. 6, 1206 (2000).

21. D. G. Williams, R. N. Mack, R. A. Black, Ecophysiology of introduced Pennisetum setaceum on Hawaii: The role of phenotypic plasticity. Ecology 76, 1569–1580 (1995).

22. E. B. Josephs, M. L. Van Etten, A. Harkess, A. Platts, R. S. Baucom, Adaptive and maladaptive expression plasticity underlying herbicide resistance in an agricultural weed. Evol Lett 5, 432–440 (2021).

23. J. P. Velotta, C. M. Ivy, C. J. Wolf, G. R. Scott, Z. A. Cheviron, Maladaptive phenotypic plasticity in cardiac muscle growth is suppressed in high-altitude deer mice. Evolution 72, 2712–2727 (2018).

24. C. K. Ghalambor, et al., Non-adaptive plasticity potentiates rapid adaptive evolution of gene expression in nature. Nature 525, 372–375 (2015).

25. J. F. Storz, A. M. Runck, H. Moriyama, R. E. Weber, A. Fago, Genetic differences in hemoglobin function between highland and lowland deer mice. J. Exp. Biol. 213, 2565–2574 (2010).

26. T. D. Price, A. Qvarnström, D. E. Irwin, The role of phenotypic plasticity in driving genetic evolution. Proc. Biol. Sci. 270, 1433–1440 (2003).

27. R. Lande, Adaptation to an extraordinary environment by evolution of phenotypic plasticity and genetic assimilation. J. Evol. Biol. 22, 1435–1446 (2009).

28. A. B. Paaby, M. V. Rockman, Cryptic genetic variation: evolution’s hidden substrate. Nat. Rev. Genet. 15, 247–258 (2014).

29. J. Hermisson, G. P. Wagner, The population genetic theory of hidden variation and genetic robustness. Genetics 168, 2271–2284 (2004).

30. U. Johanson, et al., Molecular analysis of FRIGIDA, a major determinant of natural variation in Arabidopsis flowering time. Science 290, 344–347 (2000).

31. L. Keren, et al., Massively Parallel Interrogation of the Effects of Gene Expression Levels on Fitness. Cell 166, 1282–1294.e18 (2016).

32. S.-A. A. Chen, A. F. Kern, R. M. L. Ang, Y. Xie, H. B. Fraser, Gene-by-environment interactions are pervasive among natural genetic variants. Cell Genom 3, 100273 (2023).

33. J. S. Rest, et al., Nonlinear fitness consequences of variation in expression level of a eukaryotic gene. Mol. Biol. Evol. 30, 448–456 (2013).

34. E. Sharon, et al., Functional Genetic Variants Revealed by Massively Parallel Precise Genome Editing. Cell 175, 544–557.e16 (2018).

35. F. Duveau, D. C. Yuan, B. P. H. Metzger, A. Hodgins-Davis, P. J. Wittkopp, Effects of mutation and selection on plasticity of a promoter activity in Saccharomyces cerevisiae. Proc. Natl. Acad. Sci. U. S. A. 114, E11218–E11227 (2017).

36. F. Duveau, W. Toubiana, P. J. Wittkopp, Fitness effects of cis-regulatory variants in the Saccharomyces cerevisiae TDH3 promoter. Mol. Biol. Evol. 34, 2908–2912 (2017).

37. F. Duveau, et al., Fitness effects of altering gene expression noise in Saccharomyces cerevisiae. Elife 7 (2018).

38. S. Kuroda, S. Otaka, Y. Fujisawa, Fermentable and nonfermentable carbon sources sustain constitutive levels of expression of yeast triosephosphate dehydrogenase 3 gene from distinct promoter elements. J. Biol. Chem. 269, 6153–6162 (1994).

39. G. A. Wray, et al., The evolution of transcriptional regulation in eukaryotes. Mol. Biol. Evol. 20, 1377–1419 (2003).

40. G. A. Wray, The evolutionary significance of cis-regulatory mutations. Nat. Rev. Genet. 8, 206–216 (2007).

41. P. J. Wittkopp, G. Kalay, Cis-regulatory elements: molecular mechanisms and evolutionary processes underlying divergence. Nat. Rev. Genet. 13, 59–69 (2011).

42. S. B. Carroll, Evo-devo and an expanding evolutionary synthesis: a genetic theory of morphological evolution. Cell 134, 25–36 (2008).

43. D. L. Stern, V. Orgogozo, The loci of evolution: how predictable is genetic evolution? Evolution 62, 2155–2177 (2008).

44. M. C. Kuang, et al., Repeated cis-regulatory tuning of a metabolic bottleneck gene during evolution. Mol. Biol. Evol. 35, 1968–1981 (2018).

45. J. Boocock, M. J. Sadhu, A. Durvasula, J. S. Bloom, L. Kruglyak, Ancient balancing selection maintains incompatible versions of the galactose pathway in yeast. Science 371, 415–419 (2021).

46. M. A. Siddiq, P. J. Wittkopp, Mechanisms of regulatory evolution in yeast. Curr. Opin. Genet. Dev. 77, 101998 (2022).

47. G. Gibson, L. K. Reed, Cryptic genetic variation. Curr. Biol. 18, R989–90 (2008).

48. S. L. Rutherford, S. Lindquist, Hsp90 as a capacitor for morphological evolution. Nature 396, 336–342 (1998).

49. C. Queitsch, T. A. Sangster, S. Lindquist, Hsp90 as a capacitor of phenotypic variation. Nature 417, 618–624 (2002).

50. A. M. Dean, J. W. Thornton, Mechanistic approaches to the study of evolution: the functional synthesis. Nat. Rev. Genet. 8, 675–688 (2007).

51. B. P. H. Metzger, et al., Contrasting Frequencies and Effects of cis- and trans-Regulatory Mutations Affecting Gene Expression. Mol. Biol. Evol. 33, 1131–1146 (2016).

