## Supplementary File 1 for "Plasticity and environment-specific relationships between gene expression and fitness in *Saccharomyces cerevisiae*": ExpressionFitnessAnalysis.html

- Figure 2
- Figure
  3A-B: Variation in gene expression among environments
- Figure
  3C-D: Variation in gene expression among environments
- Figure 4

This markdown file provides the code to reproduce the results
reported in Plasticity and environment-specific relationships between
gene expression and fitness in Saccharomyces cerevisiae. The data
necessary for the analyses are contained in SupplementaryTable1.xlsx.
The excel file contains the various .csv files necessary for to
reproduce the results as different sheets. Functions from the readxl
package allow the separate sheets to be directly imported, and the
sheets can be separated into .csv files and saved if necessary for
analyses in other programs. To use the script, install the packages and
adjust the paths to the files of interest accordingly; as written, the
code assumes that the SupplementaryTable1.xlsx is in the same directory
as the markdown file.

The first step is to load the data and the necessary packages.

```
# Load the necessary packages
library(tidyverse)
```

```
## ── Attaching core tidyverse packages ──────────────────────── tidyverse 2.0.0 ──
## ✔ dplyr     1.1.4     ✔ readr     2.1.5
## ✔ forcats   1.0.0     ✔ stringr   1.5.1
## ✔ ggplot2   3.5.0     ✔ tibble    3.2.1
## ✔ lubridate 1.9.3     ✔ tidyr     1.3.1
## ✔ purrr     1.0.2     
## ── Conflicts ────────────────────────────────────────── tidyverse_conflicts() ──
## ✖ dplyr::filter() masks stats::filter()
## ✖ dplyr::lag()    masks stats::lag()
## ℹ Use the conflicted package (<http://conflicted.r-lib.org/>) to force all conflicts to become errors
```

```
library(ggplot2)
library(ggsci)
library(ggthemes)
library(broom)
library(readxl)
library(gridExtra)
```

```
## 
## Attaching package: 'gridExtra'
## 
## The following object is masked from 'package:dplyr':
## 
##     combine
```

```
library(gridGraphics)
```

```
## Loading required package: grid
```

```
library(here)
```

```
## here() starts at /Users/msiddiq/Dropbox (University of Michigan)/PostDoc/Wittkopp Lab/Manuscripts/Environmental impacts on gene expression and fitness of cis-regulatory mutations in yeast/SupplementalFile
```

```
# Load the data
growth_data <- readxl::read_excel("./SupplementalTable1_bioRxiv.xlsx", sheet = "DoublingTime")

# The summary file for the expression/fitness associated with each promoter variant
ExpressionFitnessData_allEnvironments <- readxl::read_excel("./SupplementalTable1_bioRxiv.xlsx", sheet = "AllStrains-ExpressionFitness")
#Ensures numeric columns are properly formatted
ExpressionFitnessData_allEnvironments <- ExpressionFitnessData_allEnvironments %>% mutate_at(vars(YFP.MEDIAN.ADJUST.MEAN_driftcorrected, YFP.MEDIAN.ADJUST.SD_driftcorrected, YFP.MEDIAN.RELATIVE.MEAN, Fitness, Low.95, High.95,
                                                                                                  YFP.MEDIAN.RELATIVE.MEAN, YFP.MEDIAN.RELATIVE.SD), as.numeric)

# Data for replicates of the fitness assays and expression assays
FitnessData_allEnvironments <- readxl::read_excel("./SupplementalTable1_bioRxiv.xlsx", sheet = "FitnessData-Replicates")
Expression_allEnvironments_allGeno <- readxl::read_excel("./SupplementalTable1_bioRxiv.xlsx", sheet = "ExpressionData-Replicates")

## Make a folder to put the figures in. If the folder already exists, it will not create a new one.
if (!dir.exists("ManuscriptFigures"))
dir.create("ManuscriptFigures", showWarnings = TRUE)
```

### Figure 2

The following code generates the data panels in Figure 2A, 2C, and
2D. The figures will be exported to the “ManuscriptFigures” folder
(created above) as pdf files.

```
growth_data <- growth_data %>%
  rename(ENVIRONMENT = Environment) %>% #changing name to uppercase keeps column consistent with other files
  mutate(ENVIRONMENT = toupper(ENVIRONMENT))

growth_data$ENVIRONMENT <- factor(growth_data$ENVIRONMENT, levels = c("GLUCOSE", "GALACTOSE", "GLYCEROL", "ETHANOL"))
FitnessData_allEnvironments$ENVIRONMENT <- factor(FitnessData_allEnvironments$ENVIRONMENT, levels = c("GLUCOSE", "GALACTOSE", "GLYCEROL", "ETHANOL"))
Expression_allEnvironments_allGeno$ENVIRONMENT <- factor(Expression_allEnvironments_allGeno$ENVIRONMENT, levels = c("GLUCOSE", "GALACTOSE", "GLYCEROL", "ETHANOL"))

named_colors <- c("GLUCOSE" = "#0072B2", "GALACTOSE" = "#009E73", "GLYCEROL" = "#CC79A7", "ETHANOL" = "#D55E00")

# Plot of doubling time --------------------------------------------------------
plot_DoublingTime <- ggplot(growth_data, aes(x = ENVIRONMENT, y = 60/Doubling, col = ENVIRONMENT)) +
  geom_jitter(size = 3, width = 0.25, alpha = 0.5) + 
  ylim(0.2, 0.8) +
  theme_bw() +
  scale_color_manual(values = named_colors) +
  theme(legend.position = "none",
        panel.grid.major = element_blank(), panel.grid.minor = element_blank()) +
  theme(axis.text.x = element_text(angle = 45, hjust = 1)) +
  labs(x = "", y = "Growth Rate (doublings per hour)")


# Plot of expression effect among environments -----------------------------------------------------
plot_ExpressionReference_violin <- ggplot(subset(Expression_allEnvironments_allGeno, MUTATION.Expression == "REF"), 
                                   aes(x = ENVIRONMENT, y = YFP.MEDIAN.ADJUST_driftcorrected, color = ENVIRONMENT)) + 
  geom_point(position = position_jitterdodge(jitter.width = 0.4), alpha = 0.25)  +
  geom_violin(aes(fill = ENVIRONMENT), draw_quantiles = c(0.5), 
               position = position_dodge(width = 0.75), width = 0.5, alpha = 0.25, col = "Black") +  # Adjust width as needed#
  labs(x = "", y = "YFP Expression") + 
  theme_bw() +   scale_color_manual(values = named_colors) + scale_fill_manual(values = named_colors) +
  theme(panel.border = element_rect(linetype = 1, fill = "NA"),
        plot.background = element_rect(color = NA),legend.position="none",
        axis.text=element_text(size= 10), axis.title=element_text(size =14),
        panel.grid.minor = element_blank(),
        panel.grid.major = element_blank())

# Plot of deletion effect among environments-----------------------------
#Keep only observations for which the MUTATION = "BARRY" and "TDH3.Deletion"  from the fitness_summary data frame
fitness_data_DeletionEffect <- FitnessData_allEnvironments %>%
  filter(MUTATION.Fitness %in% c("REF", "TDH3.Deletion"))

plot_DeletionEffect_violin <- ggplot(fitness_data_DeletionEffect, aes(x = ENVIRONMENT, y = RelativeFitness, 
                                                               col = ENVIRONMENT, shape = MUTATION.Fitness)) +
  geom_hline(yintercept = 1, linetype = "dashed", color = "black") +
  scale_y_continuous(breaks = c(0.94, 0.96, 0.98, 1.0)) +
  geom_point(position = position_jitterdodge(jitter.width = 0.4), alpha = 0.8)  +
  geom_violin(col = "Black", aes(fill = ENVIRONMENT), alpha = 0.20,draw_quantiles = c(0.5)) +
  labs(x = "", y = "Relative Fitness") +
  theme_bw() +
  scale_color_manual(values = named_colors) +  
  scale_fill_manual(values = named_colors) +  
  theme(panel.grid.major = element_blank(), panel.grid.minor = element_blank()
        )


plot_DoublingTime
```

```
plot_ExpressionReference_violin
```

```
plot_DeletionEffect_violin
```

```
# Save the plots
ggsave("./ManuscriptFigures/Fig2_plot_ExpressionReference_violin.pdf", plot = plot_ExpressionReference_violin, width = 4.5, height = 4, units = "in", dpi = 300)
ggsave("./ManuscriptFigures/Fig2_plot_DoublingTime.pdf", plot = plot_DoublingTime, width = 3.0, height = 4)
ggsave("./ManuscriptFigures/Fig2_plot_DeletionEffect_violin.pdf", plot = plot_DeletionEffect_violin, width = 3, height = 4)
```

### Figure 3A-B: Variation in gene expression among environments

The following code produces the panels in Figure 3A and Figure 3B.
These are the fitness curves in the different environment. The x-axis
values (expression levels) have all been normalized by the value of the
unmutated, reference strain in the glucose environment.

```
#Take the ExpressionFitnessData_allEnvironments. Classify genotypes based on the number of their mutations. Create a column called NumPromMutation.
#The following strains have 1 mutation:Y1892,Y1896,Y1897,Y1898,Y1901,Y1903,Y1960,Y2528,Y2546,Y2566,Y2605,Y2651,Y2656
# The following strains have 2 mutations: Y2455,Y2458,Y2460,Y2465,Y2469,Y2475,Y2478,Y2653,Y2836,Y2842 
# The following strains have >2 mutations: Y2550,Y2812,Y2814,Y2816,Y2820,Y2824,Y2826,Y2828,Y2832,Y2834,Y2840
# The following strains have CNVs of YFP: Y2671,Y2675,Y2676,Y3006,Y3010,Y3011,Y3012,Y3015,Y3019,Y3021,Y3022,Y3025,Y3028,Y3034,Y3040

ExpressionFitnessData_allEnvironments_numProm <- ExpressionFitnessData_allEnvironments %>%
  mutate(NumPromMutation = case_when(
    STRAIN.Expression %in% c("Y1892","Y1896","Y1897","Y1898","Y1901","Y1903","Y1960","Y2528","Y2546","Y2566","Y2605","Y2651","Y2656") ~ "1",
    STRAIN.Expression %in% c("Y2455","Y2458","Y2460","Y2465","Y2469","Y2475","Y2478","Y2653","Y2836","Y2842") ~ "2",
    STRAIN.Expression %in% c("Y2550","Y2812","Y2814","Y2816","Y2820","Y2824","Y2826","Y2828","Y2832","Y2834","Y2840") ~ "> 2",
    STRAIN.Expression %in% c("Y2671","Y2675","Y2676","Y3006","Y3010","Y3011","Y3012","Y3015","Y3019","Y3021","Y3022","Y3025","Y3028","Y3034","Y3040") ~ "CNV",
      )) 

unmutated <- c("REF", "S288c", "NEG", "DUP")

ExpressionFitnessData_allEnvironments_numProm$ENVIRONMENT <- fct_relevel(ExpressionFitnessData_allEnvironments_numProm$ENVIRONMENT, c("GLUCOSE", "GALACTOSE", "GLYCEROL", "ETHANOL"))
ExpressionFitnessData_allEnvironments_numProm$NumPromMutation <- fct_relevel(ExpressionFitnessData_allEnvironments_numProm$NumPromMutation, 
                                                              c("0", "1", "2", "> 2", "CNV"))

### The function for doing pairwise t-tests between observations for each strain in different environments
### with bonferroni correction. 
### Test alternative approach -- using a function 
do_t_test <- function(df, env1, env2, adjustment_method = "bonferroni") {
  # Subset the data for the two environments
  data_subset <- df %>%
    filter(ENVIRONMENT %in% c(env1, env2)) %>%
    select(STRAIN.Expression, ENVIRONMENT, YFP.MEDIAN.RELATIVE) 
  
  # Perform the t-test within each strain and combination of environments
  t_test_results <- data_subset %>%
    group_by(STRAIN.Expression) %>%
    do(tidy(t.test(YFP.MEDIAN.RELATIVE ~ ENVIRONMENT, data = .)))
  
  # Extract the p-values and perform p-value adjustment
  p_values <- t_test_results %>%
    select(STRAIN.Expression, p.value) %>%
    mutate(p.value = p.adjust(p.value, method = adjustment_method)) %>%
    distinct()
  
  # Summarize the data and add the p-values
  summary_data <- data_subset %>%
    left_join(p_values, by = "STRAIN.Expression") %>%
    group_by(STRAIN.Expression, ENVIRONMENT) %>%
    summarise(YFP.MEDIAN.RELATIVE.MEAN = mean(YFP.MEDIAN.RELATIVE), 
              YFP.MEDIAN.RELATIVE.SD = sd(YFP.MEDIAN.RELATIVE),
              n = n(),
              p.adjusted = first(p.value)) %>%
    mutate(comp = paste(env1, "vs", env2),
           significance = if_else(p.adjusted < 0.05, "Significant", "Not significant"))
  
  return(summary_data)
}

## The pairwise environments to pass on to the function 
env_pairs <- list(
  c("GLUCOSE", "GALACTOSE"),
  c("GLUCOSE", "GLYCEROL"),
  c("GLUCOSE", "ETHANOL"),
  c("GALACTOSE", "GLYCEROL"),
  c("GALACTOSE", "ETHANOL"),
  c("GLYCEROL", "ETHANOL")
)

# For all of the data
all_results <- map_dfr(env_pairs, ~do_t_test(Expression_allEnvironments_allGeno, .x[1], .x[2]))
```

```
## `summarise()` has grouped output by 'STRAIN.Expression'. You can override using
## the `.groups` argument.
## `summarise()` has grouped output by 'STRAIN.Expression'. You can override using
## the `.groups` argument.
## `summarise()` has grouped output by 'STRAIN.Expression'. You can override using
## the `.groups` argument.
## `summarise()` has grouped output by 'STRAIN.Expression'. You can override using
## the `.groups` argument.
## `summarise()` has grouped output by 'STRAIN.Expression'. You can override using
## the `.groups` argument.
## `summarise()` has grouped output by 'STRAIN.Expression'. You can override using
## the `.groups` argument.
```

```
# Split into four data frames based on the comparison (in comp column). For each of the four datasets,
# convert the data into wide format such that there is a YFP.MEDIAN.RELATIVE.MEAN_GLU, 
# YFP.MEDIAN.RELATIVE.SD_GLU, YFP.MEDIAN.RELATIVE.MEAN_GAL, YFP.MEDIAN.RELATIVE.SD_GAL, etc.Keep the signifiance column,
# n, and p.adjusted column

data_GLU_GAL <- all_results %>%
  filter(comp == "GLUCOSE vs GALACTOSE") %>%
  pivot_wider(names_from = ENVIRONMENT, values_from = c(YFP.MEDIAN.RELATIVE.MEAN, YFP.MEDIAN.RELATIVE.SD))

data_GLU_GLY <- all_results %>%
  filter(comp == "GLUCOSE vs GLYCEROL") %>%
  pivot_wider(names_from = ENVIRONMENT, values_from = c(YFP.MEDIAN.RELATIVE.MEAN, YFP.MEDIAN.RELATIVE.SD))

data_GLU_ETH <- all_results %>%
  filter(comp == "GLUCOSE vs ETHANOL") %>%
  pivot_wider(names_from = ENVIRONMENT, values_from = c(YFP.MEDIAN.RELATIVE.MEAN, YFP.MEDIAN.RELATIVE.SD))

data_GAL_GLY <- all_results %>%
  filter(comp == "GALACTOSE vs GLYCEROL") %>%
  pivot_wider(names_from = ENVIRONMENT, values_from = c(YFP.MEDIAN.RELATIVE.MEAN, YFP.MEDIAN.RELATIVE.SD))

data_GAL_ETH <- all_results %>%
  filter(comp == "GALACTOSE vs ETHANOL") %>%
  pivot_wider(names_from = ENVIRONMENT, values_from = c(YFP.MEDIAN.RELATIVE.MEAN, YFP.MEDIAN.RELATIVE.SD))

data_GLY_ETH <- all_results %>%
  filter(comp == "GLYCEROL vs ETHANOL") %>%
  pivot_wider(names_from = ENVIRONMENT, values_from = c(YFP.MEDIAN.RELATIVE.MEAN, YFP.MEDIAN.RELATIVE.SD))

##### Plots---------
named_colors <- c("GLUCOSE" = "#0072B2", "GALACTOSE" = "#009E73", "GLYCEROL" = "#CC79A7", "ETHANOL" = "#D55E00")


# Generate scatter plots for each environment pair
plot_GLUvGAL_sig <- ggplot(data_GLU_GAL, aes(x = YFP.MEDIAN.RELATIVE.MEAN_GLUCOSE, y = YFP.MEDIAN.RELATIVE.MEAN_GALACTOSE, color = significance)) +
  geom_point(size = 2, alpha = 0.5) + #add standard error bars
  geom_errorbar(aes(ymin = YFP.MEDIAN.RELATIVE.MEAN_GALACTOSE - YFP.MEDIAN.RELATIVE.SD_GALACTOSE, 
                    ymax = YFP.MEDIAN.RELATIVE.MEAN_GALACTOSE + YFP.MEDIAN.RELATIVE.SD_GALACTOSE), 
                width = 0.0) + #add horizontal error bars
  geom_errorbarh(aes(xmin = YFP.MEDIAN.RELATIVE.MEAN_GLUCOSE - YFP.MEDIAN.RELATIVE.SD_GLUCOSE, 
                     xmax = YFP.MEDIAN.RELATIVE.MEAN_GLUCOSE + YFP.MEDIAN.RELATIVE.SD_GLUCOSE), 
                 height = 0.0) +
  geom_abline(intercept = 0, slope = 1, color = "black", linetype = "dashed", alpha = 0.2) +
  geom_hline(yintercept = 1, color = "grey") + geom_vline(xintercept = 1, color = "grey") +
  xlim(0, 2.1) + ylim(0, 2.1) +
  labs(x = "Relative Mean Expression in Glucose", y = "Relative Mean Expression in Galactose") + 
  scale_color_manual(values = c("Significant" = "red", "Not significant" = "black")) +
  theme_bw() +
  theme(legend.position = "bottom",
        axis.title = element_text(size = 16),
        axis.text.x = element_text(hjust = 1),
        panel.grid.minor = element_blank(),
        panel.grid.major = element_blank())

##Make the same plot as above for the other environment pairs
plot_GLUvGLY_sig <-ggplot(data_GLU_GLY, aes(x = YFP.MEDIAN.RELATIVE.MEAN_GLUCOSE, y = YFP.MEDIAN.RELATIVE.MEAN_GLYCEROL, color = significance)) +
  geom_point(size = 2, alpha = 0.5) + #add standard error bars
  geom_errorbar(aes(ymin = YFP.MEDIAN.RELATIVE.MEAN_GLYCEROL - YFP.MEDIAN.RELATIVE.SD_GLYCEROL, 
                    ymax = YFP.MEDIAN.RELATIVE.MEAN_GLYCEROL + YFP.MEDIAN.RELATIVE.SD_GLYCEROL), 
                width = 0.0) + #add horizontal error bars
  geom_errorbarh(aes(xmin = YFP.MEDIAN.RELATIVE.MEAN_GLUCOSE - YFP.MEDIAN.RELATIVE.SD_GLUCOSE, 
                     xmax = YFP.MEDIAN.RELATIVE.MEAN_GLUCOSE + YFP.MEDIAN.RELATIVE.SD_GLUCOSE), 
                 height = 0.0) +
  geom_abline(intercept = 0, slope = 1, color = "black", linetype = "dashed", alpha = 0.2) +
  geom_hline(yintercept = 1, color = "grey") + geom_vline(xintercept = 1, color = "grey") +
  xlim(0, 2.1) + ylim(0, 2.1) +
  labs(x = "Relative Mean Expression in Glucose", y = "Relative Mean Expression in Glycerol") + 
  scale_color_manual(values = c("Significant" = "red", "Not significant" = "black")) +
  theme_bw() +
  theme(legend.position = "bottom",
        axis.title = element_text(size = 16),
        axis.text.x = element_text(hjust = 1),
        panel.grid.minor = element_blank(),
        panel.grid.major = element_blank())

###
plot_GLUvETH_sig <- ggplot(data_GLU_ETH, aes(x = YFP.MEDIAN.RELATIVE.MEAN_GLUCOSE, y = YFP.MEDIAN.RELATIVE.MEAN_ETHANOL, color = significance)) +
  geom_point(size = 2, alpha = 0.5) + #add standard error bars
  geom_errorbar(aes(ymin = YFP.MEDIAN.RELATIVE.MEAN_ETHANOL - YFP.MEDIAN.RELATIVE.SD_ETHANOL, 
                    ymax = YFP.MEDIAN.RELATIVE.MEAN_ETHANOL + YFP.MEDIAN.RELATIVE.SD_ETHANOL), 
                width = 0.0) + #add horizontal error bars
  geom_errorbarh(aes(xmin = YFP.MEDIAN.RELATIVE.MEAN_GLUCOSE - YFP.MEDIAN.RELATIVE.SD_GLUCOSE, 
                     xmax = YFP.MEDIAN.RELATIVE.MEAN_GLUCOSE + YFP.MEDIAN.RELATIVE.SD_GLUCOSE), 
                 height = 0.0) +
  geom_abline(intercept = 0, slope = 1, color = "black", linetype = "dashed", alpha = 0.2) +
  geom_hline(yintercept = 1, color = "grey") + geom_vline(xintercept = 1, color = "grey") +
  xlim(0, 2.1) + ylim(0, 2.1) +
  labs(x = "Relative Mean Expression in Glucose", y = "Relative Mean Expression in Ethanol") + 
  scale_color_manual(values = c("Significant" = "red", "Not significant" = "black")) +
  theme_bw() +
  theme(legend.position = "bottom",
        axis.title = element_text(size = 16),
        axis.text.x = element_text(hjust = 1),
        panel.grid.minor = element_blank(),
        panel.grid.major = element_blank())

###
plot_GALvGLY_sig <- ggplot(data_GAL_GLY, aes(x = YFP.MEDIAN.RELATIVE.MEAN_GALACTOSE, y = YFP.MEDIAN.RELATIVE.MEAN_GLYCEROL, color = significance)) +
  geom_point(size = 2, alpha = 0.5) + #add standard error bars
  geom_errorbar(aes(ymin = YFP.MEDIAN.RELATIVE.MEAN_GLYCEROL - YFP.MEDIAN.RELATIVE.SD_GLYCEROL, 
                    ymax = YFP.MEDIAN.RELATIVE.MEAN_GLYCEROL + YFP.MEDIAN.RELATIVE.SD_GLYCEROL), 
                width = 0.0) + #add horizontal error bars
  geom_errorbarh(aes(xmin = YFP.MEDIAN.RELATIVE.MEAN_GALACTOSE - YFP.MEDIAN.RELATIVE.SD_GALACTOSE, 
                     xmax = YFP.MEDIAN.RELATIVE.MEAN_GALACTOSE + YFP.MEDIAN.RELATIVE.SD_GALACTOSE), 
                 height = 0.0) +
  geom_abline(intercept = 0, slope = 1, color = "black", linetype = "dashed", alpha = 0.2) +
  geom_hline(yintercept = 1, color = "grey") + geom_vline(xintercept = 1, color = "grey") +
  xlim(0, 2.1) + ylim(0, 2.1) +
  labs(x = "Relative Mean Expression in Galactose", y = "Relative Mean Expression in Glycerol") + 
  scale_color_manual(values = c("Significant" = "red", "Not significant" = "black")) +
  theme_bw() +
  theme(legend.position = "bottom",
        axis.title = element_text(size = 16),
        axis.text.x = element_text(hjust = 1),
        panel.grid.minor = element_blank(),
        panel.grid.major = element_blank())

###
plot_GALvETH_sig <- ggplot(data_GAL_ETH, aes(x = YFP.MEDIAN.RELATIVE.MEAN_GALACTOSE, y = YFP.MEDIAN.RELATIVE.MEAN_ETHANOL, color = significance)) +
  geom_point(size = 2, alpha = 0.5) + #add standard error bars
  geom_errorbar(aes(ymin = YFP.MEDIAN.RELATIVE.MEAN_ETHANOL - YFP.MEDIAN.RELATIVE.SD_ETHANOL, 
                    ymax = YFP.MEDIAN.RELATIVE.MEAN_ETHANOL + YFP.MEDIAN.RELATIVE.SD_ETHANOL), 
                width = 0.0) + #add horizontal error bars
  geom_errorbarh(aes(xmin = YFP.MEDIAN.RELATIVE.MEAN_GALACTOSE - YFP.MEDIAN.RELATIVE.SD_GALACTOSE, 
                     xmax = YFP.MEDIAN.RELATIVE.MEAN_GALACTOSE + YFP.MEDIAN.RELATIVE.SD_GALACTOSE), 
                 height = 0.0) +
  geom_abline(intercept = 0, slope = 1, color = "black", linetype = "dashed", alpha = 0.2) +
  geom_hline(yintercept = 1, color = "grey") + geom_vline(xintercept = 1, color = "grey") +
  xlim(0, 2.1) + ylim(0, 2.1) +
  labs(x = "Relative Mean Expression in Galactose", y = "Relative Mean Expression in Ethanol") + 
  scale_color_manual(values = c("Significant" = "red", "Not significant" = "black")) +
  theme_bw() +
  theme(legend.position = "bottom",
        axis.title = element_text(size = 16),
        axis.text.x = element_text(hjust = 1),
        panel.grid.minor = element_blank(),
        panel.grid.major = element_blank())


###
plot_GLYvETH_sig <- ggplot(data_GLY_ETH, aes(x = YFP.MEDIAN.RELATIVE.MEAN_GLYCEROL, y = YFP.MEDIAN.RELATIVE.MEAN_ETHANOL, color = significance)) +
  geom_point(size = 2, alpha = 0.5) + #add standard error bars
  geom_errorbar(aes(ymin = YFP.MEDIAN.RELATIVE.MEAN_ETHANOL - YFP.MEDIAN.RELATIVE.SD_ETHANOL, 
                    ymax = YFP.MEDIAN.RELATIVE.MEAN_ETHANOL + YFP.MEDIAN.RELATIVE.SD_ETHANOL), 
                width = 0.0) + #add horizontal error bars
  geom_errorbarh(aes(xmin = YFP.MEDIAN.RELATIVE.MEAN_GLYCEROL - YFP.MEDIAN.RELATIVE.SD_GLYCEROL, 
                     xmax = YFP.MEDIAN.RELATIVE.MEAN_GLYCEROL + YFP.MEDIAN.RELATIVE.SD_GLYCEROL), 
                 height = 0.0) +
  geom_abline(intercept = 0, slope = 1, color = "black", linetype = "dashed", alpha = 0.2) +
  geom_hline(yintercept = 1, color = "grey") + geom_vline(xintercept = 1, color = "grey") +
  xlim(0, 2.1) + ylim(0, 2.1) +
  labs(x = "Relative Mean Expression in Glycerol", y = "Relative Mean Expression in Ethanol") + 
  scale_color_manual(values = c("Significant" = "red", "Not significant" = "black")) +
  theme_bw() +
  theme(legend.position = "bottom",
        axis.title = element_text(size = 16),
        axis.text.x = element_text(hjust = 1),
        panel.grid.minor = element_blank(),
        panel.grid.major = element_blank())

###
plot_expMutants <- ggplot(subset(ExpressionFitnessData_allEnvironments_numProm, !(CLASS %in% unmutated)), 
       aes(x = NumPromMutation, y = YFP.MEDIAN.RELATIVE.MEAN, color = ENVIRONMENT)) + 
  geom_hline(yintercept = 1, linetype="dashed", color = "black") +
  geom_point(size = 1.5, alpha = 0.5, position = position_dodge(width = 0.5))  + 
  geom_errorbar(aes(ymin = YFP.MEDIAN.RELATIVE.MEAN - 1.96*YFP.MEDIAN.RELATIVE.SD/sqrt(4), 
                    ymax = YFP.MEDIAN.RELATIVE.MEAN + 1.96*YFP.MEDIAN.RELATIVE.SD/sqrt(4)), 
                width = 0.1,alpha = 0.5, position = position_dodge(width = 0.5)) +
  labs(x = "Mutant Genotypes", y = "Relative YFP Expression") +  
  #facet_grid(~NumPromMutation)+
  theme_bw() +   scale_color_manual(values = named_colors) + scale_fill_manual(values = named_colors) +
  theme(panel.border = element_rect(linetype = 1, fill = "NA"),
        plot.background = element_rect(color = NA),legend.position="top",
        axis.text=element_text(size= 10), axis.title=element_text(size =14),
        #        axis.text.x=element_text(angle=90,hjust=1),
        panel.grid.minor = element_blank(),
        panel.grid.major = element_blank())

plot_expMutants
```

```
plot_GLUvGAL_sig
```

```
plot_GLUvGLY_sig
```

```
plot_GLUvETH_sig
```

```
plot_GALvGLY_sig
```

```
plot_GALvETH_sig
```

```
plot_GLYvETH_sig
```

```
###Save the plots
ggsave("./ManuscriptFigures/Fig3_ExpressionEffects.pdf", plot = plot_expMutants, width = 8, height = 6, units = "in", dpi = 300)

ggsave("./ManuscriptFigures/Fig3_GLUvGAL.pdf", plot = plot_GLUvGAL_sig, width = 6, height = 6, units = "in", dpi = 300)
ggsave("./ManuscriptFigures/Fig3_GLUvGLY.pdf", plot = plot_GLUvGLY_sig, width = 6, height = 6, units = "in", dpi = 300)
ggsave("./ManuscriptFigures/Fig3_GLUvETH.pdf", plot = plot_GLUvETH_sig, width = 6, height = 6, units = "in", dpi = 300)
ggsave("./ManuscriptFigures/Fig3_GALvGLY.pdf", plot = plot_GALvGLY_sig, width = 6, height = 6, units = "in", dpi = 300)
ggsave("./ManuscriptFigures/Fig3_GALvETH.pdf", plot = plot_GALvETH_sig, width = 6, height = 6, units = "in", dpi = 300)
ggsave("./ManuscriptFigures/Fig3_GLYvETH.pdf", plot = plot_GLYvETH_sig, width = 6, height = 6, units = "in", dpi = 300)
```

### Figure 3C-D: Variation in gene expression among environments

The following code produces the panels in Figure 3C and Figure 3D.
These are the fitness curves in the different environment. The x-axis
values (expression levels) have all been normalized by the value of the
unmutated, reference strain in the glucose environment.

```
columns_to_keep <- c("YFP.MEDIAN.ADJUST.MEAN_driftcorrected", "YFP.MEDIAN.ADJUST.SD_driftcorrected", 
                     "YFP.MEDIAN.RELATIVE.MEAN", "MUTATION.Expression", "MUTATION.Fitness", "ENVIRONMENT", "Fitness", "ID", "STRAIN.Expression", "STRAIN.Fitness", "CLASS")

ExpressionFitnessData_allEnvironments$ENVIRONMENT <- fct_relevel(ExpressionFitnessData_allEnvironments$ENVIRONMENT, c("GLUCOSE", "GALACTOSE", "GLYCEROL", "ETHANOL"))

#There are many columns that have only numbers but are not treated as numeric. Change these columns to be numeric. Specifically, make the columns that only have numbers to be numeric.


ExpressionFitnessData_allEnvironments <- ExpressionFitnessData_allEnvironments %>% mutate_at(vars(YFP.MEDIAN.ADJUST.MEAN_driftcorrected, YFP.MEDIAN.ADJUST.SD_driftcorrected, YFP.MEDIAN.RELATIVE.MEAN, Fitness, Low.95, High.95,
                                                                                                  YFP.MEDIAN.RELATIVE.MEAN, YFP.MEDIAN.RELATIVE.SD), as.numeric)

#Weight the columns for LOWESS 
ExpressionFitnessData_allEnvironments_FC <- ExpressionFitnessData_allEnvironments %>% select(columns_to_keep)%>% 
  mutate(weights = ifelse(STRAIN.Fitness == "Y1189", 100, 
                          ifelse(STRAIN.Fitness == "Y1177", 100, 1))) %>%
  mutate(YFPmed_adjusted_RelToRefGLU = YFP.MEDIAN.ADJUST.MEAN_driftcorrected/subset(ExpressionFitnessData_allEnvironments, ENVIRONMENT == "GLUCOSE" & STRAIN.Fitness == "Y1189")$YFP.MEDIAN.ADJUST.MEAN_driftcorrected)

# Environment subsets
ExpressionFitnessData_GLU <- ExpressionFitnessData_allEnvironments_FC[ExpressionFitnessData_allEnvironments_FC$ENVIRONMENT == "GLUCOSE",]
ExpressionFitnessData_GAL <- ExpressionFitnessData_allEnvironments_FC[ExpressionFitnessData_allEnvironments_FC$ENVIRONMENT == "GALACTOSE",]
ExpressionFitnessData_GLY <- ExpressionFitnessData_allEnvironments_FC[ExpressionFitnessData_allEnvironments_FC$ENVIRONMENT == "GLYCEROL",]
ExpressionFitnessData_ETH <- ExpressionFitnessData_allEnvironments_FC[ExpressionFitnessData_allEnvironments_FC$ENVIRONMENT == "ETHANOL",]

### Modeling Fitness Curves --------
# Make a loess model with the weights relating YFP.MEDIAN.RELATIVE.MEAN and Fitness; The negative control
# and reference strain are weighted 100, while all other strains are weighted 1. These strains also have more
# measurements. 
model_GLU <- loess(Fitness ~ YFP.MEDIAN.RELATIVE.MEAN , data = ExpressionFitnessData_GLU, 
                   weights = weights)
model_GAL <- loess(Fitness ~ YFP.MEDIAN.RELATIVE.MEAN , data = ExpressionFitnessData_GAL, 
                   weights = weights)
model_GLY <- loess(Fitness ~ YFP.MEDIAN.RELATIVE.MEAN , data = ExpressionFitnessData_GLY, 
                   weights = weights)
model_ETH <- loess(Fitness ~ YFP.MEDIAN.RELATIVE.MEAN , data = ExpressionFitnessData_ETH, 
                   weights = weights)

## Visualize the curves
named_colors <- c("GLUCOSE" = "#0072B2", "GALACTOSE" = "#009E73", "GLYCEROL" = "#CC79A7", "ETHANOL" = "#D55E00")


plot_fitnessCurves_relGLU <- ggplot(ExpressionFitnessData_allEnvironments_FC, aes(x = YFPmed_adjusted_RelToRefGLU,
                                                                        y = Fitness, col = ENVIRONMENT)) + 
  geom_smooth(method = "loess", formula = y ~ x, se = TRUE, color = "#0072B2", fill = "#0072B2", size = 1, alpha = 0.2) + 
  geom_smooth(method = "loess", formula = y ~ x, se = TRUE, color = "#009E73", fill = "#009E73", size = 1, data = ExpressionFitnessData_GAL, alpha = 0.2) + 
  geom_smooth(method = "loess", formula = y ~ x, se = TRUE, color = "#CC79A7", fill = "#CC79A7", size = 1, data = ExpressionFitnessData_GLY, alpha = 0.2) + 
  geom_smooth(method = "loess", formula = y ~ x, se = TRUE, color = "#D55E00", fill = "#D55E00",size = 1, data = ExpressionFitnessData_ETH, alpha = 0.2) + 
  geom_hline(yintercept = 1, linetype = "dashed") + geom_vline(xintercept = 1, linetype = "dashed") +
  geom_point(data = ExpressionFitnessData_allEnvironments_FC[ExpressionFitnessData_allEnvironments_FC$STRAIN.Fitness == "Y1189",],
             aes(x = YFPmed_adjusted_RelToRefGLU, y = Fitness), 
              col = "black", size = 2.5) +
  scale_y_continuous(breaks = c(0.94, 0.97, 1.00, 1.03)) +
  theme_bw() +   scale_color_manual(values = named_colors) + scale_fill_manual(values = named_colors) +
  theme(legend.position = "bottom",
        axis.text.x = element_text(hjust = 1),
        panel.grid.minor = element_blank(),
        panel.grid.major = element_blank()) + 
  labs(x = "Expression Level (Normalized to Ref. Strain in Glucose)", y = "Relative Fitness")

plot_fitnessCurves_relGLU_facet_oneCol <- plot_fitnessCurves_relGLU + 
  geom_point(alpha = 0.5) +   
  facet_wrap(~factor(ENVIRONMENT, levels=c('GLUCOSE', 'GALACTOSE', 'GLYCEROL', 'ETHANOL')), ncol = 1) 


ggsave("./ManuscriptFigures/Fig3_fitnessCurves_overlain.pdf", plot = plot_fitnessCurves_relGLU, width = 6.0, height = 4.0, units = "in", dpi = 300)
ggsave("./ManuscriptFigures/Fig3_fitnessCurves_facet.pdf", plot = plot_fitnessCurves_relGLU_facet_oneCol, width = 6.0, height = 9, units = "in", dpi = 300)
```

### Figure 4

The following code produces the panels in Figure 4. The code also 1)
runs tests to check for normality for comparing expression and fitness
effects between TFBS and TATA box mutations and 2) runs the Mann-Whitney
U test to compare the expression effects fitness effects of TFBS and
TATA box mutations across environments that are reported in the
manuscripts. Note that t-test also appears appropriate for comparing the
expression effects of TFBS and TATA box mutations among environments and
results in the same inference.

```
# Strain numbers below correspond to "STRAIN.expression" column. The corresponding STRAIN.fitness with the mutations are also picked up
# using this approach.
mut_cisMotifs <- c("Y1896", "Y1897", "Y1898", "Y1960", "Y1892", "Y1901", "Y1903", #TFBS mutants; 
                   "Y2528", "Y2566", "Y2605", "Y2651", "Y2546", "Y2656", "Y2550") #TATA mutants


## Tests to detect effects on expression -------
Expression_cisMotifs <- ExpressionFitnessData_allEnvironments %>%
  filter(STRAIN.Expression %in% mut_cisMotifs) %>%  # Keep the necessary columns relevant for assessing fitness effects among environments
  select(STRAIN.Expression, STRAIN.Fitness, ENVIRONMENT, MUTATION.Expression, CLASS,YFP.MEDIAN.RELATIVE.MEAN ,YFP.MEDIAN.RELATIVE.SD) %>% #Change the columns with only numbers to numeric
  mutate(YFP.MEDIAN.RELATIVE.MEAN = as.numeric(YFP.MEDIAN.RELATIVE.MEAN),
         YFP.MEDIAN.RELATIVE.SD = as.numeric(YFP.MEDIAN.RELATIVE.SD))

Expression_cisMotifs_summary <- Expression_cisMotifs %>% 
  group_by(STRAIN.Expression, ENVIRONMENT, MUTATION.Expression, CLASS) %>% 
  summarise(YFP.MEDIAN.RELATIVE.MEAN = mean(YFP.MEDIAN.RELATIVE.MEAN),
            YFP.MEDIAN.RELATIVE.SD = mean(YFP.MEDIAN.RELATIVE.MEAN))
```

```
## `summarise()` has grouped output by 'STRAIN.Expression', 'ENVIRONMENT',
## 'MUTATION.Expression'. You can override using the `.groups` argument.
```

```
cisMotifs_var <- Expression_cisMotifs_summary %>%
  group_by(STRAIN.Expression, CLASS, MUTATION.Expression) %>%
  summarise(variance = var(YFP.MEDIAN.RELATIVE.MEAN))
```

```
## `summarise()` has grouped output by 'STRAIN.Expression', 'CLASS'. You can
## override using the `.groups` argument.
```

```
#Set the factor order
cisMotifs_var$CLASS <- fct_relevel(cisMotifs_var$CLASS, c("TFBS", "TATA"))

##Shapiro-Wilks normality test shows that variance is normally distributed 
## for the TFBS and TATA classes; t-test and F-test are appropriate. We will also 
## test with the MW test for robustness. 
shapiro.test(subset(cisMotifs_var, CLASS == "TFBS")$variance)
```

```
## 
##  Shapiro-Wilk normality test
## 
## data:  subset(cisMotifs_var, CLASS == "TFBS")$variance
## W = 0.91257, p-value = 0.414
```

```
shapiro.test(subset(cisMotifs_var, CLASS == "TATA")$variance)
```

```
## 
##  Shapiro-Wilk normality test
## 
## data:  subset(cisMotifs_var, CLASS == "TATA")$variance
## W = 0.93384, p-value = 0.584
```

```
t.test(variance ~ CLASS, data = cisMotifs_var, alternative = "greater")
```

```
## 
##  Welch Two Sample t-test
## 
## data:  variance by CLASS
## t = 2.3667, df = 6.5192, p-value = 0.02623
## alternative hypothesis: true difference in means between group TFBS and group TATA is greater than 0
## 95 percent confidence interval:
##  0.0005104549          Inf
## sample estimates:
## mean in group TFBS mean in group TATA 
##       0.0035317674       0.0008523344
```

```
wilcox.test(variance ~ CLASS, data = cisMotifs_var,
            alternative = "greater")  ## Also rejects the null hypothesis
```

```
## 
##  Wilcoxon rank sum exact test
## 
## data:  variance by CLASS
## W = 38, p-value = 0.04866
## alternative hypothesis: true location shift is greater than 0
```

```
## Test to detect effects on fitness -------
## Keep the columns relevant for assessing fitness effects among environments
FitnessData_cisMotifs <- ExpressionFitnessData_allEnvironments %>% 
  filter(STRAIN.Expression %in% mut_cisMotifs) %>%  # Keep the necessary columns relevant for assessing fitness effects among environments
  select(STRAIN.Expression, STRAIN.Fitness, ENVIRONMENT, MUTATION.Fitness, CLASS, Fitness, Low.95, High.95) 

#Calculate the mean RelativeFitness for each strain in each environment  
summary_FitnessData_cisMotifs <- FitnessData_cisMotifs %>%
  mutate(Fitness = as.numeric(gsub("E", "e", Fitness))) %>% 
  group_by(STRAIN.Fitness, ENVIRONMENT, CLASS, MUTATION.Fitness) %>% 
  summarise(meanRelativeFitness = mean(Fitness),
            sdRelativeFitness = sd(Fitness))
```

```
## `summarise()` has grouped output by 'STRAIN.Fitness', 'ENVIRONMENT', 'CLASS'.
## You can override using the `.groups` argument.
```

```
## Test if the meanRelativeFitness varies more for TFBS mutants than TATA mutants
## in the different environments
cisMut_varFitness <- summary_FitnessData_cisMotifs %>%
  group_by(STRAIN.Fitness, CLASS, MUTATION.Fitness) %>%
  summarise(variance = var(meanRelativeFitness))
```

```
## `summarise()` has grouped output by 'STRAIN.Fitness', 'CLASS'. You can override
## using the `.groups` argument.
```

```
shapiro.test(subset(cisMut_varFitness, CLASS == "TFBS")$variance)
```

```
## 
##  Shapiro-Wilk normality test
## 
## data:  subset(cisMut_varFitness, CLASS == "TFBS")$variance
## W = 0.88864, p-value = 0.2676
```

```
shapiro.test(subset(cisMut_varFitness, CLASS == "TATA")$variance)
```

```
## 
##  Shapiro-Wilk normality test
## 
## data:  subset(cisMut_varFitness, CLASS == "TATA")$variance
## W = 0.76193, p-value = 0.01688
```

```
# The Shapiro test suggests that t-tests are not appropriate for comparing variation 
# in fitness between TFBS and TATA mutants. The distribution of TFBS mutants is sufficiently
# normally distributed but the distribution of TATA box mutations is not. 
# We will use a Wilcoxon test to compare the variance in fitness between TFBS and TATA mutants

wilcox.test(variance ~ CLASS, data = cisMut_varFitness,
            alternative = "greater")
```

```
## 
##  Wilcoxon rank sum exact test
## 
## data:  variance by CLASS
## W = 32, p-value = 0.1914
## alternative hypothesis: true location shift is greater than 0
```

```
### Make plots showing mutational effects on expression and fitness across environments
## Reorder factors for plotting aesthetics ------
strain_order <- c("Y1896", "Y1897", "Y1898", "Y1960", "Y1892", "Y1901", "Y1903", #TFBS 
                  "Y2656", "Y2550", "Y2651", "Y2528", "Y2566", "Y2546", "Y2605")

FitnessData_cisMotifs$STRAIN.Expression <- fct_relevel(FitnessData_cisMotifs$STRAIN.Expression, strain_order)
summary_FitnessData_cisMotifs$ENVIRONMENT <- fct_relevel(summary_FitnessData_cisMotifs$ENVIRONMENT, c("GLUCOSE", "GALACTOSE", "GLYCEROL", "ETHANOL"))
summary_FitnessData_cisMotifs$CLASS <- fct_relevel(summary_FitnessData_cisMotifs$CLASS, c("TFBS", "TATA"))


Expression_cisMotifs$ENVIRONMENT <- fct_relevel(Expression_cisMotifs$ENVIRONMENT, c("GLUCOSE", "GALACTOSE", "GLYCEROL", "ETHANOL"))
Expression_cisMotifs$CLASS <- fct_relevel(Expression_cisMotifs$CLASS, c("TFBS", "TATA"))
cisMut_varFitness$CLASS <- fct_relevel(cisMotifs_var$CLASS, c("TFBS", "TATA"))


# For color consistency 
named_colors <- c("GLUCOSE" = "#0072B2", "GALACTOSE" = "#009E73", "GLYCEROL" = "#CC79A7", "ETHANOL" = "#D55E00")
motif_colors <- c("m63" = "#FBB913", "m66" = "#FBB913",
                  "m75" = "#6DC8BF", "m76" = "#6DC8BF",
                  "m89" = "#B72467", "m90" ="#B72467", "m91" = "#B72467") #RAP1, GCR1a, GCR1b, TATA colors

## Plots--expression effects of TFBS vs TATA -----
plot_expTFBSvTATA_linkedFacet <- ggplot(Expression_cisMotifs, aes(x = ENVIRONMENT, y = YFP.MEDIAN.RELATIVE.MEAN)) + 
  geom_line(aes(group = STRAIN.Expression), alpha = 0.5) +  
  geom_errorbar(aes(ymin = YFP.MEDIAN.RELATIVE.MEAN - YFP.MEDIAN.RELATIVE.SD, 
                    ymax = YFP.MEDIAN.RELATIVE.MEAN + YFP.MEDIAN.RELATIVE.SD, color = ENVIRONMENT), 
                width = 0.2, alpha = 0.5) +
  geom_point(aes(color = MUTATION.Expression), size = 2.5) + 
  theme_bw() + labs(x = "", y = "Relative Expression Level to Reference (afu)") +
  scale_color_manual(values = motif_colors) + #scale_fill_manual(values = motif_colors) +
  theme(panel.border = element_rect(linetype = 1, fill = "NA"),
        plot.background = element_rect(color = NA),legend.position="top",
        axis.text=element_text(size= 10), axis.title=element_text(size =14),
        axis.text.x=element_text(angle=45,hjust=1),
        panel.grid.minor = element_blank(),
        panel.grid.major = element_blank()) + 
  facet_wrap(~CLASS, ncol = 2, scales = "free_x",strip.position = "top") 

# Plot the variation between TFBS and TATA mutants
plot_expTFBSvTATA_variance <- ggplot(cisMotifs_var, aes(x = CLASS, y = variance)) +
  geom_boxplot(outlier.shape = NA, width = 0.5) +
  geom_jitter(aes(col = MUTATION.Expression), width = 0.15, size = 2.5) +
  scale_color_manual(values = motif_colors) + #scale_fill_manual(values = named_colors) +
  theme_bw() + labs(x = "", y = "Expression Effect Variance") +
  scale_fill_manual(values = named_colors) + 
  theme(panel.border = element_rect(linetype = 1, fill = "NA"),
        plot.background = element_rect(color = NA),legend.position="none",
        axis.text=element_text(size= 10), axis.title=element_text(size =14),
        axis.text.x=element_text(angle=45,hjust=1),
        panel.grid.minor = element_blank(),
        panel.grid.major = element_blank())

## Plots--fitness effects of TFBS vs TATA -----
plot_fitTFBSvTATA_linkedFacet <- ggplot(summary_FitnessData_cisMotifs, 
                                        aes(x = ENVIRONMENT, y = meanRelativeFitness)) + 
  geom_line(aes(group = STRAIN.Fitness)) +
  geom_errorbar(aes(ymin = meanRelativeFitness - sdRelativeFitness, 
                    ymax = meanRelativeFitness + sdRelativeFitness, color = ENVIRONMENT), width = 0.2, alpha = 0.5) +
  geom_point(aes(color = MUTATION.Fitness), size = 2.5) + 
  theme_bw() + labs(x = "", y = "Relative Fitness") +
  scale_color_manual(values = motif_colors) + #scale_fill_manual(values = named_colors) +
  theme(panel.border = element_rect(linetype = 1, fill = "NA"),
        plot.background = element_rect(color = NA),legend.position="top",
        axis.text=element_text(size= 10), axis.title=element_text(size =14),
        axis.text.x=element_text(angle=45,hjust=1),
        panel.grid.minor = element_blank(),
        panel.grid.major = element_blank()) + 
  facet_wrap(~CLASS, ncol = 2, scales = "free_x",strip.position = "top")+   
  ylim(0.94, 1.005) 

# Plot the variation between TFBS and TATA mutants
plot_fitTFBSvTATA_variance <- ggplot(cisMut_varFitness, aes(x = CLASS, y = variance)) +
  geom_boxplot(outlier.shape = NA, width = 0.5) +
  geom_jitter(aes(col = MUTATION.Fitness), width = 0.15, size = 2.5) +
  scale_color_manual(values = motif_colors) + #scale_fill_manual(values = named_colors) +
  theme_bw() + labs(x = "", y = "Fitness Effect Variance") +
  scale_fill_manual(values = named_colors) + 
  theme(panel.border = element_rect(linetype = 1, fill = "NA"),
        plot.background = element_rect(color = NA),legend.position="none",
        axis.text=element_text(size= 10), axis.title=element_text(size =14),
        axis.text.x=element_text(angle=45,hjust=1),
        panel.grid.minor = element_blank(),
        panel.grid.major = element_blank())

plot_expTFBSvTATA_linkedFacet
```

```
plot_expTFBSvTATA_variance
```

```
plot_fitTFBSvTATA_linkedFacet
```

```
plot_fitTFBSvTATA_variance
```

```
#Save the plot
ggsave("./ManuscriptFigures/Figure4_plot_expTFBSvTATA_linkedFacet.pdf", plot = plot_expTFBSvTATA_linkedFacet, width = 7, height = 5)
ggsave("./ManuscriptFigures/Figure4_plot_expTFBSvTATA_variance.pdf", plot_expTFBSvTATA_variance, width = 2.5, height = 4, units = "in")

ggsave("./ManuscriptFigures/Figure4_plot_fitTFBSvTATA_linkedFacet.pdf", plot_fitTFBSvTATA_linkedFacet, width = 7, height = 5, units = "in")
ggsave("./ManuscriptFigures/Figure4_plot_fitTFBSvTATA_variance.pdf", plot_fitTFBSvTATA_variance, width = 2.5, height = 4, units = "in")
```
