## Supplementary File 1 for "Plasticity and environment-specific relationships between gene expression and fitness in *Saccharomyces cerevisiae*": ExpressionFitnessAnalysis.tex

% Options for packages loaded elsewhere
\PassOptionsToPackage{unicode}{hyperref}
\PassOptionsToPackage{hyphens}{url}
%
\documentclass[
]{article}
\usepackage{amsmath,amssymb}
\usepackage{iftex}
\ifPDFTeX
  \usepackage[T1]{fontenc}
  \usepackage[utf8]{inputenc}
  \usepackage{textcomp} % provide euro and other symbols
\else % if luatex or xetex
  \usepackage{unicode-math} % this also loads fontspec
  \defaultfontfeatures{Scale=MatchLowercase}
  \defaultfontfeatures[\rmfamily]{Ligatures=TeX,Scale=1}
\fi
\usepackage{lmodern}
\ifPDFTeX\else
  % xetex/luatex font selection
\fi
% Use upquote if available, for straight quotes in verbatim environments
\IfFileExists{upquote.sty}{\usepackage{upquote}}{}
\IfFileExists{microtype.sty}{% use microtype if available
  \usepackage[]{microtype}
  \UseMicrotypeSet[protrusion]{basicmath} % disable protrusion for tt fonts
}{}
\makeatletter
\@ifundefined{KOMAClassName}{% if non-KOMA class
  \IfFileExists{parskip.sty}{%
    \usepackage{parskip}
  }{% else
    \setlength{\parindent}{0pt}
    \setlength{\parskip}{6pt plus 2pt minus 1pt}}
}{% if KOMA class
  \KOMAoptions{parskip=half}}
\makeatother
\usepackage{xcolor}
\usepackage[margin=1in]{geometry}
\usepackage{color}
\usepackage{fancyvrb}
\newcommand{\VerbBar}{|}
\newcommand{\VERB}{\Verb[commandchars=\\\{\}]}
\DefineVerbatimEnvironment{Highlighting}{Verbatim}{commandchars=\\\{\}}
% Add ',fontsize=\small' for more characters per line
\usepackage{framed}
\definecolor{shadecolor}{RGB}{248,248,248}
\newenvironment{Shaded}{\begin{snugshade}}{\end{snugshade}}
\newcommand{\AlertTok}[1]{\textcolor[rgb]{0.94,0.16,0.16}{#1}}
\newcommand{\AnnotationTok}[1]{\textcolor[rgb]{0.56,0.35,0.01}{\textbf{\textit{#1}}}}
\newcommand{\AttributeTok}[1]{\textcolor[rgb]{0.13,0.29,0.53}{#1}}
\newcommand{\BaseNTok}[1]{\textcolor[rgb]{0.00,0.00,0.81}{#1}}
\newcommand{\BuiltInTok}[1]{#1}
\newcommand{\CharTok}[1]{\textcolor[rgb]{0.31,0.60,0.02}{#1}}
\newcommand{\CommentTok}[1]{\textcolor[rgb]{0.56,0.35,0.01}{\textit{#1}}}
\newcommand{\CommentVarTok}[1]{\textcolor[rgb]{0.56,0.35,0.01}{\textbf{\textit{#1}}}}
\newcommand{\ConstantTok}[1]{\textcolor[rgb]{0.56,0.35,0.01}{#1}}
\newcommand{\ControlFlowTok}[1]{\textcolor[rgb]{0.13,0.29,0.53}{\textbf{#1}}}
\newcommand{\DataTypeTok}[1]{\textcolor[rgb]{0.13,0.29,0.53}{#1}}
\newcommand{\DecValTok}[1]{\textcolor[rgb]{0.00,0.00,0.81}{#1}}
\newcommand{\DocumentationTok}[1]{\textcolor[rgb]{0.56,0.35,0.01}{\textbf{\textit{#1}}}}
\newcommand{\ErrorTok}[1]{\textcolor[rgb]{0.64,0.00,0.00}{\textbf{#1}}}
\newcommand{\ExtensionTok}[1]{#1}
\newcommand{\FloatTok}[1]{\textcolor[rgb]{0.00,0.00,0.81}{#1}}
\newcommand{\FunctionTok}[1]{\textcolor[rgb]{0.13,0.29,0.53}{\textbf{#1}}}
\newcommand{\ImportTok}[1]{#1}
\newcommand{\InformationTok}[1]{\textcolor[rgb]{0.56,0.35,0.01}{\textbf{\textit{#1}}}}
\newcommand{\KeywordTok}[1]{\textcolor[rgb]{0.13,0.29,0.53}{\textbf{#1}}}
\newcommand{\NormalTok}[1]{#1}
\newcommand{\OperatorTok}[1]{\textcolor[rgb]{0.81,0.36,0.00}{\textbf{#1}}}
\newcommand{\OtherTok}[1]{\textcolor[rgb]{0.56,0.35,0.01}{#1}}
\newcommand{\PreprocessorTok}[1]{\textcolor[rgb]{0.56,0.35,0.01}{\textit{#1}}}
\newcommand{\RegionMarkerTok}[1]{#1}
\newcommand{\SpecialCharTok}[1]{\textcolor[rgb]{0.81,0.36,0.00}{\textbf{#1}}}
\newcommand{\SpecialStringTok}[1]{\textcolor[rgb]{0.31,0.60,0.02}{#1}}
\newcommand{\StringTok}[1]{\textcolor[rgb]{0.31,0.60,0.02}{#1}}
\newcommand{\VariableTok}[1]{\textcolor[rgb]{0.00,0.00,0.00}{#1}}
\newcommand{\VerbatimStringTok}[1]{\textcolor[rgb]{0.31,0.60,0.02}{#1}}
\newcommand{\WarningTok}[1]{\textcolor[rgb]{0.56,0.35,0.01}{\textbf{\textit{#1}}}}
\usepackage{graphicx}
\makeatletter
\def\maxwidth{\ifdim\Gin@nat@width>\linewidth\linewidth\else\Gin@nat@width\fi}
\def\maxheight{\ifdim\Gin@nat@height>\textheight\textheight\else\Gin@nat@height\fi}
\makeatother
% Scale images if necessary, so that they will not overflow the page
% margins by default, and it is still possible to overwrite the defaults
% using explicit options in \includegraphics[width, height, ...]{}
\setkeys{Gin}{width=\maxwidth,height=\maxheight,keepaspectratio}
% Set default figure placement to htbp
\makeatletter
\def\fps@figure{htbp}
\makeatother
\setlength{\emergencystretch}{3em} % prevent overfull lines
\providecommand{\tightlist}{%
  \setlength{\itemsep}{0pt}\setlength{\parskip}{0pt}}
\setcounter{secnumdepth}{-\maxdimen} % remove section numbering
\ifLuaTeX
  \usepackage{selnolig}  % disable illegal ligatures
\fi
\IfFileExists{bookmark.sty}{\usepackage{bookmark}}{\usepackage{hyperref}}
\IfFileExists{xurl.sty}{\usepackage{xurl}}{} % add URL line breaks if available
\urlstyle{same}
\hypersetup{
  hidelinks,
  pdfcreator={LaTeX via pandoc}}

\author{}
\date{\vspace{-2.5em}}

\begin{document}

{
\setcounter{tocdepth}{2}
\tableofcontents
}
This markdown file provides the code to reproduce the results reported
in Plasticity and environment-specific relationships between gene
expression and fitness in Saccharomyces cerevisiae. The data necessary
for the analyses are contained in SupplementaryTable1.xlsx. The excel
file contains the various .csv files necessary for to reproduce the
results in as different sheets. They can be separated into .csv files
using the readxl package in R, if necessary for analyses in other
programs.

The first step is to load the data and the necessary packages.

\begin{Shaded}
\begin{Highlighting}[]
\FunctionTok{setwd}\NormalTok{(}\StringTok{"\textasciitilde{}/Dropbox (University of Michigan)/PostDoc/Wittkopp Lab/Manuscripts/Environmental impacts on gene expression and fitness of cis{-}regulatory mutations in yeast/SupplementalFile/"}\NormalTok{)}

\CommentTok{\# Load the necessary packages}
\FunctionTok{library}\NormalTok{(tidyverse)}
\end{Highlighting}
\end{Shaded}

\begin{verbatim}
## -- Attaching core tidyverse packages ------------------------ tidyverse 2.0.0 --
## v dplyr     1.1.4     v readr     2.1.5
## v forcats   1.0.0     v stringr   1.5.1
## v ggplot2   3.5.0     v tibble    3.2.1
## v lubridate 1.9.3     v tidyr     1.3.1
## v purrr     1.0.2     
## -- Conflicts ------------------------------------------ tidyverse_conflicts() --
## x dplyr::filter() masks stats::filter()
## x dplyr::lag()    masks stats::lag()
## i Use the conflicted package (<http://conflicted.r-lib.org/>) to force all conflicts to become errors
\end{verbatim}

\begin{Shaded}
\begin{Highlighting}[]
\FunctionTok{library}\NormalTok{(ggplot2)}
\FunctionTok{library}\NormalTok{(ggsci)}
\FunctionTok{library}\NormalTok{(ggthemes)}
\FunctionTok{library}\NormalTok{(broom)}
\FunctionTok{library}\NormalTok{(readxl)}
\FunctionTok{library}\NormalTok{(gridExtra)}
\end{Highlighting}
\end{Shaded}

\begin{verbatim}
## 
## Attaching package: 'gridExtra'
## 
## The following object is masked from 'package:dplyr':
## 
##     combine
\end{verbatim}

\begin{Shaded}
\begin{Highlighting}[]
\FunctionTok{library}\NormalTok{(gridGraphics)}
\end{Highlighting}
\end{Shaded}

\begin{verbatim}
## Loading required package: grid
\end{verbatim}

\begin{Shaded}
\begin{Highlighting}[]
\CommentTok{\# Load the data}
\NormalTok{growth\_data }\OtherTok{\textless{}{-}}\NormalTok{ readxl}\SpecialCharTok{::}\FunctionTok{read\_excel}\NormalTok{(}\StringTok{"./SupplementalTable1\_bioRxiv.xlsx"}\NormalTok{, }\AttributeTok{sheet =} \StringTok{"DoublingTime"}\NormalTok{)}

\CommentTok{\# The summary file for the expression/fitness associated with each promoter variant}
\NormalTok{ExpressionFitnessData\_allEnvironments }\OtherTok{\textless{}{-}}\NormalTok{ readxl}\SpecialCharTok{::}\FunctionTok{read\_excel}\NormalTok{(}\StringTok{"./SupplementalTable1\_bioRxiv.xlsx"}\NormalTok{, }\AttributeTok{sheet =} \StringTok{"AllStrains{-}ExpressionFitness"}\NormalTok{)}
\CommentTok{\#Ensures numeric columns are properly formatted}
\NormalTok{ExpressionFitnessData\_allEnvironments }\OtherTok{\textless{}{-}}\NormalTok{ ExpressionFitnessData\_allEnvironments }\SpecialCharTok{\%\textgreater{}\%} \FunctionTok{mutate\_at}\NormalTok{(}\FunctionTok{vars}\NormalTok{(YFP.MEDIAN.ADJUST.MEAN\_driftcorrected, YFP.MEDIAN.ADJUST.SD\_driftcorrected, YFP.MEDIAN.RELATIVE.MEAN, Fitness, Low}\FloatTok{.95}\NormalTok{, High}\FloatTok{.95}\NormalTok{,}
\NormalTok{                                                                                                  YFP.MEDIAN.RELATIVE.MEAN, YFP.MEDIAN.RELATIVE.SD), as.numeric)}
\end{Highlighting}
\end{Shaded}

\begin{verbatim}
## Warning: There were 7 warnings in `mutate()`.
## The first warning was:
## i In argument: `YFP.MEDIAN.ADJUST.MEAN_driftcorrected =
##   .Primitive("as.double")(YFP.MEDIAN.ADJUST.MEAN_driftcorrected)`.
## Caused by warning:
## ! NAs introduced by coercion
## i Run `dplyr::last_dplyr_warnings()` to see the 6 remaining warnings.
\end{verbatim}

\begin{Shaded}
\begin{Highlighting}[]
\CommentTok{\# Data for replicates of the fitness assays and expression assays}
\NormalTok{FitnessData\_allEnvironments }\OtherTok{\textless{}{-}}\NormalTok{ readxl}\SpecialCharTok{::}\FunctionTok{read\_excel}\NormalTok{(}\StringTok{"./SupplementalTable1\_bioRxiv.xlsx"}\NormalTok{, }\AttributeTok{sheet =} \StringTok{"FitnessData{-}Replicates"}\NormalTok{)}
\NormalTok{Expression\_allEnvironments\_allGeno }\OtherTok{\textless{}{-}}\NormalTok{ readxl}\SpecialCharTok{::}\FunctionTok{read\_excel}\NormalTok{(}\StringTok{"./SupplementalTable1\_bioRxiv.xlsx"}\NormalTok{, }\AttributeTok{sheet =} \StringTok{"ExpressionData{-}Replicates"}\NormalTok{)}

\DocumentationTok{\#\# Make a folder to put the figures in. If the folder already exists, it will not create a new one.}
\ControlFlowTok{if}\NormalTok{ (}\SpecialCharTok{!}\FunctionTok{dir.exists}\NormalTok{(}\StringTok{"ManuscriptFigures"}\NormalTok{))}
\FunctionTok{dir.create}\NormalTok{(}\StringTok{"ManuscriptFigures"}\NormalTok{, }\AttributeTok{showWarnings =} \ConstantTok{TRUE}\NormalTok{)  }
\end{Highlighting}
\end{Shaded}

\hypertarget{figure-2}{%
\subsubsection{Figure 2}\label{figure-2}}

The following code generates the data panels in Figure 2A, 2C, and 2D.
The figures will be exported to the ``ManuscriptFigures'' folder
(created above) as pdf files.

\begin{Shaded}
\begin{Highlighting}[]
\NormalTok{growth\_data }\OtherTok{\textless{}{-}}\NormalTok{ growth\_data }\SpecialCharTok{\%\textgreater{}\%}
  \FunctionTok{rename}\NormalTok{(}\AttributeTok{ENVIRONMENT =}\NormalTok{ Environment) }\SpecialCharTok{\%\textgreater{}\%} \CommentTok{\#changing name to uppercase keeps column consistent with other files}
  \FunctionTok{mutate}\NormalTok{(}\AttributeTok{ENVIRONMENT =} \FunctionTok{toupper}\NormalTok{(ENVIRONMENT))}

\NormalTok{growth\_data}\SpecialCharTok{$}\NormalTok{ENVIRONMENT }\OtherTok{\textless{}{-}} \FunctionTok{factor}\NormalTok{(growth\_data}\SpecialCharTok{$}\NormalTok{ENVIRONMENT, }\AttributeTok{levels =} \FunctionTok{c}\NormalTok{(}\StringTok{"GLUCOSE"}\NormalTok{, }\StringTok{"GALACTOSE"}\NormalTok{, }\StringTok{"GLYCEROL"}\NormalTok{, }\StringTok{"ETHANOL"}\NormalTok{))}
\NormalTok{FitnessData\_allEnvironments}\SpecialCharTok{$}\NormalTok{ENVIRONMENT }\OtherTok{\textless{}{-}} \FunctionTok{factor}\NormalTok{(FitnessData\_allEnvironments}\SpecialCharTok{$}\NormalTok{ENVIRONMENT, }\AttributeTok{levels =} \FunctionTok{c}\NormalTok{(}\StringTok{"GLUCOSE"}\NormalTok{, }\StringTok{"GALACTOSE"}\NormalTok{, }\StringTok{"GLYCEROL"}\NormalTok{, }\StringTok{"ETHANOL"}\NormalTok{))}
\NormalTok{Expression\_allEnvironments\_allGeno}\SpecialCharTok{$}\NormalTok{ENVIRONMENT }\OtherTok{\textless{}{-}} \FunctionTok{factor}\NormalTok{(Expression\_allEnvironments\_allGeno}\SpecialCharTok{$}\NormalTok{ENVIRONMENT, }\AttributeTok{levels =} \FunctionTok{c}\NormalTok{(}\StringTok{"GLUCOSE"}\NormalTok{, }\StringTok{"GALACTOSE"}\NormalTok{, }\StringTok{"GLYCEROL"}\NormalTok{, }\StringTok{"ETHANOL"}\NormalTok{))}

\NormalTok{named\_colors }\OtherTok{\textless{}{-}} \FunctionTok{c}\NormalTok{(}\StringTok{"GLUCOSE"} \OtherTok{=} \StringTok{"\#0072B2"}\NormalTok{, }\StringTok{"GALACTOSE"} \OtherTok{=} \StringTok{"\#009E73"}\NormalTok{, }\StringTok{"GLYCEROL"} \OtherTok{=} \StringTok{"\#CC79A7"}\NormalTok{, }\StringTok{"ETHANOL"} \OtherTok{=} \StringTok{"\#D55E00"}\NormalTok{)}

\CommentTok{\# Plot of doubling time {-}{-}{-}{-}{-}{-}{-}{-}{-}{-}{-}{-}{-}{-}{-}{-}{-}{-}{-}{-}{-}{-}{-}{-}{-}{-}{-}{-}{-}{-}{-}{-}{-}{-}{-}{-}{-}{-}{-}{-}{-}{-}{-}{-}{-}{-}{-}{-}{-}{-}{-}{-}{-}{-}{-}{-}}
\NormalTok{plot\_DoublingTime }\OtherTok{\textless{}{-}} \FunctionTok{ggplot}\NormalTok{(growth\_data, }\FunctionTok{aes}\NormalTok{(}\AttributeTok{x =}\NormalTok{ ENVIRONMENT, }\AttributeTok{y =} \DecValTok{60}\SpecialCharTok{/}\NormalTok{Doubling, }\AttributeTok{col =}\NormalTok{ ENVIRONMENT)) }\SpecialCharTok{+}
  \FunctionTok{geom\_jitter}\NormalTok{(}\AttributeTok{size =} \DecValTok{3}\NormalTok{, }\AttributeTok{width =} \FloatTok{0.25}\NormalTok{, }\AttributeTok{alpha =} \FloatTok{0.5}\NormalTok{) }\SpecialCharTok{+} 
  \FunctionTok{ylim}\NormalTok{(}\FloatTok{0.2}\NormalTok{, }\FloatTok{0.8}\NormalTok{) }\SpecialCharTok{+}
  \FunctionTok{theme\_bw}\NormalTok{() }\SpecialCharTok{+}
  \FunctionTok{scale\_color\_manual}\NormalTok{(}\AttributeTok{values =}\NormalTok{ named\_colors) }\SpecialCharTok{+}
  \FunctionTok{theme}\NormalTok{(}\AttributeTok{legend.position =} \StringTok{"none"}\NormalTok{,}
        \AttributeTok{panel.grid.major =} \FunctionTok{element\_blank}\NormalTok{(), }\AttributeTok{panel.grid.minor =} \FunctionTok{element\_blank}\NormalTok{()) }\SpecialCharTok{+}
  \FunctionTok{theme}\NormalTok{(}\AttributeTok{axis.text.x =} \FunctionTok{element\_text}\NormalTok{(}\AttributeTok{angle =} \DecValTok{45}\NormalTok{, }\AttributeTok{hjust =} \DecValTok{1}\NormalTok{)) }\SpecialCharTok{+}
  \FunctionTok{labs}\NormalTok{(}\AttributeTok{x =} \StringTok{""}\NormalTok{, }\AttributeTok{y =} \StringTok{"Growth Rate (doublings per hour)"}\NormalTok{)}


\CommentTok{\# Plot of expression effect among environments {-}{-}{-}{-}{-}{-}{-}{-}{-}{-}{-}{-}{-}{-}{-}{-}{-}{-}{-}{-}{-}{-}{-}{-}{-}{-}{-}{-}{-}{-}{-}{-}{-}{-}{-}{-}{-}{-}{-}{-}{-}{-}{-}{-}{-}{-}{-}{-}{-}{-}{-}{-}{-}}
\NormalTok{plot\_ExpressionReference\_violin }\OtherTok{\textless{}{-}} \FunctionTok{ggplot}\NormalTok{(}\FunctionTok{subset}\NormalTok{(Expression\_allEnvironments\_allGeno, MUTATION.Expression }\SpecialCharTok{==} \StringTok{"REF"}\NormalTok{), }
                                   \FunctionTok{aes}\NormalTok{(}\AttributeTok{x =}\NormalTok{ ENVIRONMENT, }\AttributeTok{y =}\NormalTok{ YFP.MEDIAN.ADJUST\_driftcorrected, }\AttributeTok{color =}\NormalTok{ ENVIRONMENT)) }\SpecialCharTok{+} 
  \FunctionTok{geom\_point}\NormalTok{(}\AttributeTok{position =} \FunctionTok{position\_jitterdodge}\NormalTok{(}\AttributeTok{jitter.width =} \FloatTok{0.4}\NormalTok{), }\AttributeTok{alpha =} \FloatTok{0.25}\NormalTok{)  }\SpecialCharTok{+}
  \FunctionTok{geom\_violin}\NormalTok{(}\FunctionTok{aes}\NormalTok{(}\AttributeTok{fill =}\NormalTok{ ENVIRONMENT), }\AttributeTok{draw\_quantiles =} \FunctionTok{c}\NormalTok{(}\FloatTok{0.5}\NormalTok{), }
               \AttributeTok{position =} \FunctionTok{position\_dodge}\NormalTok{(}\AttributeTok{width =} \FloatTok{0.75}\NormalTok{), }\AttributeTok{width =} \FloatTok{0.5}\NormalTok{, }\AttributeTok{alpha =} \FloatTok{0.25}\NormalTok{, }\AttributeTok{col =} \StringTok{"Black"}\NormalTok{) }\SpecialCharTok{+}  \CommentTok{\# Adjust width as needed\#}
  \FunctionTok{labs}\NormalTok{(}\AttributeTok{x =} \StringTok{""}\NormalTok{, }\AttributeTok{y =} \StringTok{"YFP Expression"}\NormalTok{) }\SpecialCharTok{+} 
  \FunctionTok{theme\_bw}\NormalTok{() }\SpecialCharTok{+}   \FunctionTok{scale\_color\_manual}\NormalTok{(}\AttributeTok{values =}\NormalTok{ named\_colors) }\SpecialCharTok{+} \FunctionTok{scale\_fill\_manual}\NormalTok{(}\AttributeTok{values =}\NormalTok{ named\_colors) }\SpecialCharTok{+}
  \FunctionTok{theme}\NormalTok{(}\AttributeTok{panel.border =} \FunctionTok{element\_rect}\NormalTok{(}\AttributeTok{linetype =} \DecValTok{1}\NormalTok{, }\AttributeTok{fill =} \StringTok{"NA"}\NormalTok{),}
        \AttributeTok{plot.background =} \FunctionTok{element\_rect}\NormalTok{(}\AttributeTok{color =} \ConstantTok{NA}\NormalTok{),}\AttributeTok{legend.position=}\StringTok{"none"}\NormalTok{,}
        \AttributeTok{axis.text=}\FunctionTok{element\_text}\NormalTok{(}\AttributeTok{size=} \DecValTok{10}\NormalTok{), }\AttributeTok{axis.title=}\FunctionTok{element\_text}\NormalTok{(}\AttributeTok{size =}\DecValTok{14}\NormalTok{),}
        \AttributeTok{panel.grid.minor =} \FunctionTok{element\_blank}\NormalTok{(),}
        \AttributeTok{panel.grid.major =} \FunctionTok{element\_blank}\NormalTok{())}

\CommentTok{\# Plot of deletion effect among environments{-}{-}{-}{-}{-}{-}{-}{-}{-}{-}{-}{-}{-}{-}{-}{-}{-}{-}{-}{-}{-}{-}{-}{-}{-}{-}{-}{-}{-}}
\CommentTok{\#Keep only observations for which the MUTATION = "BARRY" and "TDH3.Deletion"  from the fitness\_summary data frame}
\NormalTok{fitness\_data\_DeletionEffect }\OtherTok{\textless{}{-}}\NormalTok{ FitnessData\_allEnvironments }\SpecialCharTok{\%\textgreater{}\%}
  \FunctionTok{filter}\NormalTok{(MUTATION.Fitness }\SpecialCharTok{\%in\%} \FunctionTok{c}\NormalTok{(}\StringTok{"REF"}\NormalTok{, }\StringTok{"TDH3.Deletion"}\NormalTok{))}

\NormalTok{plot\_DeletionEffect\_violin }\OtherTok{\textless{}{-}} \FunctionTok{ggplot}\NormalTok{(fitness\_data\_DeletionEffect, }\FunctionTok{aes}\NormalTok{(}\AttributeTok{x =}\NormalTok{ ENVIRONMENT, }\AttributeTok{y =}\NormalTok{ RelativeFitness, }
                                                               \AttributeTok{col =}\NormalTok{ ENVIRONMENT, }\AttributeTok{shape =}\NormalTok{ MUTATION.Fitness)) }\SpecialCharTok{+}
  \FunctionTok{geom\_hline}\NormalTok{(}\AttributeTok{yintercept =} \DecValTok{1}\NormalTok{, }\AttributeTok{linetype =} \StringTok{"dashed"}\NormalTok{, }\AttributeTok{color =} \StringTok{"black"}\NormalTok{) }\SpecialCharTok{+}
  \FunctionTok{scale\_y\_continuous}\NormalTok{(}\AttributeTok{breaks =} \FunctionTok{c}\NormalTok{(}\FloatTok{0.94}\NormalTok{, }\FloatTok{0.96}\NormalTok{, }\FloatTok{0.98}\NormalTok{, }\FloatTok{1.0}\NormalTok{)) }\SpecialCharTok{+}
  \FunctionTok{geom\_point}\NormalTok{(}\AttributeTok{position =} \FunctionTok{position\_jitterdodge}\NormalTok{(}\AttributeTok{jitter.width =} \FloatTok{0.4}\NormalTok{), }\AttributeTok{alpha =} \FloatTok{0.8}\NormalTok{)  }\SpecialCharTok{+}
  \FunctionTok{geom\_violin}\NormalTok{(}\AttributeTok{col =} \StringTok{"Black"}\NormalTok{, }\FunctionTok{aes}\NormalTok{(}\AttributeTok{fill =}\NormalTok{ ENVIRONMENT), }\AttributeTok{alpha =} \FloatTok{0.20}\NormalTok{,}\AttributeTok{draw\_quantiles =} \FunctionTok{c}\NormalTok{(}\FloatTok{0.5}\NormalTok{)) }\SpecialCharTok{+}
  \FunctionTok{labs}\NormalTok{(}\AttributeTok{x =} \StringTok{""}\NormalTok{, }\AttributeTok{y =} \StringTok{"Relative Fitness"}\NormalTok{) }\SpecialCharTok{+}
  \FunctionTok{theme\_bw}\NormalTok{() }\SpecialCharTok{+}
  \FunctionTok{scale\_color\_manual}\NormalTok{(}\AttributeTok{values =}\NormalTok{ named\_colors) }\SpecialCharTok{+}  
  \FunctionTok{scale\_fill\_manual}\NormalTok{(}\AttributeTok{values =}\NormalTok{ named\_colors) }\SpecialCharTok{+}  
  \FunctionTok{theme}\NormalTok{(}\AttributeTok{panel.grid.major =} \FunctionTok{element\_blank}\NormalTok{(), }\AttributeTok{panel.grid.minor =} \FunctionTok{element\_blank}\NormalTok{()}
\NormalTok{        )}


\NormalTok{plot\_DoublingTime}
\end{Highlighting}
\end{Shaded}

\includegraphics{ExpressionFitnessAnalysis_files/figure-latex/unnamed-chunk-2-1.pdf}

\begin{Shaded}
\begin{Highlighting}[]
\NormalTok{plot\_ExpressionReference\_violin}
\end{Highlighting}
\end{Shaded}

\includegraphics{ExpressionFitnessAnalysis_files/figure-latex/unnamed-chunk-2-2.pdf}

\begin{Shaded}
\begin{Highlighting}[]
\NormalTok{plot\_DeletionEffect\_violin}
\end{Highlighting}
\end{Shaded}

\includegraphics{ExpressionFitnessAnalysis_files/figure-latex/unnamed-chunk-2-3.pdf}

\begin{Shaded}
\begin{Highlighting}[]
\CommentTok{\# Save the plots}
\FunctionTok{ggsave}\NormalTok{(}\StringTok{"./ManuscriptFigures/Fig2\_plot\_ExpressionReference\_violin.pdf"}\NormalTok{, }\AttributeTok{plot =}\NormalTok{ plot\_ExpressionReference\_violin, }\AttributeTok{width =} \FloatTok{4.5}\NormalTok{, }\AttributeTok{height =} \DecValTok{4}\NormalTok{, }\AttributeTok{units =} \StringTok{"in"}\NormalTok{, }\AttributeTok{dpi =} \DecValTok{300}\NormalTok{)}
\FunctionTok{ggsave}\NormalTok{(}\StringTok{"./ManuscriptFigures/Fig2\_plot\_DoublingTime.pdf"}\NormalTok{, }\AttributeTok{plot =}\NormalTok{ plot\_DoublingTime, }\AttributeTok{width =} \FloatTok{3.0}\NormalTok{, }\AttributeTok{height =} \DecValTok{4}\NormalTok{)}
\FunctionTok{ggsave}\NormalTok{(}\StringTok{"./ManuscriptFigures/Fig2\_plot\_DeletionEffect\_violin.pdf"}\NormalTok{, }\AttributeTok{plot =}\NormalTok{ plot\_DeletionEffect\_violin, }\AttributeTok{width =} \DecValTok{3}\NormalTok{, }\AttributeTok{height =} \DecValTok{4}\NormalTok{)}
\end{Highlighting}
\end{Shaded}

\hypertarget{figure-3a-b-variation-in-gene-expression-among-environments}{%
\subsubsection{Figure 3A-B: Variation in gene expression among
environments}\label{figure-3a-b-variation-in-gene-expression-among-environments}}

The following code produces the panels in Figure 3A and Figure 3B. These
are the fitness curves in the different environment. The x-axis values
(expression levels) have all been normalized by the value of the
unmutated, reference strain in the glucose environment.

\begin{Shaded}
\begin{Highlighting}[]
\CommentTok{\#Take the ExpressionFitnessData\_allEnvironments. Classify genotypes based on the number of their mutations. Create a column called NumPromMutation.}
\CommentTok{\#The following strains have 1 mutation:Y1892,Y1896,Y1897,Y1898,Y1901,Y1903,Y1960,Y2528,Y2546,Y2566,Y2605,Y2651,Y2656}
\CommentTok{\# The following strains have 2 mutations: Y2455,Y2458,Y2460,Y2465,Y2469,Y2475,Y2478,Y2653,Y2836,Y2842 }
\CommentTok{\# The following strains have \textgreater{}2 mutations: Y2550,Y2812,Y2814,Y2816,Y2820,Y2824,Y2826,Y2828,Y2832,Y2834,Y2840}
\CommentTok{\# The following strains have CNVs of YFP: Y2671,Y2675,Y2676,Y3006,Y3010,Y3011,Y3012,Y3015,Y3019,Y3021,Y3022,Y3025,Y3028,Y3034,Y3040}

\NormalTok{ExpressionFitnessData\_allEnvironments\_numProm }\OtherTok{\textless{}{-}}\NormalTok{ ExpressionFitnessData\_allEnvironments }\SpecialCharTok{\%\textgreater{}\%}
  \FunctionTok{mutate}\NormalTok{(}\AttributeTok{NumPromMutation =} \FunctionTok{case\_when}\NormalTok{(}
\NormalTok{    STRAIN.Expression }\SpecialCharTok{\%in\%} \FunctionTok{c}\NormalTok{(}\StringTok{"Y1892"}\NormalTok{,}\StringTok{"Y1896"}\NormalTok{,}\StringTok{"Y1897"}\NormalTok{,}\StringTok{"Y1898"}\NormalTok{,}\StringTok{"Y1901"}\NormalTok{,}\StringTok{"Y1903"}\NormalTok{,}\StringTok{"Y1960"}\NormalTok{,}\StringTok{"Y2528"}\NormalTok{,}\StringTok{"Y2546"}\NormalTok{,}\StringTok{"Y2566"}\NormalTok{,}\StringTok{"Y2605"}\NormalTok{,}\StringTok{"Y2651"}\NormalTok{,}\StringTok{"Y2656"}\NormalTok{) }\SpecialCharTok{\textasciitilde{}} \StringTok{"1"}\NormalTok{,}
\NormalTok{    STRAIN.Expression }\SpecialCharTok{\%in\%} \FunctionTok{c}\NormalTok{(}\StringTok{"Y2455"}\NormalTok{,}\StringTok{"Y2458"}\NormalTok{,}\StringTok{"Y2460"}\NormalTok{,}\StringTok{"Y2465"}\NormalTok{,}\StringTok{"Y2469"}\NormalTok{,}\StringTok{"Y2475"}\NormalTok{,}\StringTok{"Y2478"}\NormalTok{,}\StringTok{"Y2653"}\NormalTok{,}\StringTok{"Y2836"}\NormalTok{,}\StringTok{"Y2842"}\NormalTok{) }\SpecialCharTok{\textasciitilde{}} \StringTok{"2"}\NormalTok{,}
\NormalTok{    STRAIN.Expression }\SpecialCharTok{\%in\%} \FunctionTok{c}\NormalTok{(}\StringTok{"Y2550"}\NormalTok{,}\StringTok{"Y2812"}\NormalTok{,}\StringTok{"Y2814"}\NormalTok{,}\StringTok{"Y2816"}\NormalTok{,}\StringTok{"Y2820"}\NormalTok{,}\StringTok{"Y2824"}\NormalTok{,}\StringTok{"Y2826"}\NormalTok{,}\StringTok{"Y2828"}\NormalTok{,}\StringTok{"Y2832"}\NormalTok{,}\StringTok{"Y2834"}\NormalTok{,}\StringTok{"Y2840"}\NormalTok{) }\SpecialCharTok{\textasciitilde{}} \StringTok{"\textgreater{} 2"}\NormalTok{,}
\NormalTok{    STRAIN.Expression }\SpecialCharTok{\%in\%} \FunctionTok{c}\NormalTok{(}\StringTok{"Y2671"}\NormalTok{,}\StringTok{"Y2675"}\NormalTok{,}\StringTok{"Y2676"}\NormalTok{,}\StringTok{"Y3006"}\NormalTok{,}\StringTok{"Y3010"}\NormalTok{,}\StringTok{"Y3011"}\NormalTok{,}\StringTok{"Y3012"}\NormalTok{,}\StringTok{"Y3015"}\NormalTok{,}\StringTok{"Y3019"}\NormalTok{,}\StringTok{"Y3021"}\NormalTok{,}\StringTok{"Y3022"}\NormalTok{,}\StringTok{"Y3025"}\NormalTok{,}\StringTok{"Y3028"}\NormalTok{,}\StringTok{"Y3034"}\NormalTok{,}\StringTok{"Y3040"}\NormalTok{) }\SpecialCharTok{\textasciitilde{}} \StringTok{"CNV"}\NormalTok{,}
\NormalTok{      )) }

\NormalTok{unmutated }\OtherTok{\textless{}{-}} \FunctionTok{c}\NormalTok{(}\StringTok{"REF"}\NormalTok{, }\StringTok{"S288c"}\NormalTok{, }\StringTok{"NEG"}\NormalTok{, }\StringTok{"DUP"}\NormalTok{)}

\NormalTok{ExpressionFitnessData\_allEnvironments\_numProm}\SpecialCharTok{$}\NormalTok{ENVIRONMENT }\OtherTok{\textless{}{-}} \FunctionTok{fct\_relevel}\NormalTok{(ExpressionFitnessData\_allEnvironments\_numProm}\SpecialCharTok{$}\NormalTok{ENVIRONMENT, }\FunctionTok{c}\NormalTok{(}\StringTok{"GLUCOSE"}\NormalTok{, }\StringTok{"GALACTOSE"}\NormalTok{, }\StringTok{"GLYCEROL"}\NormalTok{, }\StringTok{"ETHANOL"}\NormalTok{))}
\NormalTok{ExpressionFitnessData\_allEnvironments\_numProm}\SpecialCharTok{$}\NormalTok{NumPromMutation }\OtherTok{\textless{}{-}} \FunctionTok{fct\_relevel}\NormalTok{(ExpressionFitnessData\_allEnvironments\_numProm}\SpecialCharTok{$}\NormalTok{NumPromMutation, }
                                                              \FunctionTok{c}\NormalTok{(}\StringTok{"0"}\NormalTok{, }\StringTok{"1"}\NormalTok{, }\StringTok{"2"}\NormalTok{, }\StringTok{"\textgreater{} 2"}\NormalTok{, }\StringTok{"CNV"}\NormalTok{))}
\end{Highlighting}
\end{Shaded}

\begin{verbatim}
## Warning: 1 unknown level in `f`: 0
\end{verbatim}

\begin{Shaded}
\begin{Highlighting}[]
\DocumentationTok{\#\#\# The function for doing pairwise t{-}tests between observations for each strain in different environments}
\DocumentationTok{\#\#\# with bonferroni correction. }
\DocumentationTok{\#\#\# Test alternative approach {-}{-} using a function }
\NormalTok{do\_t\_test }\OtherTok{\textless{}{-}} \ControlFlowTok{function}\NormalTok{(df, env1, env2, }\AttributeTok{adjustment\_method =} \StringTok{"bonferroni"}\NormalTok{) \{}
  \CommentTok{\# Subset the data for the two environments}
\NormalTok{  data\_subset }\OtherTok{\textless{}{-}}\NormalTok{ df }\SpecialCharTok{\%\textgreater{}\%}
    \FunctionTok{filter}\NormalTok{(ENVIRONMENT }\SpecialCharTok{\%in\%} \FunctionTok{c}\NormalTok{(env1, env2)) }\SpecialCharTok{\%\textgreater{}\%}
    \FunctionTok{select}\NormalTok{(STRAIN.Expression, ENVIRONMENT, YFP.MEDIAN.RELATIVE) }
  
  \CommentTok{\# Perform the t{-}test within each strain and combination of environments}
\NormalTok{  t\_test\_results }\OtherTok{\textless{}{-}}\NormalTok{ data\_subset }\SpecialCharTok{\%\textgreater{}\%}
    \FunctionTok{group\_by}\NormalTok{(STRAIN.Expression) }\SpecialCharTok{\%\textgreater{}\%}
    \FunctionTok{do}\NormalTok{(}\FunctionTok{tidy}\NormalTok{(}\FunctionTok{t.test}\NormalTok{(YFP.MEDIAN.RELATIVE }\SpecialCharTok{\textasciitilde{}}\NormalTok{ ENVIRONMENT, }\AttributeTok{data =}\NormalTok{ .)))}
  
  \CommentTok{\# Extract the p{-}values and perform p{-}value adjustment}
\NormalTok{  p\_values }\OtherTok{\textless{}{-}}\NormalTok{ t\_test\_results }\SpecialCharTok{\%\textgreater{}\%}
    \FunctionTok{select}\NormalTok{(STRAIN.Expression, p.value) }\SpecialCharTok{\%\textgreater{}\%}
    \FunctionTok{mutate}\NormalTok{(}\AttributeTok{p.value =} \FunctionTok{p.adjust}\NormalTok{(p.value, }\AttributeTok{method =}\NormalTok{ adjustment\_method)) }\SpecialCharTok{\%\textgreater{}\%}
    \FunctionTok{distinct}\NormalTok{()}
  
  \CommentTok{\# Summarize the data and add the p{-}values}
\NormalTok{  summary\_data }\OtherTok{\textless{}{-}}\NormalTok{ data\_subset }\SpecialCharTok{\%\textgreater{}\%}
    \FunctionTok{left\_join}\NormalTok{(p\_values, }\AttributeTok{by =} \StringTok{"STRAIN.Expression"}\NormalTok{) }\SpecialCharTok{\%\textgreater{}\%}
    \FunctionTok{group\_by}\NormalTok{(STRAIN.Expression, ENVIRONMENT) }\SpecialCharTok{\%\textgreater{}\%}
    \FunctionTok{summarise}\NormalTok{(}\AttributeTok{YFP.MEDIAN.RELATIVE.MEAN =} \FunctionTok{mean}\NormalTok{(YFP.MEDIAN.RELATIVE), }
              \AttributeTok{YFP.MEDIAN.RELATIVE.SD =} \FunctionTok{sd}\NormalTok{(YFP.MEDIAN.RELATIVE),}
              \AttributeTok{n =} \FunctionTok{n}\NormalTok{(),}
              \AttributeTok{p.adjusted =} \FunctionTok{first}\NormalTok{(p.value)) }\SpecialCharTok{\%\textgreater{}\%}
    \FunctionTok{mutate}\NormalTok{(}\AttributeTok{comp =} \FunctionTok{paste}\NormalTok{(env1, }\StringTok{"vs"}\NormalTok{, env2),}
           \AttributeTok{significance =} \FunctionTok{if\_else}\NormalTok{(p.adjusted }\SpecialCharTok{\textless{}} \FloatTok{0.05}\NormalTok{, }\StringTok{"Significant"}\NormalTok{, }\StringTok{"Not significant"}\NormalTok{))}
  
  \FunctionTok{return}\NormalTok{(summary\_data)}
\NormalTok{\}}

\DocumentationTok{\#\# The pairwise environments to pass on to the function }
\NormalTok{env\_pairs }\OtherTok{\textless{}{-}} \FunctionTok{list}\NormalTok{(}
  \FunctionTok{c}\NormalTok{(}\StringTok{"GLUCOSE"}\NormalTok{, }\StringTok{"GALACTOSE"}\NormalTok{),}
  \FunctionTok{c}\NormalTok{(}\StringTok{"GLUCOSE"}\NormalTok{, }\StringTok{"GLYCEROL"}\NormalTok{),}
  \FunctionTok{c}\NormalTok{(}\StringTok{"GLUCOSE"}\NormalTok{, }\StringTok{"ETHANOL"}\NormalTok{),}
  \FunctionTok{c}\NormalTok{(}\StringTok{"GALACTOSE"}\NormalTok{, }\StringTok{"GLYCEROL"}\NormalTok{),}
  \FunctionTok{c}\NormalTok{(}\StringTok{"GALACTOSE"}\NormalTok{, }\StringTok{"ETHANOL"}\NormalTok{),}
  \FunctionTok{c}\NormalTok{(}\StringTok{"GLYCEROL"}\NormalTok{, }\StringTok{"ETHANOL"}\NormalTok{)}
\NormalTok{)}

\CommentTok{\# For all of the data}
\NormalTok{all\_results }\OtherTok{\textless{}{-}} \FunctionTok{map\_dfr}\NormalTok{(env\_pairs, }\SpecialCharTok{\textasciitilde{}}\FunctionTok{do\_t\_test}\NormalTok{(Expression\_allEnvironments\_allGeno, .x[}\DecValTok{1}\NormalTok{], .x[}\DecValTok{2}\NormalTok{]))}
\end{Highlighting}
\end{Shaded}

\begin{verbatim}
## `summarise()` has grouped output by 'STRAIN.Expression'. You can override using
## the `.groups` argument.
## `summarise()` has grouped output by 'STRAIN.Expression'. You can override using
## the `.groups` argument.
## `summarise()` has grouped output by 'STRAIN.Expression'. You can override using
## the `.groups` argument.
## `summarise()` has grouped output by 'STRAIN.Expression'. You can override using
## the `.groups` argument.
## `summarise()` has grouped output by 'STRAIN.Expression'. You can override using
## the `.groups` argument.
## `summarise()` has grouped output by 'STRAIN.Expression'. You can override using
## the `.groups` argument.
\end{verbatim}

\begin{Shaded}
\begin{Highlighting}[]
\CommentTok{\# Split into four data frames based on the comparison (in comp column). For each of the four datasets,}
\CommentTok{\# convert the data into wide format such that there is a YFP.MEDIAN.RELATIVE.MEAN\_GLU, }
\CommentTok{\# YFP.MEDIAN.RELATIVE.SD\_GLU, YFP.MEDIAN.RELATIVE.MEAN\_GAL, YFP.MEDIAN.RELATIVE.SD\_GAL, etc.Keep the signifiance column,}
\CommentTok{\# n, and p.adjusted column}

\NormalTok{data\_GLU\_GAL }\OtherTok{\textless{}{-}}\NormalTok{ all\_results }\SpecialCharTok{\%\textgreater{}\%}
  \FunctionTok{filter}\NormalTok{(comp }\SpecialCharTok{==} \StringTok{"GLUCOSE vs GALACTOSE"}\NormalTok{) }\SpecialCharTok{\%\textgreater{}\%}
  \FunctionTok{pivot\_wider}\NormalTok{(}\AttributeTok{names\_from =}\NormalTok{ ENVIRONMENT, }\AttributeTok{values\_from =} \FunctionTok{c}\NormalTok{(YFP.MEDIAN.RELATIVE.MEAN, YFP.MEDIAN.RELATIVE.SD))}

\NormalTok{data\_GLU\_GLY }\OtherTok{\textless{}{-}}\NormalTok{ all\_results }\SpecialCharTok{\%\textgreater{}\%}
  \FunctionTok{filter}\NormalTok{(comp }\SpecialCharTok{==} \StringTok{"GLUCOSE vs GLYCEROL"}\NormalTok{) }\SpecialCharTok{\%\textgreater{}\%}
  \FunctionTok{pivot\_wider}\NormalTok{(}\AttributeTok{names\_from =}\NormalTok{ ENVIRONMENT, }\AttributeTok{values\_from =} \FunctionTok{c}\NormalTok{(YFP.MEDIAN.RELATIVE.MEAN, YFP.MEDIAN.RELATIVE.SD))}

\NormalTok{data\_GLU\_ETH }\OtherTok{\textless{}{-}}\NormalTok{ all\_results }\SpecialCharTok{\%\textgreater{}\%}
  \FunctionTok{filter}\NormalTok{(comp }\SpecialCharTok{==} \StringTok{"GLUCOSE vs ETHANOL"}\NormalTok{) }\SpecialCharTok{\%\textgreater{}\%}
  \FunctionTok{pivot\_wider}\NormalTok{(}\AttributeTok{names\_from =}\NormalTok{ ENVIRONMENT, }\AttributeTok{values\_from =} \FunctionTok{c}\NormalTok{(YFP.MEDIAN.RELATIVE.MEAN, YFP.MEDIAN.RELATIVE.SD))}

\NormalTok{data\_GAL\_GLY }\OtherTok{\textless{}{-}}\NormalTok{ all\_results }\SpecialCharTok{\%\textgreater{}\%}
  \FunctionTok{filter}\NormalTok{(comp }\SpecialCharTok{==} \StringTok{"GALACTOSE vs GLYCEROL"}\NormalTok{) }\SpecialCharTok{\%\textgreater{}\%}
  \FunctionTok{pivot\_wider}\NormalTok{(}\AttributeTok{names\_from =}\NormalTok{ ENVIRONMENT, }\AttributeTok{values\_from =} \FunctionTok{c}\NormalTok{(YFP.MEDIAN.RELATIVE.MEAN, YFP.MEDIAN.RELATIVE.SD))}

\NormalTok{data\_GAL\_ETH }\OtherTok{\textless{}{-}}\NormalTok{ all\_results }\SpecialCharTok{\%\textgreater{}\%}
  \FunctionTok{filter}\NormalTok{(comp }\SpecialCharTok{==} \StringTok{"GALACTOSE vs ETHANOL"}\NormalTok{) }\SpecialCharTok{\%\textgreater{}\%}
  \FunctionTok{pivot\_wider}\NormalTok{(}\AttributeTok{names\_from =}\NormalTok{ ENVIRONMENT, }\AttributeTok{values\_from =} \FunctionTok{c}\NormalTok{(YFP.MEDIAN.RELATIVE.MEAN, YFP.MEDIAN.RELATIVE.SD))}

\NormalTok{data\_GLY\_ETH }\OtherTok{\textless{}{-}}\NormalTok{ all\_results }\SpecialCharTok{\%\textgreater{}\%}
  \FunctionTok{filter}\NormalTok{(comp }\SpecialCharTok{==} \StringTok{"GLYCEROL vs ETHANOL"}\NormalTok{) }\SpecialCharTok{\%\textgreater{}\%}
  \FunctionTok{pivot\_wider}\NormalTok{(}\AttributeTok{names\_from =}\NormalTok{ ENVIRONMENT, }\AttributeTok{values\_from =} \FunctionTok{c}\NormalTok{(YFP.MEDIAN.RELATIVE.MEAN, YFP.MEDIAN.RELATIVE.SD))}

\DocumentationTok{\#\#\#\#\# Plots{-}{-}{-}{-}{-}{-}{-}{-}{-}}
\NormalTok{named\_colors }\OtherTok{\textless{}{-}} \FunctionTok{c}\NormalTok{(}\StringTok{"GLUCOSE"} \OtherTok{=} \StringTok{"\#0072B2"}\NormalTok{, }\StringTok{"GALACTOSE"} \OtherTok{=} \StringTok{"\#009E73"}\NormalTok{, }\StringTok{"GLYCEROL"} \OtherTok{=} \StringTok{"\#CC79A7"}\NormalTok{, }\StringTok{"ETHANOL"} \OtherTok{=} \StringTok{"\#D55E00"}\NormalTok{)}


\CommentTok{\# Generate scatter plots for each environment pair}
\NormalTok{plot\_GLUvGAL\_sig }\OtherTok{\textless{}{-}} \FunctionTok{ggplot}\NormalTok{(data\_GLU\_GAL, }\FunctionTok{aes}\NormalTok{(}\AttributeTok{x =}\NormalTok{ YFP.MEDIAN.RELATIVE.MEAN\_GLUCOSE, }\AttributeTok{y =}\NormalTok{ YFP.MEDIAN.RELATIVE.MEAN\_GALACTOSE, }\AttributeTok{color =}\NormalTok{ significance)) }\SpecialCharTok{+}
  \FunctionTok{geom\_point}\NormalTok{(}\AttributeTok{size =} \DecValTok{2}\NormalTok{, }\AttributeTok{alpha =} \FloatTok{0.5}\NormalTok{) }\SpecialCharTok{+} \CommentTok{\#add standard error bars}
  \FunctionTok{geom\_errorbar}\NormalTok{(}\FunctionTok{aes}\NormalTok{(}\AttributeTok{ymin =}\NormalTok{ YFP.MEDIAN.RELATIVE.MEAN\_GALACTOSE }\SpecialCharTok{{-}}\NormalTok{ YFP.MEDIAN.RELATIVE.SD\_GALACTOSE, }
                    \AttributeTok{ymax =}\NormalTok{ YFP.MEDIAN.RELATIVE.MEAN\_GALACTOSE }\SpecialCharTok{+}\NormalTok{ YFP.MEDIAN.RELATIVE.SD\_GALACTOSE), }
                \AttributeTok{width =} \FloatTok{0.0}\NormalTok{) }\SpecialCharTok{+} \CommentTok{\#add horizontal error bars}
  \FunctionTok{geom\_errorbarh}\NormalTok{(}\FunctionTok{aes}\NormalTok{(}\AttributeTok{xmin =}\NormalTok{ YFP.MEDIAN.RELATIVE.MEAN\_GLUCOSE }\SpecialCharTok{{-}}\NormalTok{ YFP.MEDIAN.RELATIVE.SD\_GLUCOSE, }
                     \AttributeTok{xmax =}\NormalTok{ YFP.MEDIAN.RELATIVE.MEAN\_GLUCOSE }\SpecialCharTok{+}\NormalTok{ YFP.MEDIAN.RELATIVE.SD\_GLUCOSE), }
                 \AttributeTok{height =} \FloatTok{0.0}\NormalTok{) }\SpecialCharTok{+}
  \FunctionTok{geom\_abline}\NormalTok{(}\AttributeTok{intercept =} \DecValTok{0}\NormalTok{, }\AttributeTok{slope =} \DecValTok{1}\NormalTok{, }\AttributeTok{color =} \StringTok{"black"}\NormalTok{, }\AttributeTok{linetype =} \StringTok{"dashed"}\NormalTok{, }\AttributeTok{alpha =} \FloatTok{0.2}\NormalTok{) }\SpecialCharTok{+}
  \FunctionTok{geom\_hline}\NormalTok{(}\AttributeTok{yintercept =} \DecValTok{1}\NormalTok{, }\AttributeTok{color =} \StringTok{"grey"}\NormalTok{) }\SpecialCharTok{+} \FunctionTok{geom\_vline}\NormalTok{(}\AttributeTok{xintercept =} \DecValTok{1}\NormalTok{, }\AttributeTok{color =} \StringTok{"grey"}\NormalTok{) }\SpecialCharTok{+}
  \FunctionTok{xlim}\NormalTok{(}\DecValTok{0}\NormalTok{, }\FloatTok{2.1}\NormalTok{) }\SpecialCharTok{+} \FunctionTok{ylim}\NormalTok{(}\DecValTok{0}\NormalTok{, }\FloatTok{2.1}\NormalTok{) }\SpecialCharTok{+}
  \FunctionTok{labs}\NormalTok{(}\AttributeTok{x =} \StringTok{"Relative Mean Expression in Glucose"}\NormalTok{, }\AttributeTok{y =} \StringTok{"Relative Mean Expression in Galactose"}\NormalTok{) }\SpecialCharTok{+} 
  \FunctionTok{scale\_color\_manual}\NormalTok{(}\AttributeTok{values =} \FunctionTok{c}\NormalTok{(}\StringTok{"Significant"} \OtherTok{=} \StringTok{"red"}\NormalTok{, }\StringTok{"Not significant"} \OtherTok{=} \StringTok{"black"}\NormalTok{)) }\SpecialCharTok{+}
  \FunctionTok{theme\_bw}\NormalTok{() }\SpecialCharTok{+}
  \FunctionTok{theme}\NormalTok{(}\AttributeTok{legend.position =} \StringTok{"bottom"}\NormalTok{,}
        \AttributeTok{axis.title =} \FunctionTok{element\_text}\NormalTok{(}\AttributeTok{size =} \DecValTok{16}\NormalTok{),}
        \AttributeTok{axis.text.x =} \FunctionTok{element\_text}\NormalTok{(}\AttributeTok{hjust =} \DecValTok{1}\NormalTok{),}
        \AttributeTok{panel.grid.minor =} \FunctionTok{element\_blank}\NormalTok{(),}
        \AttributeTok{panel.grid.major =} \FunctionTok{element\_blank}\NormalTok{())}

\DocumentationTok{\#\#Make the same plot as above for the other environment pairs}
\NormalTok{plot\_GLUvGLY\_sig }\OtherTok{\textless{}{-}}\FunctionTok{ggplot}\NormalTok{(data\_GLU\_GLY, }\FunctionTok{aes}\NormalTok{(}\AttributeTok{x =}\NormalTok{ YFP.MEDIAN.RELATIVE.MEAN\_GLUCOSE, }\AttributeTok{y =}\NormalTok{ YFP.MEDIAN.RELATIVE.MEAN\_GLYCEROL, }\AttributeTok{color =}\NormalTok{ significance)) }\SpecialCharTok{+}
  \FunctionTok{geom\_point}\NormalTok{(}\AttributeTok{size =} \DecValTok{2}\NormalTok{, }\AttributeTok{alpha =} \FloatTok{0.5}\NormalTok{) }\SpecialCharTok{+} \CommentTok{\#add standard error bars}
  \FunctionTok{geom\_errorbar}\NormalTok{(}\FunctionTok{aes}\NormalTok{(}\AttributeTok{ymin =}\NormalTok{ YFP.MEDIAN.RELATIVE.MEAN\_GLYCEROL }\SpecialCharTok{{-}}\NormalTok{ YFP.MEDIAN.RELATIVE.SD\_GLYCEROL, }
                    \AttributeTok{ymax =}\NormalTok{ YFP.MEDIAN.RELATIVE.MEAN\_GLYCEROL }\SpecialCharTok{+}\NormalTok{ YFP.MEDIAN.RELATIVE.SD\_GLYCEROL), }
                \AttributeTok{width =} \FloatTok{0.0}\NormalTok{) }\SpecialCharTok{+} \CommentTok{\#add horizontal error bars}
  \FunctionTok{geom\_errorbarh}\NormalTok{(}\FunctionTok{aes}\NormalTok{(}\AttributeTok{xmin =}\NormalTok{ YFP.MEDIAN.RELATIVE.MEAN\_GLUCOSE }\SpecialCharTok{{-}}\NormalTok{ YFP.MEDIAN.RELATIVE.SD\_GLUCOSE, }
                     \AttributeTok{xmax =}\NormalTok{ YFP.MEDIAN.RELATIVE.MEAN\_GLUCOSE }\SpecialCharTok{+}\NormalTok{ YFP.MEDIAN.RELATIVE.SD\_GLUCOSE), }
                 \AttributeTok{height =} \FloatTok{0.0}\NormalTok{) }\SpecialCharTok{+}
  \FunctionTok{geom\_abline}\NormalTok{(}\AttributeTok{intercept =} \DecValTok{0}\NormalTok{, }\AttributeTok{slope =} \DecValTok{1}\NormalTok{, }\AttributeTok{color =} \StringTok{"black"}\NormalTok{, }\AttributeTok{linetype =} \StringTok{"dashed"}\NormalTok{, }\AttributeTok{alpha =} \FloatTok{0.2}\NormalTok{) }\SpecialCharTok{+}
  \FunctionTok{geom\_hline}\NormalTok{(}\AttributeTok{yintercept =} \DecValTok{1}\NormalTok{, }\AttributeTok{color =} \StringTok{"grey"}\NormalTok{) }\SpecialCharTok{+} \FunctionTok{geom\_vline}\NormalTok{(}\AttributeTok{xintercept =} \DecValTok{1}\NormalTok{, }\AttributeTok{color =} \StringTok{"grey"}\NormalTok{) }\SpecialCharTok{+}
  \FunctionTok{xlim}\NormalTok{(}\DecValTok{0}\NormalTok{, }\FloatTok{2.1}\NormalTok{) }\SpecialCharTok{+} \FunctionTok{ylim}\NormalTok{(}\DecValTok{0}\NormalTok{, }\FloatTok{2.1}\NormalTok{) }\SpecialCharTok{+}
  \FunctionTok{labs}\NormalTok{(}\AttributeTok{x =} \StringTok{"Relative Mean Expression in Glucose"}\NormalTok{, }\AttributeTok{y =} \StringTok{"Relative Mean Expression in Glycerol"}\NormalTok{) }\SpecialCharTok{+} 
  \FunctionTok{scale\_color\_manual}\NormalTok{(}\AttributeTok{values =} \FunctionTok{c}\NormalTok{(}\StringTok{"Significant"} \OtherTok{=} \StringTok{"red"}\NormalTok{, }\StringTok{"Not significant"} \OtherTok{=} \StringTok{"black"}\NormalTok{)) }\SpecialCharTok{+}
  \FunctionTok{theme\_bw}\NormalTok{() }\SpecialCharTok{+}
  \FunctionTok{theme}\NormalTok{(}\AttributeTok{legend.position =} \StringTok{"bottom"}\NormalTok{,}
        \AttributeTok{axis.title =} \FunctionTok{element\_text}\NormalTok{(}\AttributeTok{size =} \DecValTok{16}\NormalTok{),}
        \AttributeTok{axis.text.x =} \FunctionTok{element\_text}\NormalTok{(}\AttributeTok{hjust =} \DecValTok{1}\NormalTok{),}
        \AttributeTok{panel.grid.minor =} \FunctionTok{element\_blank}\NormalTok{(),}
        \AttributeTok{panel.grid.major =} \FunctionTok{element\_blank}\NormalTok{())}

\DocumentationTok{\#\#\#}
\NormalTok{plot\_GLUvETH\_sig }\OtherTok{\textless{}{-}} \FunctionTok{ggplot}\NormalTok{(data\_GLU\_ETH, }\FunctionTok{aes}\NormalTok{(}\AttributeTok{x =}\NormalTok{ YFP.MEDIAN.RELATIVE.MEAN\_GLUCOSE, }\AttributeTok{y =}\NormalTok{ YFP.MEDIAN.RELATIVE.MEAN\_ETHANOL, }\AttributeTok{color =}\NormalTok{ significance)) }\SpecialCharTok{+}
  \FunctionTok{geom\_point}\NormalTok{(}\AttributeTok{size =} \DecValTok{2}\NormalTok{, }\AttributeTok{alpha =} \FloatTok{0.5}\NormalTok{) }\SpecialCharTok{+} \CommentTok{\#add standard error bars}
  \FunctionTok{geom\_errorbar}\NormalTok{(}\FunctionTok{aes}\NormalTok{(}\AttributeTok{ymin =}\NormalTok{ YFP.MEDIAN.RELATIVE.MEAN\_ETHANOL }\SpecialCharTok{{-}}\NormalTok{ YFP.MEDIAN.RELATIVE.SD\_ETHANOL, }
                    \AttributeTok{ymax =}\NormalTok{ YFP.MEDIAN.RELATIVE.MEAN\_ETHANOL }\SpecialCharTok{+}\NormalTok{ YFP.MEDIAN.RELATIVE.SD\_ETHANOL), }
                \AttributeTok{width =} \FloatTok{0.0}\NormalTok{) }\SpecialCharTok{+} \CommentTok{\#add horizontal error bars}
  \FunctionTok{geom\_errorbarh}\NormalTok{(}\FunctionTok{aes}\NormalTok{(}\AttributeTok{xmin =}\NormalTok{ YFP.MEDIAN.RELATIVE.MEAN\_GLUCOSE }\SpecialCharTok{{-}}\NormalTok{ YFP.MEDIAN.RELATIVE.SD\_GLUCOSE, }
                     \AttributeTok{xmax =}\NormalTok{ YFP.MEDIAN.RELATIVE.MEAN\_GLUCOSE }\SpecialCharTok{+}\NormalTok{ YFP.MEDIAN.RELATIVE.SD\_GLUCOSE), }
                 \AttributeTok{height =} \FloatTok{0.0}\NormalTok{) }\SpecialCharTok{+}
  \FunctionTok{geom\_abline}\NormalTok{(}\AttributeTok{intercept =} \DecValTok{0}\NormalTok{, }\AttributeTok{slope =} \DecValTok{1}\NormalTok{, }\AttributeTok{color =} \StringTok{"black"}\NormalTok{, }\AttributeTok{linetype =} \StringTok{"dashed"}\NormalTok{, }\AttributeTok{alpha =} \FloatTok{0.2}\NormalTok{) }\SpecialCharTok{+}
  \FunctionTok{geom\_hline}\NormalTok{(}\AttributeTok{yintercept =} \DecValTok{1}\NormalTok{, }\AttributeTok{color =} \StringTok{"grey"}\NormalTok{) }\SpecialCharTok{+} \FunctionTok{geom\_vline}\NormalTok{(}\AttributeTok{xintercept =} \DecValTok{1}\NormalTok{, }\AttributeTok{color =} \StringTok{"grey"}\NormalTok{) }\SpecialCharTok{+}
  \FunctionTok{xlim}\NormalTok{(}\DecValTok{0}\NormalTok{, }\FloatTok{2.1}\NormalTok{) }\SpecialCharTok{+} \FunctionTok{ylim}\NormalTok{(}\DecValTok{0}\NormalTok{, }\FloatTok{2.1}\NormalTok{) }\SpecialCharTok{+}
  \FunctionTok{labs}\NormalTok{(}\AttributeTok{x =} \StringTok{"Relative Mean Expression in Glucose"}\NormalTok{, }\AttributeTok{y =} \StringTok{"Relative Mean Expression in Ethanol"}\NormalTok{) }\SpecialCharTok{+} 
  \FunctionTok{scale\_color\_manual}\NormalTok{(}\AttributeTok{values =} \FunctionTok{c}\NormalTok{(}\StringTok{"Significant"} \OtherTok{=} \StringTok{"red"}\NormalTok{, }\StringTok{"Not significant"} \OtherTok{=} \StringTok{"black"}\NormalTok{)) }\SpecialCharTok{+}
  \FunctionTok{theme\_bw}\NormalTok{() }\SpecialCharTok{+}
  \FunctionTok{theme}\NormalTok{(}\AttributeTok{legend.position =} \StringTok{"bottom"}\NormalTok{,}
        \AttributeTok{axis.title =} \FunctionTok{element\_text}\NormalTok{(}\AttributeTok{size =} \DecValTok{16}\NormalTok{),}
        \AttributeTok{axis.text.x =} \FunctionTok{element\_text}\NormalTok{(}\AttributeTok{hjust =} \DecValTok{1}\NormalTok{),}
        \AttributeTok{panel.grid.minor =} \FunctionTok{element\_blank}\NormalTok{(),}
        \AttributeTok{panel.grid.major =} \FunctionTok{element\_blank}\NormalTok{())}

\DocumentationTok{\#\#\#}
\NormalTok{plot\_GALvGLY\_sig }\OtherTok{\textless{}{-}} \FunctionTok{ggplot}\NormalTok{(data\_GAL\_GLY, }\FunctionTok{aes}\NormalTok{(}\AttributeTok{x =}\NormalTok{ YFP.MEDIAN.RELATIVE.MEAN\_GALACTOSE, }\AttributeTok{y =}\NormalTok{ YFP.MEDIAN.RELATIVE.MEAN\_GLYCEROL, }\AttributeTok{color =}\NormalTok{ significance)) }\SpecialCharTok{+}
  \FunctionTok{geom\_point}\NormalTok{(}\AttributeTok{size =} \DecValTok{2}\NormalTok{, }\AttributeTok{alpha =} \FloatTok{0.5}\NormalTok{) }\SpecialCharTok{+} \CommentTok{\#add standard error bars}
  \FunctionTok{geom\_errorbar}\NormalTok{(}\FunctionTok{aes}\NormalTok{(}\AttributeTok{ymin =}\NormalTok{ YFP.MEDIAN.RELATIVE.MEAN\_GLYCEROL }\SpecialCharTok{{-}}\NormalTok{ YFP.MEDIAN.RELATIVE.SD\_GLYCEROL, }
                    \AttributeTok{ymax =}\NormalTok{ YFP.MEDIAN.RELATIVE.MEAN\_GLYCEROL }\SpecialCharTok{+}\NormalTok{ YFP.MEDIAN.RELATIVE.SD\_GLYCEROL), }
                \AttributeTok{width =} \FloatTok{0.0}\NormalTok{) }\SpecialCharTok{+} \CommentTok{\#add horizontal error bars}
  \FunctionTok{geom\_errorbarh}\NormalTok{(}\FunctionTok{aes}\NormalTok{(}\AttributeTok{xmin =}\NormalTok{ YFP.MEDIAN.RELATIVE.MEAN\_GALACTOSE }\SpecialCharTok{{-}}\NormalTok{ YFP.MEDIAN.RELATIVE.SD\_GALACTOSE, }
                     \AttributeTok{xmax =}\NormalTok{ YFP.MEDIAN.RELATIVE.MEAN\_GALACTOSE }\SpecialCharTok{+}\NormalTok{ YFP.MEDIAN.RELATIVE.SD\_GALACTOSE), }
                 \AttributeTok{height =} \FloatTok{0.0}\NormalTok{) }\SpecialCharTok{+}
  \FunctionTok{geom\_abline}\NormalTok{(}\AttributeTok{intercept =} \DecValTok{0}\NormalTok{, }\AttributeTok{slope =} \DecValTok{1}\NormalTok{, }\AttributeTok{color =} \StringTok{"black"}\NormalTok{, }\AttributeTok{linetype =} \StringTok{"dashed"}\NormalTok{, }\AttributeTok{alpha =} \FloatTok{0.2}\NormalTok{) }\SpecialCharTok{+}
  \FunctionTok{geom\_hline}\NormalTok{(}\AttributeTok{yintercept =} \DecValTok{1}\NormalTok{, }\AttributeTok{color =} \StringTok{"grey"}\NormalTok{) }\SpecialCharTok{+} \FunctionTok{geom\_vline}\NormalTok{(}\AttributeTok{xintercept =} \DecValTok{1}\NormalTok{, }\AttributeTok{color =} \StringTok{"grey"}\NormalTok{) }\SpecialCharTok{+}
  \FunctionTok{xlim}\NormalTok{(}\DecValTok{0}\NormalTok{, }\FloatTok{2.1}\NormalTok{) }\SpecialCharTok{+} \FunctionTok{ylim}\NormalTok{(}\DecValTok{0}\NormalTok{, }\FloatTok{2.1}\NormalTok{) }\SpecialCharTok{+}
  \FunctionTok{labs}\NormalTok{(}\AttributeTok{x =} \StringTok{"Relative Mean Expression in Galactose"}\NormalTok{, }\AttributeTok{y =} \StringTok{"Relative Mean Expression in Glycerol"}\NormalTok{) }\SpecialCharTok{+} 
  \FunctionTok{scale\_color\_manual}\NormalTok{(}\AttributeTok{values =} \FunctionTok{c}\NormalTok{(}\StringTok{"Significant"} \OtherTok{=} \StringTok{"red"}\NormalTok{, }\StringTok{"Not significant"} \OtherTok{=} \StringTok{"black"}\NormalTok{)) }\SpecialCharTok{+}
  \FunctionTok{theme\_bw}\NormalTok{() }\SpecialCharTok{+}
  \FunctionTok{theme}\NormalTok{(}\AttributeTok{legend.position =} \StringTok{"bottom"}\NormalTok{,}
        \AttributeTok{axis.title =} \FunctionTok{element\_text}\NormalTok{(}\AttributeTok{size =} \DecValTok{16}\NormalTok{),}
        \AttributeTok{axis.text.x =} \FunctionTok{element\_text}\NormalTok{(}\AttributeTok{hjust =} \DecValTok{1}\NormalTok{),}
        \AttributeTok{panel.grid.minor =} \FunctionTok{element\_blank}\NormalTok{(),}
        \AttributeTok{panel.grid.major =} \FunctionTok{element\_blank}\NormalTok{())}

\DocumentationTok{\#\#\#}
\NormalTok{plot\_GALvETH\_sig }\OtherTok{\textless{}{-}} \FunctionTok{ggplot}\NormalTok{(data\_GAL\_ETH, }\FunctionTok{aes}\NormalTok{(}\AttributeTok{x =}\NormalTok{ YFP.MEDIAN.RELATIVE.MEAN\_GALACTOSE, }\AttributeTok{y =}\NormalTok{ YFP.MEDIAN.RELATIVE.MEAN\_ETHANOL, }\AttributeTok{color =}\NormalTok{ significance)) }\SpecialCharTok{+}
  \FunctionTok{geom\_point}\NormalTok{(}\AttributeTok{size =} \DecValTok{2}\NormalTok{, }\AttributeTok{alpha =} \FloatTok{0.5}\NormalTok{) }\SpecialCharTok{+} \CommentTok{\#add standard error bars}
  \FunctionTok{geom\_errorbar}\NormalTok{(}\FunctionTok{aes}\NormalTok{(}\AttributeTok{ymin =}\NormalTok{ YFP.MEDIAN.RELATIVE.MEAN\_ETHANOL }\SpecialCharTok{{-}}\NormalTok{ YFP.MEDIAN.RELATIVE.SD\_ETHANOL, }
                    \AttributeTok{ymax =}\NormalTok{ YFP.MEDIAN.RELATIVE.MEAN\_ETHANOL }\SpecialCharTok{+}\NormalTok{ YFP.MEDIAN.RELATIVE.SD\_ETHANOL), }
                \AttributeTok{width =} \FloatTok{0.0}\NormalTok{) }\SpecialCharTok{+} \CommentTok{\#add horizontal error bars}
  \FunctionTok{geom\_errorbarh}\NormalTok{(}\FunctionTok{aes}\NormalTok{(}\AttributeTok{xmin =}\NormalTok{ YFP.MEDIAN.RELATIVE.MEAN\_GALACTOSE }\SpecialCharTok{{-}}\NormalTok{ YFP.MEDIAN.RELATIVE.SD\_GALACTOSE, }
                     \AttributeTok{xmax =}\NormalTok{ YFP.MEDIAN.RELATIVE.MEAN\_GALACTOSE }\SpecialCharTok{+}\NormalTok{ YFP.MEDIAN.RELATIVE.SD\_GALACTOSE), }
                 \AttributeTok{height =} \FloatTok{0.0}\NormalTok{) }\SpecialCharTok{+}
  \FunctionTok{geom\_abline}\NormalTok{(}\AttributeTok{intercept =} \DecValTok{0}\NormalTok{, }\AttributeTok{slope =} \DecValTok{1}\NormalTok{, }\AttributeTok{color =} \StringTok{"black"}\NormalTok{, }\AttributeTok{linetype =} \StringTok{"dashed"}\NormalTok{, }\AttributeTok{alpha =} \FloatTok{0.2}\NormalTok{) }\SpecialCharTok{+}
  \FunctionTok{geom\_hline}\NormalTok{(}\AttributeTok{yintercept =} \DecValTok{1}\NormalTok{, }\AttributeTok{color =} \StringTok{"grey"}\NormalTok{) }\SpecialCharTok{+} \FunctionTok{geom\_vline}\NormalTok{(}\AttributeTok{xintercept =} \DecValTok{1}\NormalTok{, }\AttributeTok{color =} \StringTok{"grey"}\NormalTok{) }\SpecialCharTok{+}
  \FunctionTok{xlim}\NormalTok{(}\DecValTok{0}\NormalTok{, }\FloatTok{2.1}\NormalTok{) }\SpecialCharTok{+} \FunctionTok{ylim}\NormalTok{(}\DecValTok{0}\NormalTok{, }\FloatTok{2.1}\NormalTok{) }\SpecialCharTok{+}
  \FunctionTok{labs}\NormalTok{(}\AttributeTok{x =} \StringTok{"Relative Mean Expression in Galactose"}\NormalTok{, }\AttributeTok{y =} \StringTok{"Relative Mean Expression in Glycerol"}\NormalTok{) }\SpecialCharTok{+} 
  \FunctionTok{scale\_color\_manual}\NormalTok{(}\AttributeTok{values =} \FunctionTok{c}\NormalTok{(}\StringTok{"Significant"} \OtherTok{=} \StringTok{"red"}\NormalTok{, }\StringTok{"Not significant"} \OtherTok{=} \StringTok{"black"}\NormalTok{)) }\SpecialCharTok{+}
  \FunctionTok{theme\_bw}\NormalTok{() }\SpecialCharTok{+}
  \FunctionTok{theme}\NormalTok{(}\AttributeTok{legend.position =} \StringTok{"bottom"}\NormalTok{,}
        \AttributeTok{axis.title =} \FunctionTok{element\_text}\NormalTok{(}\AttributeTok{size =} \DecValTok{16}\NormalTok{),}
        \AttributeTok{axis.text.x =} \FunctionTok{element\_text}\NormalTok{(}\AttributeTok{hjust =} \DecValTok{1}\NormalTok{),}
        \AttributeTok{panel.grid.minor =} \FunctionTok{element\_blank}\NormalTok{(),}
        \AttributeTok{panel.grid.major =} \FunctionTok{element\_blank}\NormalTok{())}


\DocumentationTok{\#\#\#}
\NormalTok{plot\_GLYvETH\_sig }\OtherTok{\textless{}{-}} \FunctionTok{ggplot}\NormalTok{(data\_GLY\_ETH, }\FunctionTok{aes}\NormalTok{(}\AttributeTok{x =}\NormalTok{ YFP.MEDIAN.RELATIVE.MEAN\_GLYCEROL, }\AttributeTok{y =}\NormalTok{ YFP.MEDIAN.RELATIVE.MEAN\_ETHANOL, }\AttributeTok{color =}\NormalTok{ significance)) }\SpecialCharTok{+}
  \FunctionTok{geom\_point}\NormalTok{(}\AttributeTok{size =} \DecValTok{2}\NormalTok{, }\AttributeTok{alpha =} \FloatTok{0.5}\NormalTok{) }\SpecialCharTok{+} \CommentTok{\#add standard error bars}
  \FunctionTok{geom\_errorbar}\NormalTok{(}\FunctionTok{aes}\NormalTok{(}\AttributeTok{ymin =}\NormalTok{ YFP.MEDIAN.RELATIVE.MEAN\_ETHANOL }\SpecialCharTok{{-}}\NormalTok{ YFP.MEDIAN.RELATIVE.SD\_ETHANOL, }
                    \AttributeTok{ymax =}\NormalTok{ YFP.MEDIAN.RELATIVE.MEAN\_ETHANOL }\SpecialCharTok{+}\NormalTok{ YFP.MEDIAN.RELATIVE.SD\_ETHANOL), }
                \AttributeTok{width =} \FloatTok{0.0}\NormalTok{) }\SpecialCharTok{+} \CommentTok{\#add horizontal error bars}
  \FunctionTok{geom\_errorbarh}\NormalTok{(}\FunctionTok{aes}\NormalTok{(}\AttributeTok{xmin =}\NormalTok{ YFP.MEDIAN.RELATIVE.MEAN\_GLYCEROL }\SpecialCharTok{{-}}\NormalTok{ YFP.MEDIAN.RELATIVE.SD\_GLYCEROL, }
                     \AttributeTok{xmax =}\NormalTok{ YFP.MEDIAN.RELATIVE.MEAN\_GLYCEROL }\SpecialCharTok{+}\NormalTok{ YFP.MEDIAN.RELATIVE.SD\_GLYCEROL), }
                 \AttributeTok{height =} \FloatTok{0.0}\NormalTok{) }\SpecialCharTok{+}
  \FunctionTok{geom\_abline}\NormalTok{(}\AttributeTok{intercept =} \DecValTok{0}\NormalTok{, }\AttributeTok{slope =} \DecValTok{1}\NormalTok{, }\AttributeTok{color =} \StringTok{"black"}\NormalTok{, }\AttributeTok{linetype =} \StringTok{"dashed"}\NormalTok{, }\AttributeTok{alpha =} \FloatTok{0.2}\NormalTok{) }\SpecialCharTok{+}
  \FunctionTok{geom\_hline}\NormalTok{(}\AttributeTok{yintercept =} \DecValTok{1}\NormalTok{, }\AttributeTok{color =} \StringTok{"grey"}\NormalTok{) }\SpecialCharTok{+} \FunctionTok{geom\_vline}\NormalTok{(}\AttributeTok{xintercept =} \DecValTok{1}\NormalTok{, }\AttributeTok{color =} \StringTok{"grey"}\NormalTok{) }\SpecialCharTok{+}
  \FunctionTok{xlim}\NormalTok{(}\DecValTok{0}\NormalTok{, }\FloatTok{2.1}\NormalTok{) }\SpecialCharTok{+} \FunctionTok{ylim}\NormalTok{(}\DecValTok{0}\NormalTok{, }\FloatTok{2.1}\NormalTok{) }\SpecialCharTok{+}
  \FunctionTok{labs}\NormalTok{(}\AttributeTok{x =} \StringTok{"Relative Mean Expression in Glycerol"}\NormalTok{, }\AttributeTok{y =} \StringTok{"Relative Mean Expression in Ethanol"}\NormalTok{) }\SpecialCharTok{+} 
  \FunctionTok{scale\_color\_manual}\NormalTok{(}\AttributeTok{values =} \FunctionTok{c}\NormalTok{(}\StringTok{"Significant"} \OtherTok{=} \StringTok{"red"}\NormalTok{, }\StringTok{"Not significant"} \OtherTok{=} \StringTok{"black"}\NormalTok{)) }\SpecialCharTok{+}
  \FunctionTok{theme\_bw}\NormalTok{() }\SpecialCharTok{+}
  \FunctionTok{theme}\NormalTok{(}\AttributeTok{legend.position =} \StringTok{"bottom"}\NormalTok{,}
        \AttributeTok{axis.title =} \FunctionTok{element\_text}\NormalTok{(}\AttributeTok{size =} \DecValTok{16}\NormalTok{),}
        \AttributeTok{axis.text.x =} \FunctionTok{element\_text}\NormalTok{(}\AttributeTok{hjust =} \DecValTok{1}\NormalTok{),}
        \AttributeTok{panel.grid.minor =} \FunctionTok{element\_blank}\NormalTok{(),}
        \AttributeTok{panel.grid.major =} \FunctionTok{element\_blank}\NormalTok{())}

\DocumentationTok{\#\#\#}
\NormalTok{plot\_expMutants }\OtherTok{\textless{}{-}} \FunctionTok{ggplot}\NormalTok{(}\FunctionTok{subset}\NormalTok{(ExpressionFitnessData\_allEnvironments\_numProm, }\SpecialCharTok{!}\NormalTok{(CLASS }\SpecialCharTok{\%in\%}\NormalTok{ unmutated)), }
       \FunctionTok{aes}\NormalTok{(}\AttributeTok{x =}\NormalTok{ NumPromMutation, }\AttributeTok{y =}\NormalTok{ YFP.MEDIAN.RELATIVE.MEAN, }\AttributeTok{color =}\NormalTok{ ENVIRONMENT)) }\SpecialCharTok{+} 
  \FunctionTok{geom\_hline}\NormalTok{(}\AttributeTok{yintercept =} \DecValTok{1}\NormalTok{, }\AttributeTok{linetype=}\StringTok{"dashed"}\NormalTok{, }\AttributeTok{color =} \StringTok{"black"}\NormalTok{) }\SpecialCharTok{+}
  \FunctionTok{geom\_point}\NormalTok{(}\AttributeTok{size =} \FloatTok{1.5}\NormalTok{, }\AttributeTok{alpha =} \FloatTok{0.5}\NormalTok{, }\AttributeTok{position =} \FunctionTok{position\_dodge}\NormalTok{(}\AttributeTok{width =} \FloatTok{0.5}\NormalTok{))  }\SpecialCharTok{+} 
  \FunctionTok{geom\_errorbar}\NormalTok{(}\FunctionTok{aes}\NormalTok{(}\AttributeTok{ymin =}\NormalTok{ YFP.MEDIAN.RELATIVE.MEAN }\SpecialCharTok{{-}} \FloatTok{1.96}\SpecialCharTok{*}\NormalTok{YFP.MEDIAN.RELATIVE.SD}\SpecialCharTok{/}\FunctionTok{sqrt}\NormalTok{(}\DecValTok{4}\NormalTok{), }
                    \AttributeTok{ymax =}\NormalTok{ YFP.MEDIAN.RELATIVE.MEAN }\SpecialCharTok{+} \FloatTok{1.96}\SpecialCharTok{*}\NormalTok{YFP.MEDIAN.RELATIVE.SD}\SpecialCharTok{/}\FunctionTok{sqrt}\NormalTok{(}\DecValTok{4}\NormalTok{)), }
                \AttributeTok{width =} \FloatTok{0.1}\NormalTok{,}\AttributeTok{alpha =} \FloatTok{0.5}\NormalTok{, }\AttributeTok{position =} \FunctionTok{position\_dodge}\NormalTok{(}\AttributeTok{width =} \FloatTok{0.5}\NormalTok{)) }\SpecialCharTok{+}
  \FunctionTok{labs}\NormalTok{(}\AttributeTok{x =} \StringTok{"Mutant Genotypes"}\NormalTok{, }\AttributeTok{y =} \StringTok{"Relative YFP Expression"}\NormalTok{) }\SpecialCharTok{+}  
  \CommentTok{\#facet\_grid(\textasciitilde{}NumPromMutation)+}
  \FunctionTok{theme\_bw}\NormalTok{() }\SpecialCharTok{+}   \FunctionTok{scale\_color\_manual}\NormalTok{(}\AttributeTok{values =}\NormalTok{ named\_colors) }\SpecialCharTok{+} \FunctionTok{scale\_fill\_manual}\NormalTok{(}\AttributeTok{values =}\NormalTok{ named\_colors) }\SpecialCharTok{+}
  \FunctionTok{theme}\NormalTok{(}\AttributeTok{panel.border =} \FunctionTok{element\_rect}\NormalTok{(}\AttributeTok{linetype =} \DecValTok{1}\NormalTok{, }\AttributeTok{fill =} \StringTok{"NA"}\NormalTok{),}
        \AttributeTok{plot.background =} \FunctionTok{element\_rect}\NormalTok{(}\AttributeTok{color =} \ConstantTok{NA}\NormalTok{),}\AttributeTok{legend.position=}\StringTok{"top"}\NormalTok{,}
        \AttributeTok{axis.text=}\FunctionTok{element\_text}\NormalTok{(}\AttributeTok{size=} \DecValTok{10}\NormalTok{), }\AttributeTok{axis.title=}\FunctionTok{element\_text}\NormalTok{(}\AttributeTok{size =}\DecValTok{14}\NormalTok{),}
        \CommentTok{\#        axis.text.x=element\_text(angle=90,hjust=1),}
        \AttributeTok{panel.grid.minor =} \FunctionTok{element\_blank}\NormalTok{(),}
        \AttributeTok{panel.grid.major =} \FunctionTok{element\_blank}\NormalTok{())}

\DocumentationTok{\#\#\#Save the plots}
\FunctionTok{ggsave}\NormalTok{(}\StringTok{"./ManuscriptFigures/Fig3\_ExpressionEffects.pdf"}\NormalTok{, }\AttributeTok{plot =}\NormalTok{ plot\_expMutants, }\AttributeTok{width =} \DecValTok{8}\NormalTok{, }\AttributeTok{height =} \DecValTok{6}\NormalTok{, }\AttributeTok{units =} \StringTok{"in"}\NormalTok{, }\AttributeTok{dpi =} \DecValTok{300}\NormalTok{)}
\end{Highlighting}
\end{Shaded}

\begin{verbatim}
## Warning: No shared levels found between `names(values)` of the manual scale and the
## data's fill values.
\end{verbatim}

\begin{Shaded}
\begin{Highlighting}[]
\FunctionTok{ggsave}\NormalTok{(}\StringTok{"./ManuscriptFigures/Fig3\_GLUvGAL.pdf"}\NormalTok{, }\AttributeTok{plot =}\NormalTok{ plot\_GLUvGAL\_sig, }\AttributeTok{width =} \DecValTok{6}\NormalTok{, }\AttributeTok{height =} \DecValTok{6}\NormalTok{, }\AttributeTok{units =} \StringTok{"in"}\NormalTok{, }\AttributeTok{dpi =} \DecValTok{300}\NormalTok{)}
\end{Highlighting}
\end{Shaded}

\begin{verbatim}
## Warning: Removed 2 rows containing missing values or values outside the scale range
## (`geom_point()`).
\end{verbatim}

\begin{verbatim}
## Warning: Removed 3 rows containing missing values or values outside the scale range
## (`geom_errorbarh()`).
\end{verbatim}

\begin{Shaded}
\begin{Highlighting}[]
\FunctionTok{ggsave}\NormalTok{(}\StringTok{"./ManuscriptFigures/Fig3\_GLUvGLY.pdf"}\NormalTok{, }\AttributeTok{plot =}\NormalTok{ plot\_GLUvGLY\_sig, }\AttributeTok{width =} \DecValTok{6}\NormalTok{, }\AttributeTok{height =} \DecValTok{6}\NormalTok{, }\AttributeTok{units =} \StringTok{"in"}\NormalTok{, }\AttributeTok{dpi =} \DecValTok{300}\NormalTok{)}
\end{Highlighting}
\end{Shaded}

\begin{verbatim}
## Warning: Removed 3 rows containing missing values or values outside the scale range
## (`geom_point()`).
\end{verbatim}

\begin{verbatim}
## Warning: Removed 4 rows containing missing values or values outside the scale range
## (`geom_errorbarh()`).
\end{verbatim}

\begin{Shaded}
\begin{Highlighting}[]
\FunctionTok{ggsave}\NormalTok{(}\StringTok{"./ManuscriptFigures/Fig3\_GLUvETH.pdf"}\NormalTok{, }\AttributeTok{plot =}\NormalTok{ plot\_GLUvETH\_sig, }\AttributeTok{width =} \DecValTok{6}\NormalTok{, }\AttributeTok{height =} \DecValTok{6}\NormalTok{, }\AttributeTok{units =} \StringTok{"in"}\NormalTok{, }\AttributeTok{dpi =} \DecValTok{300}\NormalTok{)}
\end{Highlighting}
\end{Shaded}

\begin{verbatim}
## Warning: Removed 2 rows containing missing values or values outside the scale range
## (`geom_point()`).
\end{verbatim}

\begin{verbatim}
## Warning: Removed 3 rows containing missing values or values outside the scale range
## (`geom_errorbarh()`).
\end{verbatim}

\begin{Shaded}
\begin{Highlighting}[]
\FunctionTok{ggsave}\NormalTok{(}\StringTok{"./ManuscriptFigures/Fig3\_GALvGLY.pdf"}\NormalTok{, }\AttributeTok{plot =}\NormalTok{ plot\_GALvGLY\_sig, }\AttributeTok{width =} \DecValTok{6}\NormalTok{, }\AttributeTok{height =} \DecValTok{6}\NormalTok{, }\AttributeTok{units =} \StringTok{"in"}\NormalTok{, }\AttributeTok{dpi =} \DecValTok{300}\NormalTok{)}
\end{Highlighting}
\end{Shaded}

\begin{verbatim}
## Warning: Removed 2 rows containing missing values or values outside the scale range
## (`geom_point()`).
## Removed 3 rows containing missing values or values outside the scale range
## (`geom_errorbarh()`).
\end{verbatim}

\begin{Shaded}
\begin{Highlighting}[]
\FunctionTok{ggsave}\NormalTok{(}\StringTok{"./ManuscriptFigures/Fig3\_GALvETH.pdf"}\NormalTok{, }\AttributeTok{plot =}\NormalTok{ plot\_GALvETH\_sig, }\AttributeTok{width =} \DecValTok{6}\NormalTok{, }\AttributeTok{height =} \DecValTok{6}\NormalTok{, }\AttributeTok{units =} \StringTok{"in"}\NormalTok{, }\AttributeTok{dpi =} \DecValTok{300}\NormalTok{)}
\end{Highlighting}
\end{Shaded}

\begin{verbatim}
## Warning: Removed 1 row containing missing values or values outside the scale range
## (`geom_point()`).
\end{verbatim}

\begin{verbatim}
## Warning: Removed 2 rows containing missing values or values outside the scale range
## (`geom_errorbarh()`).
\end{verbatim}

\begin{Shaded}
\begin{Highlighting}[]
\FunctionTok{ggsave}\NormalTok{(}\StringTok{"./ManuscriptFigures/Fig3\_GLYvETH.pdf"}\NormalTok{, }\AttributeTok{plot =}\NormalTok{ plot\_GLYvETH\_sig, }\AttributeTok{width =} \DecValTok{6}\NormalTok{, }\AttributeTok{height =} \DecValTok{6}\NormalTok{, }\AttributeTok{units =} \StringTok{"in"}\NormalTok{, }\AttributeTok{dpi =} \DecValTok{300}\NormalTok{)}
\end{Highlighting}
\end{Shaded}

\begin{verbatim}
## Warning: Removed 2 rows containing missing values or values outside the scale range
## (`geom_point()`).
\end{verbatim}

\begin{verbatim}
## Warning: Removed 3 rows containing missing values or values outside the scale range
## (`geom_errorbarh()`).
\end{verbatim}

\hypertarget{figure-3c-d-variation-in-gene-expression-among-environments}{%
\subsubsection{Figure 3C-D: Variation in gene expression among
environments}\label{figure-3c-d-variation-in-gene-expression-among-environments}}

The following code produces the panels in Figure 3C and Figure 3D. These
are the fitness curves in the different environment. The x-axis values
(expression levels) have all been normalized by the value of the
unmutated, reference strain in the glucose environment.

\begin{Shaded}
\begin{Highlighting}[]
\NormalTok{columns\_to\_keep }\OtherTok{\textless{}{-}} \FunctionTok{c}\NormalTok{(}\StringTok{"YFP.MEDIAN.ADJUST.MEAN\_driftcorrected"}\NormalTok{, }\StringTok{"YFP.MEDIAN.ADJUST.SD\_driftcorrected"}\NormalTok{, }
                     \StringTok{"YFP.MEDIAN.RELATIVE.MEAN"}\NormalTok{, }\StringTok{"MUTATION.Expression"}\NormalTok{, }\StringTok{"MUTATION.Fitness"}\NormalTok{, }\StringTok{"ENVIRONMENT"}\NormalTok{, }\StringTok{"Fitness"}\NormalTok{, }\StringTok{"ID"}\NormalTok{, }\StringTok{"STRAIN.Expression"}\NormalTok{, }\StringTok{"STRAIN.Fitness"}\NormalTok{, }\StringTok{"CLASS"}\NormalTok{)}

\NormalTok{ExpressionFitnessData\_allEnvironments}\SpecialCharTok{$}\NormalTok{ENVIRONMENT }\OtherTok{\textless{}{-}} \FunctionTok{fct\_relevel}\NormalTok{(ExpressionFitnessData\_allEnvironments}\SpecialCharTok{$}\NormalTok{ENVIRONMENT, }\FunctionTok{c}\NormalTok{(}\StringTok{"GLUCOSE"}\NormalTok{, }\StringTok{"GALACTOSE"}\NormalTok{, }\StringTok{"GLYCEROL"}\NormalTok{, }\StringTok{"ETHANOL"}\NormalTok{))}

\CommentTok{\#There are many columns that have only numbers but are not treated as numeric. Change these columns to be numeric. Specifically, make the columns that only have numbers to be numeric.}


\NormalTok{ExpressionFitnessData\_allEnvironments }\OtherTok{\textless{}{-}}\NormalTok{ ExpressionFitnessData\_allEnvironments }\SpecialCharTok{\%\textgreater{}\%} \FunctionTok{mutate\_at}\NormalTok{(}\FunctionTok{vars}\NormalTok{(YFP.MEDIAN.ADJUST.MEAN\_driftcorrected, YFP.MEDIAN.ADJUST.SD\_driftcorrected, YFP.MEDIAN.RELATIVE.MEAN, Fitness, Low}\FloatTok{.95}\NormalTok{, High}\FloatTok{.95}\NormalTok{,}
\NormalTok{                                                                                                  YFP.MEDIAN.RELATIVE.MEAN, YFP.MEDIAN.RELATIVE.SD), as.numeric)}

\CommentTok{\#Weight the columns for LOWESS }
\NormalTok{ExpressionFitnessData\_allEnvironments\_FC }\OtherTok{\textless{}{-}}\NormalTok{ ExpressionFitnessData\_allEnvironments }\SpecialCharTok{\%\textgreater{}\%} \FunctionTok{select}\NormalTok{(columns\_to\_keep)}\SpecialCharTok{\%\textgreater{}\%} 
  \FunctionTok{mutate}\NormalTok{(}\AttributeTok{weights =} \FunctionTok{ifelse}\NormalTok{(STRAIN.Fitness }\SpecialCharTok{==} \StringTok{"Y1189"}\NormalTok{, }\DecValTok{100}\NormalTok{, }
                          \FunctionTok{ifelse}\NormalTok{(STRAIN.Fitness }\SpecialCharTok{==} \StringTok{"Y1177"}\NormalTok{, }\DecValTok{100}\NormalTok{, }\DecValTok{1}\NormalTok{))) }\SpecialCharTok{\%\textgreater{}\%}
  \FunctionTok{mutate}\NormalTok{(}\AttributeTok{YFPmed\_adjusted\_RelToRefGLU =}\NormalTok{ YFP.MEDIAN.ADJUST.MEAN\_driftcorrected}\SpecialCharTok{/}\FunctionTok{subset}\NormalTok{(ExpressionFitnessData\_allEnvironments, ENVIRONMENT }\SpecialCharTok{==} \StringTok{"GLUCOSE"} \SpecialCharTok{\&}\NormalTok{ STRAIN.Fitness }\SpecialCharTok{==} \StringTok{"Y1189"}\NormalTok{)}\SpecialCharTok{$}\NormalTok{YFP.MEDIAN.ADJUST.MEAN\_driftcorrected)}
\end{Highlighting}
\end{Shaded}

\begin{verbatim}
## Warning: Using an external vector in selections was deprecated in tidyselect 1.1.0.
## i Please use `all_of()` or `any_of()` instead.
##   # Was:
##   data %>% select(columns_to_keep)
## 
##   # Now:
##   data %>% select(all_of(columns_to_keep))
## 
## See <https://tidyselect.r-lib.org/reference/faq-external-vector.html>.
## This warning is displayed once every 8 hours.
## Call `lifecycle::last_lifecycle_warnings()` to see where this warning was
## generated.
\end{verbatim}

\begin{Shaded}
\begin{Highlighting}[]
\CommentTok{\# Environment subsets}
\NormalTok{ExpressionFitnessData\_GLU }\OtherTok{\textless{}{-}}\NormalTok{ ExpressionFitnessData\_allEnvironments\_FC[ExpressionFitnessData\_allEnvironments\_FC}\SpecialCharTok{$}\NormalTok{ENVIRONMENT }\SpecialCharTok{==} \StringTok{"GLUCOSE"}\NormalTok{,]}
\NormalTok{ExpressionFitnessData\_GAL }\OtherTok{\textless{}{-}}\NormalTok{ ExpressionFitnessData\_allEnvironments\_FC[ExpressionFitnessData\_allEnvironments\_FC}\SpecialCharTok{$}\NormalTok{ENVIRONMENT }\SpecialCharTok{==} \StringTok{"GALACTOSE"}\NormalTok{,]}
\NormalTok{ExpressionFitnessData\_GLY }\OtherTok{\textless{}{-}}\NormalTok{ ExpressionFitnessData\_allEnvironments\_FC[ExpressionFitnessData\_allEnvironments\_FC}\SpecialCharTok{$}\NormalTok{ENVIRONMENT }\SpecialCharTok{==} \StringTok{"GLYCEROL"}\NormalTok{,]}
\NormalTok{ExpressionFitnessData\_ETH }\OtherTok{\textless{}{-}}\NormalTok{ ExpressionFitnessData\_allEnvironments\_FC[ExpressionFitnessData\_allEnvironments\_FC}\SpecialCharTok{$}\NormalTok{ENVIRONMENT }\SpecialCharTok{==} \StringTok{"ETHANOL"}\NormalTok{,]}

\DocumentationTok{\#\#\# Modeling Fitness Curves {-}{-}{-}{-}{-}{-}{-}{-}}
\CommentTok{\# Make a loess model with the weights relating YFP.MEDIAN.RELATIVE.MEAN and Fitness; The negative control}
\CommentTok{\# and reference strain are weighted 100, while all other strains are weighted 1. These strains also have more}
\CommentTok{\# measurements. }
\NormalTok{model\_GLU }\OtherTok{\textless{}{-}} \FunctionTok{loess}\NormalTok{(Fitness }\SpecialCharTok{\textasciitilde{}}\NormalTok{ YFP.MEDIAN.RELATIVE.MEAN , }\AttributeTok{data =}\NormalTok{ ExpressionFitnessData\_GLU, }
                   \AttributeTok{weights =}\NormalTok{ weights)}
\NormalTok{model\_GAL }\OtherTok{\textless{}{-}} \FunctionTok{loess}\NormalTok{(Fitness }\SpecialCharTok{\textasciitilde{}}\NormalTok{ YFP.MEDIAN.RELATIVE.MEAN , }\AttributeTok{data =}\NormalTok{ ExpressionFitnessData\_GAL, }
                   \AttributeTok{weights =}\NormalTok{ weights)}
\NormalTok{model\_GLY }\OtherTok{\textless{}{-}} \FunctionTok{loess}\NormalTok{(Fitness }\SpecialCharTok{\textasciitilde{}}\NormalTok{ YFP.MEDIAN.RELATIVE.MEAN , }\AttributeTok{data =}\NormalTok{ ExpressionFitnessData\_GLY, }
                   \AttributeTok{weights =}\NormalTok{ weights)}
\NormalTok{model\_ETH }\OtherTok{\textless{}{-}} \FunctionTok{loess}\NormalTok{(Fitness }\SpecialCharTok{\textasciitilde{}}\NormalTok{ YFP.MEDIAN.RELATIVE.MEAN , }\AttributeTok{data =}\NormalTok{ ExpressionFitnessData\_ETH, }
                   \AttributeTok{weights =}\NormalTok{ weights)}

\DocumentationTok{\#\# Visualize the curves}
\NormalTok{named\_colors }\OtherTok{\textless{}{-}} \FunctionTok{c}\NormalTok{(}\StringTok{"GLUCOSE"} \OtherTok{=} \StringTok{"\#0072B2"}\NormalTok{, }\StringTok{"GALACTOSE"} \OtherTok{=} \StringTok{"\#009E73"}\NormalTok{, }\StringTok{"GLYCEROL"} \OtherTok{=} \StringTok{"\#CC79A7"}\NormalTok{, }\StringTok{"ETHANOL"} \OtherTok{=} \StringTok{"\#D55E00"}\NormalTok{)}


\NormalTok{plot\_fitnessCurves\_relGLU }\OtherTok{\textless{}{-}} \FunctionTok{ggplot}\NormalTok{(ExpressionFitnessData\_allEnvironments\_FC, }\FunctionTok{aes}\NormalTok{(}\AttributeTok{x =}\NormalTok{ YFPmed\_adjusted\_RelToRefGLU,}
                                                                        \AttributeTok{y =}\NormalTok{ Fitness, }\AttributeTok{col =}\NormalTok{ ENVIRONMENT)) }\SpecialCharTok{+} 
  \FunctionTok{geom\_smooth}\NormalTok{(}\AttributeTok{method =} \StringTok{"loess"}\NormalTok{, }\AttributeTok{formula =}\NormalTok{ y }\SpecialCharTok{\textasciitilde{}}\NormalTok{ x, }\AttributeTok{se =} \ConstantTok{TRUE}\NormalTok{, }\AttributeTok{color =} \StringTok{"\#0072B2"}\NormalTok{, }\AttributeTok{fill =} \StringTok{"\#0072B2"}\NormalTok{, }\AttributeTok{size =} \DecValTok{1}\NormalTok{, }\AttributeTok{alpha =} \FloatTok{0.2}\NormalTok{) }\SpecialCharTok{+} 
  \FunctionTok{geom\_smooth}\NormalTok{(}\AttributeTok{method =} \StringTok{"loess"}\NormalTok{, }\AttributeTok{formula =}\NormalTok{ y }\SpecialCharTok{\textasciitilde{}}\NormalTok{ x, }\AttributeTok{se =} \ConstantTok{TRUE}\NormalTok{, }\AttributeTok{color =} \StringTok{"\#009E73"}\NormalTok{, }\AttributeTok{fill =} \StringTok{"\#009E73"}\NormalTok{, }\AttributeTok{size =} \DecValTok{1}\NormalTok{, }\AttributeTok{data =}\NormalTok{ ExpressionFitnessData\_GAL, }\AttributeTok{alpha =} \FloatTok{0.2}\NormalTok{) }\SpecialCharTok{+} 
  \FunctionTok{geom\_smooth}\NormalTok{(}\AttributeTok{method =} \StringTok{"loess"}\NormalTok{, }\AttributeTok{formula =}\NormalTok{ y }\SpecialCharTok{\textasciitilde{}}\NormalTok{ x, }\AttributeTok{se =} \ConstantTok{TRUE}\NormalTok{, }\AttributeTok{color =} \StringTok{"\#CC79A7"}\NormalTok{, }\AttributeTok{fill =} \StringTok{"\#CC79A7"}\NormalTok{, }\AttributeTok{size =} \DecValTok{1}\NormalTok{, }\AttributeTok{data =}\NormalTok{ ExpressionFitnessData\_GLY, }\AttributeTok{alpha =} \FloatTok{0.2}\NormalTok{) }\SpecialCharTok{+} 
  \FunctionTok{geom\_smooth}\NormalTok{(}\AttributeTok{method =} \StringTok{"loess"}\NormalTok{, }\AttributeTok{formula =}\NormalTok{ y }\SpecialCharTok{\textasciitilde{}}\NormalTok{ x, }\AttributeTok{se =} \ConstantTok{TRUE}\NormalTok{, }\AttributeTok{color =} \StringTok{"\#D55E00"}\NormalTok{, }\AttributeTok{fill =} \StringTok{"\#D55E00"}\NormalTok{,}\AttributeTok{size =} \DecValTok{1}\NormalTok{, }\AttributeTok{data =}\NormalTok{ ExpressionFitnessData\_ETH, }\AttributeTok{alpha =} \FloatTok{0.2}\NormalTok{) }\SpecialCharTok{+} 
  \FunctionTok{geom\_hline}\NormalTok{(}\AttributeTok{yintercept =} \DecValTok{1}\NormalTok{, }\AttributeTok{linetype =} \StringTok{"dashed"}\NormalTok{) }\SpecialCharTok{+} \FunctionTok{geom\_vline}\NormalTok{(}\AttributeTok{xintercept =} \DecValTok{1}\NormalTok{, }\AttributeTok{linetype =} \StringTok{"dashed"}\NormalTok{) }\SpecialCharTok{+}
  \FunctionTok{geom\_point}\NormalTok{(}\AttributeTok{data =}\NormalTok{ ExpressionFitnessData\_allEnvironments\_FC[ExpressionFitnessData\_allEnvironments\_FC}\SpecialCharTok{$}\NormalTok{STRAIN.Fitness }\SpecialCharTok{==} \StringTok{"Y1189"}\NormalTok{,],}
             \FunctionTok{aes}\NormalTok{(}\AttributeTok{x =}\NormalTok{ YFPmed\_adjusted\_RelToRefGLU, }\AttributeTok{y =}\NormalTok{ Fitness), }
              \AttributeTok{col =} \StringTok{"black"}\NormalTok{, }\AttributeTok{size =} \FloatTok{2.5}\NormalTok{) }\SpecialCharTok{+}
  \FunctionTok{scale\_y\_continuous}\NormalTok{(}\AttributeTok{breaks =} \FunctionTok{c}\NormalTok{(}\FloatTok{0.94}\NormalTok{, }\FloatTok{0.97}\NormalTok{, }\FloatTok{1.00}\NormalTok{, }\FloatTok{1.03}\NormalTok{)) }\SpecialCharTok{+}
  \FunctionTok{theme\_bw}\NormalTok{() }\SpecialCharTok{+}   \FunctionTok{scale\_color\_manual}\NormalTok{(}\AttributeTok{values =}\NormalTok{ named\_colors) }\SpecialCharTok{+} \FunctionTok{scale\_fill\_manual}\NormalTok{(}\AttributeTok{values =}\NormalTok{ named\_colors) }\SpecialCharTok{+}
  \FunctionTok{theme}\NormalTok{(}\AttributeTok{legend.position =} \StringTok{"bottom"}\NormalTok{,}
        \AttributeTok{axis.text.x =} \FunctionTok{element\_text}\NormalTok{(}\AttributeTok{hjust =} \DecValTok{1}\NormalTok{),}
        \AttributeTok{panel.grid.minor =} \FunctionTok{element\_blank}\NormalTok{(),}
        \AttributeTok{panel.grid.major =} \FunctionTok{element\_blank}\NormalTok{()) }\SpecialCharTok{+} 
  \FunctionTok{labs}\NormalTok{(}\AttributeTok{x =} \StringTok{"Expression Level (Normalized to Ref. Strain in Glucose)"}\NormalTok{, }\AttributeTok{y =} \StringTok{"Relative Fitness"}\NormalTok{)}
\end{Highlighting}
\end{Shaded}

\begin{verbatim}
## Warning: Using `size` aesthetic for lines was deprecated in ggplot2 3.4.0.
## i Please use `linewidth` instead.
## This warning is displayed once every 8 hours.
## Call `lifecycle::last_lifecycle_warnings()` to see where this warning was
## generated.
\end{verbatim}

\begin{Shaded}
\begin{Highlighting}[]
\NormalTok{plot\_fitnessCurves\_relGLU\_facet\_oneCol }\OtherTok{\textless{}{-}}\NormalTok{ plot\_fitnessCurves\_relGLU }\SpecialCharTok{+} 
  \FunctionTok{geom\_point}\NormalTok{(}\AttributeTok{alpha =} \FloatTok{0.5}\NormalTok{) }\SpecialCharTok{+}   
  \FunctionTok{facet\_wrap}\NormalTok{(}\SpecialCharTok{\textasciitilde{}}\FunctionTok{factor}\NormalTok{(ENVIRONMENT, }\AttributeTok{levels=}\FunctionTok{c}\NormalTok{(}\StringTok{\textquotesingle{}GLUCOSE\textquotesingle{}}\NormalTok{, }\StringTok{\textquotesingle{}GALACTOSE\textquotesingle{}}\NormalTok{, }\StringTok{\textquotesingle{}GLYCEROL\textquotesingle{}}\NormalTok{, }\StringTok{\textquotesingle{}ETHANOL\textquotesingle{}}\NormalTok{)), }\AttributeTok{ncol =} \DecValTok{1}\NormalTok{) }


\FunctionTok{ggsave}\NormalTok{(}\StringTok{"./ManuscriptFigures/Fig3\_fitnessCurves\_overlain.pdf"}\NormalTok{, }\AttributeTok{plot =}\NormalTok{ plot\_fitnessCurves\_relGLU, }\AttributeTok{width =} \FloatTok{6.0}\NormalTok{, }\AttributeTok{height =} \FloatTok{4.0}\NormalTok{, }\AttributeTok{units =} \StringTok{"in"}\NormalTok{, }\AttributeTok{dpi =} \DecValTok{300}\NormalTok{)}
\end{Highlighting}
\end{Shaded}

\begin{verbatim}
## Warning: Removed 12 rows containing non-finite outside the scale range
## (`stat_smooth()`).
\end{verbatim}

\begin{verbatim}
## Warning: Removed 3 rows containing non-finite outside the scale range (`stat_smooth()`).
## Removed 3 rows containing non-finite outside the scale range (`stat_smooth()`).
## Removed 3 rows containing non-finite outside the scale range (`stat_smooth()`).
\end{verbatim}

\begin{verbatim}
## Warning: No shared levels found between `names(values)` of the manual scale and the
## data's colour values.
\end{verbatim}

\begin{verbatim}
## Warning: No shared levels found between `names(values)` of the manual scale and the
## data's fill values.
\end{verbatim}

\begin{Shaded}
\begin{Highlighting}[]
\FunctionTok{ggsave}\NormalTok{(}\StringTok{"./ManuscriptFigures/Fig3\_fitnessCurves\_facet.pdf"}\NormalTok{, }\AttributeTok{plot =}\NormalTok{ plot\_fitnessCurves\_relGLU\_facet\_oneCol, }\AttributeTok{width =} \FloatTok{6.0}\NormalTok{, }\AttributeTok{height =} \DecValTok{9}\NormalTok{, }\AttributeTok{units =} \StringTok{"in"}\NormalTok{, }\AttributeTok{dpi =} \DecValTok{300}\NormalTok{)}
\end{Highlighting}
\end{Shaded}

\begin{verbatim}
## Warning: Removed 12 rows containing non-finite outside the scale range
## (`stat_smooth()`).
\end{verbatim}

\begin{verbatim}
## Warning: Removed 3 rows containing non-finite outside the scale range (`stat_smooth()`).
## Removed 3 rows containing non-finite outside the scale range (`stat_smooth()`).
## Removed 3 rows containing non-finite outside the scale range (`stat_smooth()`).
\end{verbatim}

\begin{verbatim}
## Warning: No shared levels found between `names(values)` of the manual scale and the
## data's fill values.
\end{verbatim}

\begin{verbatim}
## Warning: Removed 12 rows containing missing values or values outside the scale range
## (`geom_point()`).
\end{verbatim}

\hypertarget{figure-4}{%
\subsubsection{Figure 4}\label{figure-4}}

The following code produces the panels in Figure 4. The code also 1)
runs tests to check for normality for comparing expression and fitness
effects between TFBS and TATA box mutations and 2) runs the Mann-Whitney
U test to compare the expression effects fitness effects of TFBS and
TATA box mutations across environments that are reported in the
manuscripts. Note that t-test also appears appropriate for comparing the
expression effects of TFBS and TATA box mutations among environments and
results in the same inference.

\begin{Shaded}
\begin{Highlighting}[]
\CommentTok{\# Strain numbers below correspond to "STRAIN.expression" column. The corresponding STRAIN.fitness with the mutations are also picked up}
\CommentTok{\# using this approach.}
\NormalTok{mut\_cisMotifs }\OtherTok{\textless{}{-}} \FunctionTok{c}\NormalTok{(}\StringTok{"Y1896"}\NormalTok{, }\StringTok{"Y1897"}\NormalTok{, }\StringTok{"Y1898"}\NormalTok{, }\StringTok{"Y1960"}\NormalTok{, }\StringTok{"Y1892"}\NormalTok{, }\StringTok{"Y1901"}\NormalTok{, }\StringTok{"Y1903"}\NormalTok{, }\CommentTok{\#TFBS mutants; }
                   \StringTok{"Y2528"}\NormalTok{, }\StringTok{"Y2566"}\NormalTok{, }\StringTok{"Y2605"}\NormalTok{, }\StringTok{"Y2651"}\NormalTok{, }\StringTok{"Y2546"}\NormalTok{, }\StringTok{"Y2656"}\NormalTok{, }\StringTok{"Y2550"}\NormalTok{) }\CommentTok{\#TATA mutants}


\DocumentationTok{\#\# Tests to detect effects on expression {-}{-}{-}{-}{-}{-}{-}}
\NormalTok{Expression\_cisMotifs }\OtherTok{\textless{}{-}}\NormalTok{ ExpressionFitnessData\_allEnvironments }\SpecialCharTok{\%\textgreater{}\%}
  \FunctionTok{filter}\NormalTok{(STRAIN.Expression }\SpecialCharTok{\%in\%}\NormalTok{ mut\_cisMotifs) }\SpecialCharTok{\%\textgreater{}\%}  \CommentTok{\# Keep the necessary columns relevant for assessing fitness effects among environments}
  \FunctionTok{select}\NormalTok{(STRAIN.Expression, STRAIN.Fitness, ENVIRONMENT, MUTATION.Expression, CLASS,YFP.MEDIAN.RELATIVE.MEAN ,YFP.MEDIAN.RELATIVE.SD) }\SpecialCharTok{\%\textgreater{}\%} \CommentTok{\#Change the columns with only numbers to numeric}
  \FunctionTok{mutate}\NormalTok{(}\AttributeTok{YFP.MEDIAN.RELATIVE.MEAN =} \FunctionTok{as.numeric}\NormalTok{(YFP.MEDIAN.RELATIVE.MEAN),}
         \AttributeTok{YFP.MEDIAN.RELATIVE.SD =} \FunctionTok{as.numeric}\NormalTok{(YFP.MEDIAN.RELATIVE.SD))}

\NormalTok{Expression\_cisMotifs\_summary }\OtherTok{\textless{}{-}}\NormalTok{ Expression\_cisMotifs }\SpecialCharTok{\%\textgreater{}\%} 
  \FunctionTok{group\_by}\NormalTok{(STRAIN.Expression, ENVIRONMENT, MUTATION.Expression, CLASS) }\SpecialCharTok{\%\textgreater{}\%} 
  \FunctionTok{summarise}\NormalTok{(}\AttributeTok{YFP.MEDIAN.RELATIVE.MEAN =} \FunctionTok{mean}\NormalTok{(YFP.MEDIAN.RELATIVE.MEAN),}
            \AttributeTok{YFP.MEDIAN.RELATIVE.SD =} \FunctionTok{mean}\NormalTok{(YFP.MEDIAN.RELATIVE.MEAN)) }
\end{Highlighting}
\end{Shaded}

\begin{verbatim}
## `summarise()` has grouped output by 'STRAIN.Expression', 'ENVIRONMENT',
## 'MUTATION.Expression'. You can override using the `.groups` argument.
\end{verbatim}

\begin{Shaded}
\begin{Highlighting}[]
\NormalTok{cisMotifs\_var }\OtherTok{\textless{}{-}}\NormalTok{ Expression\_cisMotifs\_summary }\SpecialCharTok{\%\textgreater{}\%}
  \FunctionTok{group\_by}\NormalTok{(STRAIN.Expression, CLASS, MUTATION.Expression) }\SpecialCharTok{\%\textgreater{}\%}
  \FunctionTok{summarise}\NormalTok{(}\AttributeTok{variance =} \FunctionTok{var}\NormalTok{(YFP.MEDIAN.RELATIVE.MEAN))}
\end{Highlighting}
\end{Shaded}

\begin{verbatim}
## `summarise()` has grouped output by 'STRAIN.Expression', 'CLASS'. You can
## override using the `.groups` argument.
\end{verbatim}

\begin{Shaded}
\begin{Highlighting}[]
\CommentTok{\#Set the factor order}
\NormalTok{cisMotifs\_var}\SpecialCharTok{$}\NormalTok{CLASS }\OtherTok{\textless{}{-}} \FunctionTok{fct\_relevel}\NormalTok{(cisMotifs\_var}\SpecialCharTok{$}\NormalTok{CLASS, }\FunctionTok{c}\NormalTok{(}\StringTok{"TFBS"}\NormalTok{, }\StringTok{"TATA"}\NormalTok{))}

\DocumentationTok{\#\#Shapiro{-}Wilks normality test shows that variance is normally distributed }
\DocumentationTok{\#\# for the TFBS and TATA classes; t{-}test and F{-}test are appropriate. We will also }
\DocumentationTok{\#\# test with the MW test for robustness. }
\FunctionTok{shapiro.test}\NormalTok{(}\FunctionTok{subset}\NormalTok{(cisMotifs\_var, CLASS }\SpecialCharTok{==} \StringTok{"TFBS"}\NormalTok{)}\SpecialCharTok{$}\NormalTok{variance)}
\end{Highlighting}
\end{Shaded}

\begin{verbatim}
## 
##  Shapiro-Wilk normality test
## 
## data:  subset(cisMotifs_var, CLASS == "TFBS")$variance
## W = 0.91257, p-value = 0.414
\end{verbatim}

\begin{Shaded}
\begin{Highlighting}[]
\FunctionTok{shapiro.test}\NormalTok{(}\FunctionTok{subset}\NormalTok{(cisMotifs\_var, CLASS }\SpecialCharTok{==} \StringTok{"TATA"}\NormalTok{)}\SpecialCharTok{$}\NormalTok{variance)}
\end{Highlighting}
\end{Shaded}

\begin{verbatim}
## 
##  Shapiro-Wilk normality test
## 
## data:  subset(cisMotifs_var, CLASS == "TATA")$variance
## W = 0.93384, p-value = 0.584
\end{verbatim}

\begin{Shaded}
\begin{Highlighting}[]
\FunctionTok{t.test}\NormalTok{(variance }\SpecialCharTok{\textasciitilde{}}\NormalTok{ CLASS, }\AttributeTok{data =}\NormalTok{ cisMotifs\_var, }\AttributeTok{alternative =} \StringTok{"greater"}\NormalTok{)}
\end{Highlighting}
\end{Shaded}

\begin{verbatim}
## 
##  Welch Two Sample t-test
## 
## data:  variance by CLASS
## t = 2.3667, df = 6.5192, p-value = 0.02623
## alternative hypothesis: true difference in means between group TFBS and group TATA is greater than 0
## 95 percent confidence interval:
##  0.0005104549          Inf
## sample estimates:
## mean in group TFBS mean in group TATA 
##       0.0035317674       0.0008523344
\end{verbatim}

\begin{Shaded}
\begin{Highlighting}[]
\FunctionTok{wilcox.test}\NormalTok{(variance }\SpecialCharTok{\textasciitilde{}}\NormalTok{ CLASS, }\AttributeTok{data =}\NormalTok{ cisMotifs\_var,}
            \AttributeTok{alternative =} \StringTok{"greater"}\NormalTok{)  }\DocumentationTok{\#\# Also rejects the null hypothesis}
\end{Highlighting}
\end{Shaded}

\begin{verbatim}
## 
##  Wilcoxon rank sum exact test
## 
## data:  variance by CLASS
## W = 38, p-value = 0.04866
## alternative hypothesis: true location shift is greater than 0
\end{verbatim}

\begin{Shaded}
\begin{Highlighting}[]
\DocumentationTok{\#\# Test to detect effects on fitness {-}{-}{-}{-}{-}{-}{-}}
\DocumentationTok{\#\# Keep the columns relevant for assessing fitness effects among environments}
\NormalTok{FitnessData\_cisMotifs }\OtherTok{\textless{}{-}}\NormalTok{ ExpressionFitnessData\_allEnvironments }\SpecialCharTok{\%\textgreater{}\%} 
  \FunctionTok{filter}\NormalTok{(STRAIN.Expression }\SpecialCharTok{\%in\%}\NormalTok{ mut\_cisMotifs) }\SpecialCharTok{\%\textgreater{}\%}  \CommentTok{\# Keep the necessary columns relevant for assessing fitness effects among environments}
  \FunctionTok{select}\NormalTok{(STRAIN.Expression, STRAIN.Fitness, ENVIRONMENT, MUTATION.Fitness, CLASS, Fitness, Low}\FloatTok{.95}\NormalTok{, High}\FloatTok{.95}\NormalTok{) }

\CommentTok{\#Calculate the mean RelativeFitness for each strain in each environment  }
\NormalTok{summary\_FitnessData\_cisMotifs }\OtherTok{\textless{}{-}}\NormalTok{ FitnessData\_cisMotifs }\SpecialCharTok{\%\textgreater{}\%}
  \FunctionTok{mutate}\NormalTok{(}\AttributeTok{Fitness =} \FunctionTok{as.numeric}\NormalTok{(}\FunctionTok{gsub}\NormalTok{(}\StringTok{"E"}\NormalTok{, }\StringTok{"e"}\NormalTok{, Fitness))) }\SpecialCharTok{\%\textgreater{}\%} 
  \FunctionTok{group\_by}\NormalTok{(STRAIN.Fitness, ENVIRONMENT, CLASS, MUTATION.Fitness) }\SpecialCharTok{\%\textgreater{}\%} 
  \FunctionTok{summarise}\NormalTok{(}\AttributeTok{meanRelativeFitness =} \FunctionTok{mean}\NormalTok{(Fitness),}
            \AttributeTok{sdRelativeFitness =} \FunctionTok{sd}\NormalTok{(Fitness))}
\end{Highlighting}
\end{Shaded}

\begin{verbatim}
## `summarise()` has grouped output by 'STRAIN.Fitness', 'ENVIRONMENT', 'CLASS'.
## You can override using the `.groups` argument.
\end{verbatim}

\begin{Shaded}
\begin{Highlighting}[]
\DocumentationTok{\#\# Test if the meanRelativeFitness varies more for TFBS mutants than TATA mutants}
\DocumentationTok{\#\# in the different environments}
\NormalTok{cisMut\_varFitness }\OtherTok{\textless{}{-}}\NormalTok{ summary\_FitnessData\_cisMotifs }\SpecialCharTok{\%\textgreater{}\%}
  \FunctionTok{group\_by}\NormalTok{(STRAIN.Fitness, CLASS, MUTATION.Fitness) }\SpecialCharTok{\%\textgreater{}\%}
  \FunctionTok{summarise}\NormalTok{(}\AttributeTok{variance =} \FunctionTok{var}\NormalTok{(meanRelativeFitness))}
\end{Highlighting}
\end{Shaded}

\begin{verbatim}
## `summarise()` has grouped output by 'STRAIN.Fitness', 'CLASS'. You can override
## using the `.groups` argument.
\end{verbatim}

\begin{Shaded}
\begin{Highlighting}[]
\FunctionTok{shapiro.test}\NormalTok{(}\FunctionTok{subset}\NormalTok{(cisMut\_varFitness, CLASS }\SpecialCharTok{==} \StringTok{"TFBS"}\NormalTok{)}\SpecialCharTok{$}\NormalTok{variance)}
\end{Highlighting}
\end{Shaded}

\begin{verbatim}
## 
##  Shapiro-Wilk normality test
## 
## data:  subset(cisMut_varFitness, CLASS == "TFBS")$variance
## W = 0.88864, p-value = 0.2676
\end{verbatim}

\begin{Shaded}
\begin{Highlighting}[]
\FunctionTok{shapiro.test}\NormalTok{(}\FunctionTok{subset}\NormalTok{(cisMut\_varFitness, CLASS }\SpecialCharTok{==} \StringTok{"TATA"}\NormalTok{)}\SpecialCharTok{$}\NormalTok{variance)}
\end{Highlighting}
\end{Shaded}

\begin{verbatim}
## 
##  Shapiro-Wilk normality test
## 
## data:  subset(cisMut_varFitness, CLASS == "TATA")$variance
## W = 0.76193, p-value = 0.01688
\end{verbatim}

\begin{Shaded}
\begin{Highlighting}[]
\CommentTok{\# The Shapiro test suggests that t{-}tests are not appropriate for comparing variation }
\CommentTok{\# in fitness between TFBS and TATA mutants. The distribution of TFBS mutants is sufficiently}
\CommentTok{\# normally distributed but the distribution of TATA box mutations is not. }
\CommentTok{\# We will use a Wilcoxon test to compare the variance in fitness between TFBS and TATA mutants}

\FunctionTok{wilcox.test}\NormalTok{(variance }\SpecialCharTok{\textasciitilde{}}\NormalTok{ CLASS, }\AttributeTok{data =}\NormalTok{ cisMut\_varFitness,}
            \AttributeTok{alternative =} \StringTok{"greater"}\NormalTok{)}
\end{Highlighting}
\end{Shaded}

\begin{verbatim}
## 
##  Wilcoxon rank sum exact test
## 
## data:  variance by CLASS
## W = 32, p-value = 0.1914
## alternative hypothesis: true location shift is greater than 0
\end{verbatim}

\begin{Shaded}
\begin{Highlighting}[]
\DocumentationTok{\#\#\# Make plots showing mutational effects on expression and fitness across environments}
\DocumentationTok{\#\# Reorder factors for plotting aesthetics {-}{-}{-}{-}{-}{-}}
\NormalTok{strain\_order }\OtherTok{\textless{}{-}} \FunctionTok{c}\NormalTok{(}\StringTok{"Y1896"}\NormalTok{, }\StringTok{"Y1897"}\NormalTok{, }\StringTok{"Y1898"}\NormalTok{, }\StringTok{"Y1960"}\NormalTok{, }\StringTok{"Y1892"}\NormalTok{, }\StringTok{"Y1901"}\NormalTok{, }\StringTok{"Y1903"}\NormalTok{, }\CommentTok{\#TFBS }
                  \StringTok{"Y2656"}\NormalTok{, }\StringTok{"Y2550"}\NormalTok{, }\StringTok{"Y2651"}\NormalTok{, }\StringTok{"Y2528"}\NormalTok{, }\StringTok{"Y2566"}\NormalTok{, }\StringTok{"Y2546"}\NormalTok{, }\StringTok{"Y2605"}\NormalTok{)}

\NormalTok{FitnessData\_cisMotifs}\SpecialCharTok{$}\NormalTok{STRAIN.Expression }\OtherTok{\textless{}{-}} \FunctionTok{fct\_relevel}\NormalTok{(FitnessData\_cisMotifs}\SpecialCharTok{$}\NormalTok{STRAIN.Expression, strain\_order)}
\NormalTok{summary\_FitnessData\_cisMotifs}\SpecialCharTok{$}\NormalTok{ENVIRONMENT }\OtherTok{\textless{}{-}} \FunctionTok{fct\_relevel}\NormalTok{(summary\_FitnessData\_cisMotifs}\SpecialCharTok{$}\NormalTok{ENVIRONMENT, }\FunctionTok{c}\NormalTok{(}\StringTok{"GLUCOSE"}\NormalTok{, }\StringTok{"GALACTOSE"}\NormalTok{, }\StringTok{"GLYCEROL"}\NormalTok{, }\StringTok{"ETHANOL"}\NormalTok{))}
\NormalTok{summary\_FitnessData\_cisMotifs}\SpecialCharTok{$}\NormalTok{CLASS }\OtherTok{\textless{}{-}} \FunctionTok{fct\_relevel}\NormalTok{(summary\_FitnessData\_cisMotifs}\SpecialCharTok{$}\NormalTok{CLASS, }\FunctionTok{c}\NormalTok{(}\StringTok{"TFBS"}\NormalTok{, }\StringTok{"TATA"}\NormalTok{))}


\NormalTok{Expression\_cisMotifs}\SpecialCharTok{$}\NormalTok{ENVIRONMENT }\OtherTok{\textless{}{-}} \FunctionTok{fct\_relevel}\NormalTok{(Expression\_cisMotifs}\SpecialCharTok{$}\NormalTok{ENVIRONMENT, }\FunctionTok{c}\NormalTok{(}\StringTok{"GLUCOSE"}\NormalTok{, }\StringTok{"GALACTOSE"}\NormalTok{, }\StringTok{"GLYCEROL"}\NormalTok{, }\StringTok{"ETHANOL"}\NormalTok{))}
\NormalTok{Expression\_cisMotifs}\SpecialCharTok{$}\NormalTok{CLASS }\OtherTok{\textless{}{-}} \FunctionTok{fct\_relevel}\NormalTok{(Expression\_cisMotifs}\SpecialCharTok{$}\NormalTok{CLASS, }\FunctionTok{c}\NormalTok{(}\StringTok{"TFBS"}\NormalTok{, }\StringTok{"TATA"}\NormalTok{))}
\NormalTok{cisMut\_varFitness}\SpecialCharTok{$}\NormalTok{CLASS }\OtherTok{\textless{}{-}} \FunctionTok{fct\_relevel}\NormalTok{(cisMotifs\_var}\SpecialCharTok{$}\NormalTok{CLASS, }\FunctionTok{c}\NormalTok{(}\StringTok{"TFBS"}\NormalTok{, }\StringTok{"TATA"}\NormalTok{))}


\CommentTok{\# For color consistency }
\NormalTok{named\_colors }\OtherTok{\textless{}{-}} \FunctionTok{c}\NormalTok{(}\StringTok{"GLUCOSE"} \OtherTok{=} \StringTok{"\#0072B2"}\NormalTok{, }\StringTok{"GALACTOSE"} \OtherTok{=} \StringTok{"\#009E73"}\NormalTok{, }\StringTok{"GLYCEROL"} \OtherTok{=} \StringTok{"\#CC79A7"}\NormalTok{, }\StringTok{"ETHANOL"} \OtherTok{=} \StringTok{"\#D55E00"}\NormalTok{)}
\NormalTok{motif\_colors }\OtherTok{\textless{}{-}} \FunctionTok{c}\NormalTok{(}\StringTok{"m63"} \OtherTok{=} \StringTok{"\#FBB913"}\NormalTok{, }\StringTok{"m66"} \OtherTok{=} \StringTok{"\#FBB913"}\NormalTok{,}
                  \StringTok{"m75"} \OtherTok{=} \StringTok{"\#6DC8BF"}\NormalTok{, }\StringTok{"m76"} \OtherTok{=} \StringTok{"\#6DC8BF"}\NormalTok{,}
                  \StringTok{"m89"} \OtherTok{=} \StringTok{"\#B72467"}\NormalTok{, }\StringTok{"m90"} \OtherTok{=}\StringTok{"\#B72467"}\NormalTok{, }\StringTok{"m91"} \OtherTok{=} \StringTok{"\#B72467"}\NormalTok{) }\CommentTok{\#RAP1, GCR1a, GCR1b, TATA colors}

\DocumentationTok{\#\# Plots{-}{-}expression effects of TFBS vs TATA {-}{-}{-}{-}{-}}
\NormalTok{plot\_expTFBSvTATA\_linkedFacet }\OtherTok{\textless{}{-}} \FunctionTok{ggplot}\NormalTok{(Expression\_cisMotifs, }\FunctionTok{aes}\NormalTok{(}\AttributeTok{x =}\NormalTok{ ENVIRONMENT, }\AttributeTok{y =}\NormalTok{ YFP.MEDIAN.RELATIVE.MEAN)) }\SpecialCharTok{+} 
  \FunctionTok{geom\_line}\NormalTok{(}\FunctionTok{aes}\NormalTok{(}\AttributeTok{group =}\NormalTok{ STRAIN.Expression), }\AttributeTok{alpha =} \FloatTok{0.5}\NormalTok{) }\SpecialCharTok{+}  
  \FunctionTok{geom\_errorbar}\NormalTok{(}\FunctionTok{aes}\NormalTok{(}\AttributeTok{ymin =}\NormalTok{ YFP.MEDIAN.RELATIVE.MEAN }\SpecialCharTok{{-}}\NormalTok{ YFP.MEDIAN.RELATIVE.SD, }
                    \AttributeTok{ymax =}\NormalTok{ YFP.MEDIAN.RELATIVE.MEAN }\SpecialCharTok{+}\NormalTok{ YFP.MEDIAN.RELATIVE.SD, }\AttributeTok{color =}\NormalTok{ ENVIRONMENT), }
                \AttributeTok{width =} \FloatTok{0.2}\NormalTok{, }\AttributeTok{alpha =} \FloatTok{0.5}\NormalTok{) }\SpecialCharTok{+}
  \FunctionTok{geom\_point}\NormalTok{(}\FunctionTok{aes}\NormalTok{(}\AttributeTok{color =}\NormalTok{ MUTATION.Expression), }\AttributeTok{size =} \FloatTok{2.5}\NormalTok{) }\SpecialCharTok{+} 
  \FunctionTok{theme\_bw}\NormalTok{() }\SpecialCharTok{+} \FunctionTok{labs}\NormalTok{(}\AttributeTok{x =} \StringTok{""}\NormalTok{, }\AttributeTok{y =} \StringTok{"Relative Expression Level to Reference (afu)"}\NormalTok{) }\SpecialCharTok{+}
  \FunctionTok{scale\_color\_manual}\NormalTok{(}\AttributeTok{values =}\NormalTok{ motif\_colors) }\SpecialCharTok{+} \CommentTok{\#scale\_fill\_manual(values = motif\_colors) +}
  \FunctionTok{theme}\NormalTok{(}\AttributeTok{panel.border =} \FunctionTok{element\_rect}\NormalTok{(}\AttributeTok{linetype =} \DecValTok{1}\NormalTok{, }\AttributeTok{fill =} \StringTok{"NA"}\NormalTok{),}
        \AttributeTok{plot.background =} \FunctionTok{element\_rect}\NormalTok{(}\AttributeTok{color =} \ConstantTok{NA}\NormalTok{),}\AttributeTok{legend.position=}\StringTok{"top"}\NormalTok{,}
        \AttributeTok{axis.text=}\FunctionTok{element\_text}\NormalTok{(}\AttributeTok{size=} \DecValTok{10}\NormalTok{), }\AttributeTok{axis.title=}\FunctionTok{element\_text}\NormalTok{(}\AttributeTok{size =}\DecValTok{14}\NormalTok{),}
        \AttributeTok{axis.text.x=}\FunctionTok{element\_text}\NormalTok{(}\AttributeTok{angle=}\DecValTok{45}\NormalTok{,}\AttributeTok{hjust=}\DecValTok{1}\NormalTok{),}
        \AttributeTok{panel.grid.minor =} \FunctionTok{element\_blank}\NormalTok{(),}
        \AttributeTok{panel.grid.major =} \FunctionTok{element\_blank}\NormalTok{()) }\SpecialCharTok{+} 
  \FunctionTok{facet\_wrap}\NormalTok{(}\SpecialCharTok{\textasciitilde{}}\NormalTok{CLASS, }\AttributeTok{ncol =} \DecValTok{2}\NormalTok{, }\AttributeTok{scales =} \StringTok{"free\_x"}\NormalTok{,}\AttributeTok{strip.position =} \StringTok{"top"}\NormalTok{) }

\CommentTok{\# Plot the variation between TFBS and TATA mutants}
\NormalTok{plot\_expTFBSvTATA\_variance }\OtherTok{\textless{}{-}} \FunctionTok{ggplot}\NormalTok{(cisMotifs\_var, }\FunctionTok{aes}\NormalTok{(}\AttributeTok{x =}\NormalTok{ CLASS, }\AttributeTok{y =}\NormalTok{ variance)) }\SpecialCharTok{+}
  \FunctionTok{geom\_boxplot}\NormalTok{(}\AttributeTok{outlier.shape =} \ConstantTok{NA}\NormalTok{, }\AttributeTok{width =} \FloatTok{0.5}\NormalTok{) }\SpecialCharTok{+}
  \FunctionTok{geom\_jitter}\NormalTok{(}\FunctionTok{aes}\NormalTok{(}\AttributeTok{col =}\NormalTok{ MUTATION.Expression), }\AttributeTok{width =} \FloatTok{0.15}\NormalTok{, }\AttributeTok{size =} \FloatTok{2.5}\NormalTok{) }\SpecialCharTok{+}
  \FunctionTok{scale\_color\_manual}\NormalTok{(}\AttributeTok{values =}\NormalTok{ motif\_colors) }\SpecialCharTok{+} \CommentTok{\#scale\_fill\_manual(values = named\_colors) +}
  \FunctionTok{theme\_bw}\NormalTok{() }\SpecialCharTok{+} \FunctionTok{labs}\NormalTok{(}\AttributeTok{x =} \StringTok{""}\NormalTok{, }\AttributeTok{y =} \StringTok{"Expression Effect Variance"}\NormalTok{) }\SpecialCharTok{+}
  \FunctionTok{scale\_fill\_manual}\NormalTok{(}\AttributeTok{values =}\NormalTok{ named\_colors) }\SpecialCharTok{+} 
  \FunctionTok{theme}\NormalTok{(}\AttributeTok{panel.border =} \FunctionTok{element\_rect}\NormalTok{(}\AttributeTok{linetype =} \DecValTok{1}\NormalTok{, }\AttributeTok{fill =} \StringTok{"NA"}\NormalTok{),}
        \AttributeTok{plot.background =} \FunctionTok{element\_rect}\NormalTok{(}\AttributeTok{color =} \ConstantTok{NA}\NormalTok{),}\AttributeTok{legend.position=}\StringTok{"none"}\NormalTok{,}
        \AttributeTok{axis.text=}\FunctionTok{element\_text}\NormalTok{(}\AttributeTok{size=} \DecValTok{10}\NormalTok{), }\AttributeTok{axis.title=}\FunctionTok{element\_text}\NormalTok{(}\AttributeTok{size =}\DecValTok{14}\NormalTok{),}
        \AttributeTok{axis.text.x=}\FunctionTok{element\_text}\NormalTok{(}\AttributeTok{angle=}\DecValTok{45}\NormalTok{,}\AttributeTok{hjust=}\DecValTok{1}\NormalTok{),}
        \AttributeTok{panel.grid.minor =} \FunctionTok{element\_blank}\NormalTok{(),}
        \AttributeTok{panel.grid.major =} \FunctionTok{element\_blank}\NormalTok{())}

\DocumentationTok{\#\# Plots{-}{-}fitness effects of TFBS vs TATA {-}{-}{-}{-}{-}}
\NormalTok{plot\_fitTFBSvTATA\_linkedFacet }\OtherTok{\textless{}{-}} \FunctionTok{ggplot}\NormalTok{(summary\_FitnessData\_cisMotifs, }
                                        \FunctionTok{aes}\NormalTok{(}\AttributeTok{x =}\NormalTok{ ENVIRONMENT, }\AttributeTok{y =}\NormalTok{ meanRelativeFitness)) }\SpecialCharTok{+} 
  \FunctionTok{geom\_line}\NormalTok{(}\FunctionTok{aes}\NormalTok{(}\AttributeTok{group =}\NormalTok{ STRAIN.Fitness)) }\SpecialCharTok{+}
  \FunctionTok{geom\_errorbar}\NormalTok{(}\FunctionTok{aes}\NormalTok{(}\AttributeTok{ymin =}\NormalTok{ meanRelativeFitness }\SpecialCharTok{{-}}\NormalTok{ sdRelativeFitness, }
                    \AttributeTok{ymax =}\NormalTok{ meanRelativeFitness }\SpecialCharTok{+}\NormalTok{ sdRelativeFitness, }\AttributeTok{color =}\NormalTok{ ENVIRONMENT), }\AttributeTok{width =} \FloatTok{0.2}\NormalTok{, }\AttributeTok{alpha =} \FloatTok{0.5}\NormalTok{) }\SpecialCharTok{+}
  \FunctionTok{geom\_point}\NormalTok{(}\FunctionTok{aes}\NormalTok{(}\AttributeTok{color =}\NormalTok{ MUTATION.Fitness), }\AttributeTok{size =} \FloatTok{2.5}\NormalTok{) }\SpecialCharTok{+} 
  \FunctionTok{theme\_bw}\NormalTok{() }\SpecialCharTok{+} \FunctionTok{labs}\NormalTok{(}\AttributeTok{x =} \StringTok{""}\NormalTok{, }\AttributeTok{y =} \StringTok{"Relative Fitness"}\NormalTok{) }\SpecialCharTok{+}
  \FunctionTok{scale\_color\_manual}\NormalTok{(}\AttributeTok{values =}\NormalTok{ motif\_colors) }\SpecialCharTok{+} \CommentTok{\#scale\_fill\_manual(values = named\_colors) +}
  \FunctionTok{theme}\NormalTok{(}\AttributeTok{panel.border =} \FunctionTok{element\_rect}\NormalTok{(}\AttributeTok{linetype =} \DecValTok{1}\NormalTok{, }\AttributeTok{fill =} \StringTok{"NA"}\NormalTok{),}
        \AttributeTok{plot.background =} \FunctionTok{element\_rect}\NormalTok{(}\AttributeTok{color =} \ConstantTok{NA}\NormalTok{),}\AttributeTok{legend.position=}\StringTok{"top"}\NormalTok{,}
        \AttributeTok{axis.text=}\FunctionTok{element\_text}\NormalTok{(}\AttributeTok{size=} \DecValTok{10}\NormalTok{), }\AttributeTok{axis.title=}\FunctionTok{element\_text}\NormalTok{(}\AttributeTok{size =}\DecValTok{14}\NormalTok{),}
        \AttributeTok{axis.text.x=}\FunctionTok{element\_text}\NormalTok{(}\AttributeTok{angle=}\DecValTok{45}\NormalTok{,}\AttributeTok{hjust=}\DecValTok{1}\NormalTok{),}
        \AttributeTok{panel.grid.minor =} \FunctionTok{element\_blank}\NormalTok{(),}
        \AttributeTok{panel.grid.major =} \FunctionTok{element\_blank}\NormalTok{()) }\SpecialCharTok{+} 
  \FunctionTok{facet\_wrap}\NormalTok{(}\SpecialCharTok{\textasciitilde{}}\NormalTok{CLASS, }\AttributeTok{ncol =} \DecValTok{2}\NormalTok{, }\AttributeTok{scales =} \StringTok{"free\_x"}\NormalTok{,}\AttributeTok{strip.position =} \StringTok{"top"}\NormalTok{)}\SpecialCharTok{+}   
  \FunctionTok{ylim}\NormalTok{(}\FloatTok{0.94}\NormalTok{, }\FloatTok{1.005}\NormalTok{) }

\CommentTok{\# Plot the variation between TFBS and TATA mutants}
\NormalTok{plot\_fitTFBSvTATA\_variance }\OtherTok{\textless{}{-}} \FunctionTok{ggplot}\NormalTok{(cisMut\_varFitness, }\FunctionTok{aes}\NormalTok{(}\AttributeTok{x =}\NormalTok{ CLASS, }\AttributeTok{y =}\NormalTok{ variance)) }\SpecialCharTok{+}
  \FunctionTok{geom\_boxplot}\NormalTok{(}\AttributeTok{outlier.shape =} \ConstantTok{NA}\NormalTok{, }\AttributeTok{width =} \FloatTok{0.5}\NormalTok{) }\SpecialCharTok{+}
  \FunctionTok{geom\_jitter}\NormalTok{(}\FunctionTok{aes}\NormalTok{(}\AttributeTok{col =}\NormalTok{ MUTATION.Fitness), }\AttributeTok{width =} \FloatTok{0.15}\NormalTok{, }\AttributeTok{size =} \FloatTok{2.5}\NormalTok{) }\SpecialCharTok{+}
  \FunctionTok{scale\_color\_manual}\NormalTok{(}\AttributeTok{values =}\NormalTok{ motif\_colors) }\SpecialCharTok{+} \CommentTok{\#scale\_fill\_manual(values = named\_colors) +}
  \FunctionTok{theme\_bw}\NormalTok{() }\SpecialCharTok{+} \FunctionTok{labs}\NormalTok{(}\AttributeTok{x =} \StringTok{""}\NormalTok{, }\AttributeTok{y =} \StringTok{"Fitness Effect Variance"}\NormalTok{) }\SpecialCharTok{+}
  \FunctionTok{scale\_fill\_manual}\NormalTok{(}\AttributeTok{values =}\NormalTok{ named\_colors) }\SpecialCharTok{+} 
  \FunctionTok{theme}\NormalTok{(}\AttributeTok{panel.border =} \FunctionTok{element\_rect}\NormalTok{(}\AttributeTok{linetype =} \DecValTok{1}\NormalTok{, }\AttributeTok{fill =} \StringTok{"NA"}\NormalTok{),}
        \AttributeTok{plot.background =} \FunctionTok{element\_rect}\NormalTok{(}\AttributeTok{color =} \ConstantTok{NA}\NormalTok{),}\AttributeTok{legend.position=}\StringTok{"none"}\NormalTok{,}
        \AttributeTok{axis.text=}\FunctionTok{element\_text}\NormalTok{(}\AttributeTok{size=} \DecValTok{10}\NormalTok{), }\AttributeTok{axis.title=}\FunctionTok{element\_text}\NormalTok{(}\AttributeTok{size =}\DecValTok{14}\NormalTok{),}
        \AttributeTok{axis.text.x=}\FunctionTok{element\_text}\NormalTok{(}\AttributeTok{angle=}\DecValTok{45}\NormalTok{,}\AttributeTok{hjust=}\DecValTok{1}\NormalTok{),}
        \AttributeTok{panel.grid.minor =} \FunctionTok{element\_blank}\NormalTok{(),}
        \AttributeTok{panel.grid.major =} \FunctionTok{element\_blank}\NormalTok{())}

\CommentTok{\#Save the plot}
\FunctionTok{ggsave}\NormalTok{(}\StringTok{"./ManuscriptFigures/Figure4\_plot\_expTFBSvTATA\_linkedFacet.pdf"}\NormalTok{, }\AttributeTok{plot =}\NormalTok{ plot\_expTFBSvTATA\_linkedFacet, }\AttributeTok{width =} \DecValTok{7}\NormalTok{, }\AttributeTok{height =} \DecValTok{5}\NormalTok{)}
\FunctionTok{ggsave}\NormalTok{(}\StringTok{"./ManuscriptFigures/Figure4\_plot\_expTFBSvTATA\_variance.pdf"}\NormalTok{, plot\_expTFBSvTATA\_variance, }\AttributeTok{width =} \FloatTok{2.5}\NormalTok{, }\AttributeTok{height =} \DecValTok{4}\NormalTok{, }\AttributeTok{units =} \StringTok{"in"}\NormalTok{)}
\end{Highlighting}
\end{Shaded}

\begin{verbatim}
## Warning: No shared levels found between `names(values)` of the manual scale and the
## data's fill values.
\end{verbatim}

\begin{Shaded}
\begin{Highlighting}[]
\FunctionTok{ggsave}\NormalTok{(}\StringTok{"./ManuscriptFigures/Figure4\_plot\_fitTFBSvTATA\_linkedFacet.pdf"}\NormalTok{, plot\_fitTFBSvTATA\_linkedFacet, }\AttributeTok{width =} \DecValTok{7}\NormalTok{, }\AttributeTok{height =} \DecValTok{5}\NormalTok{, }\AttributeTok{units =} \StringTok{"in"}\NormalTok{)}
\FunctionTok{ggsave}\NormalTok{(}\StringTok{"./ManuscriptFigures/Figure4\_plot\_fitTFBSvTATA\_variance.pdf"}\NormalTok{, plot\_fitTFBSvTATA\_variance, }\AttributeTok{width =} \FloatTok{2.5}\NormalTok{, }\AttributeTok{height =} \DecValTok{4}\NormalTok{, }\AttributeTok{units =} \StringTok{"in"}\NormalTok{)}
\end{Highlighting}
\end{Shaded}

\begin{verbatim}
## Warning: No shared levels found between `names(values)` of the manual scale and the
## data's fill values.
\end{verbatim}

\end{document}
