## Supplementary figures and images for "Plasticity and environment-specific relationships between gene expression and fitness in *Saccharomyces cerevisiae*"

### Fig2_plot_DeletionEffect_violin.pdf

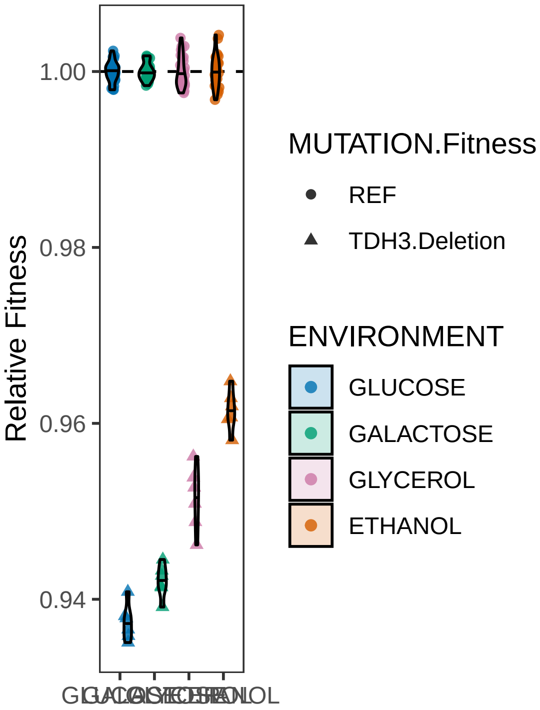

### Fig2_plot_DoublingTime.pdf

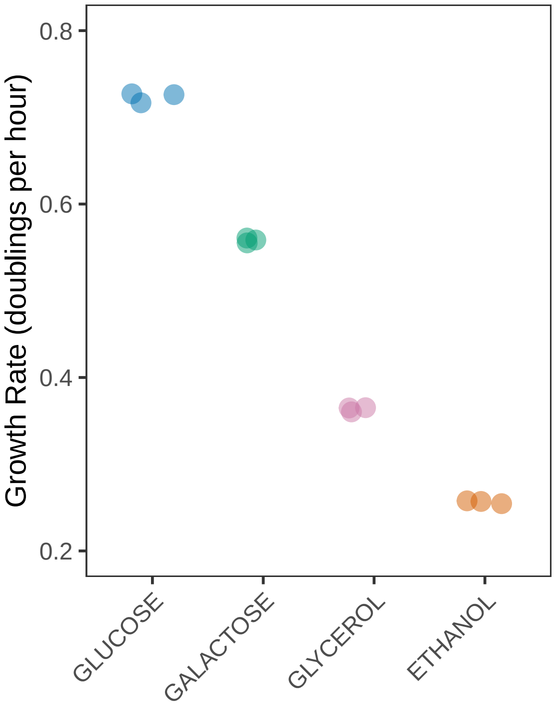

### Fig2_plot_ExpressionReference_violin.pdf

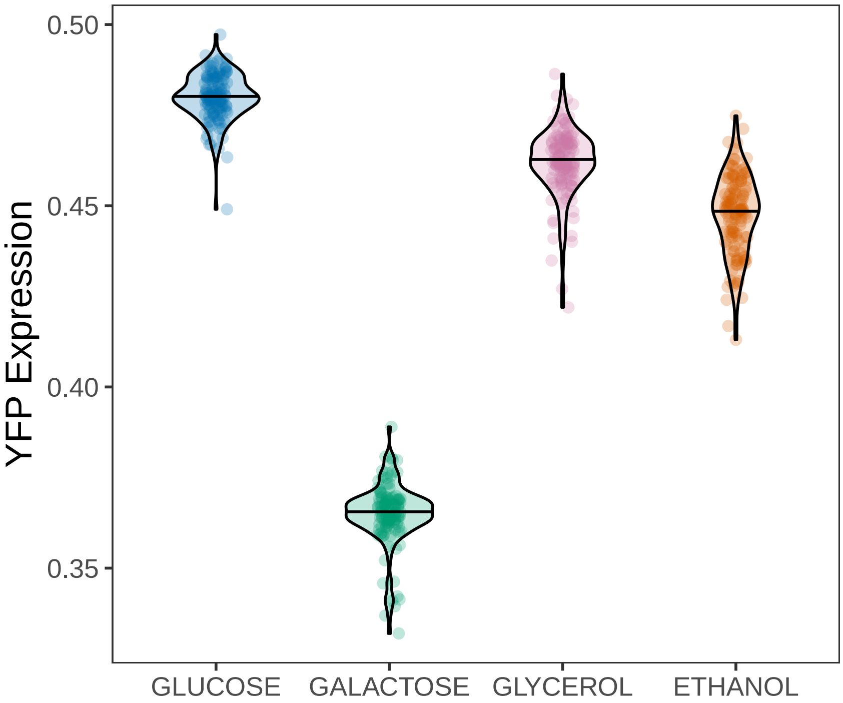

### Fig3_ExpressionEffects.pdf

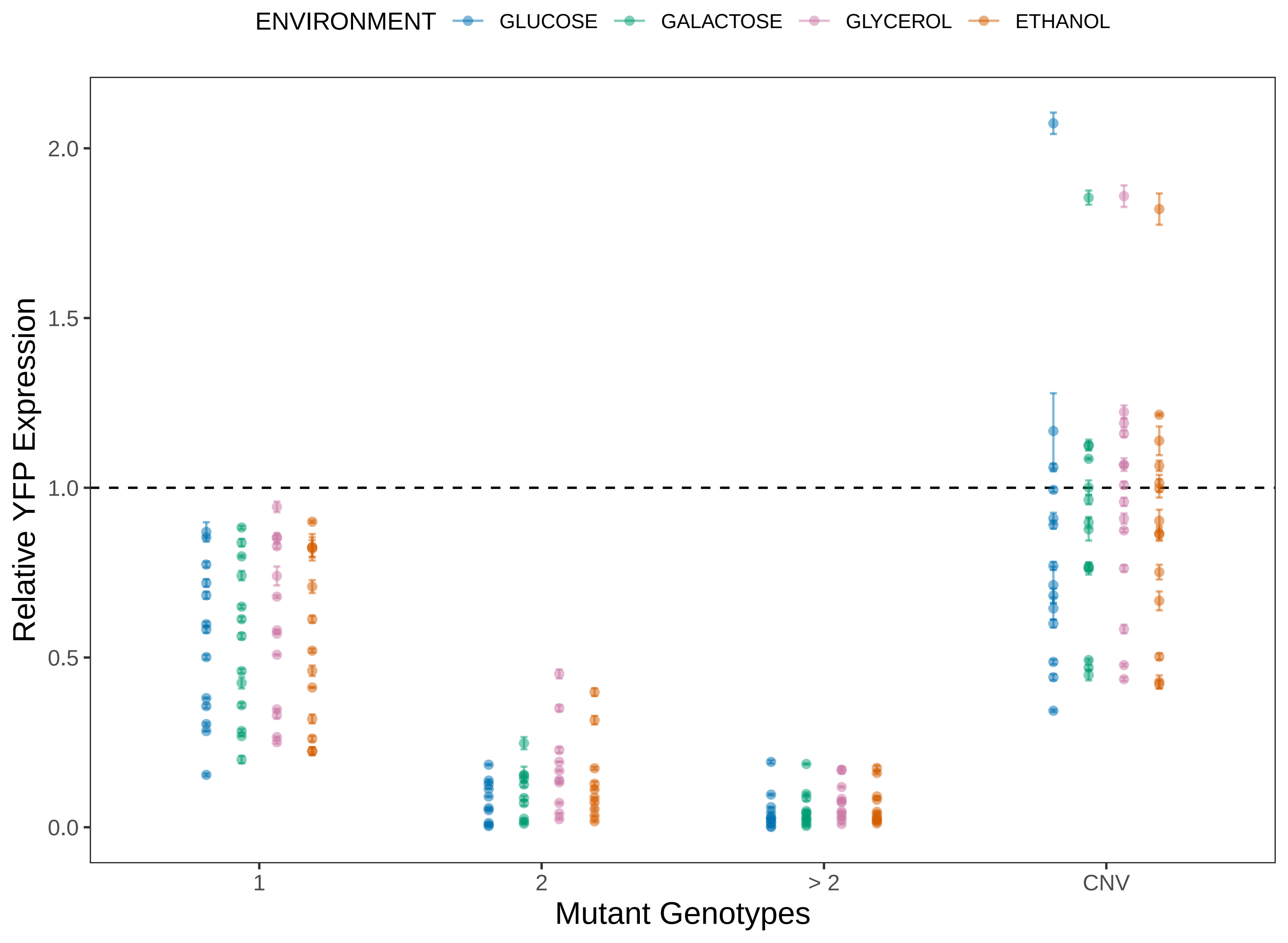

### Fig3_fitnessCurves_facet.pdf

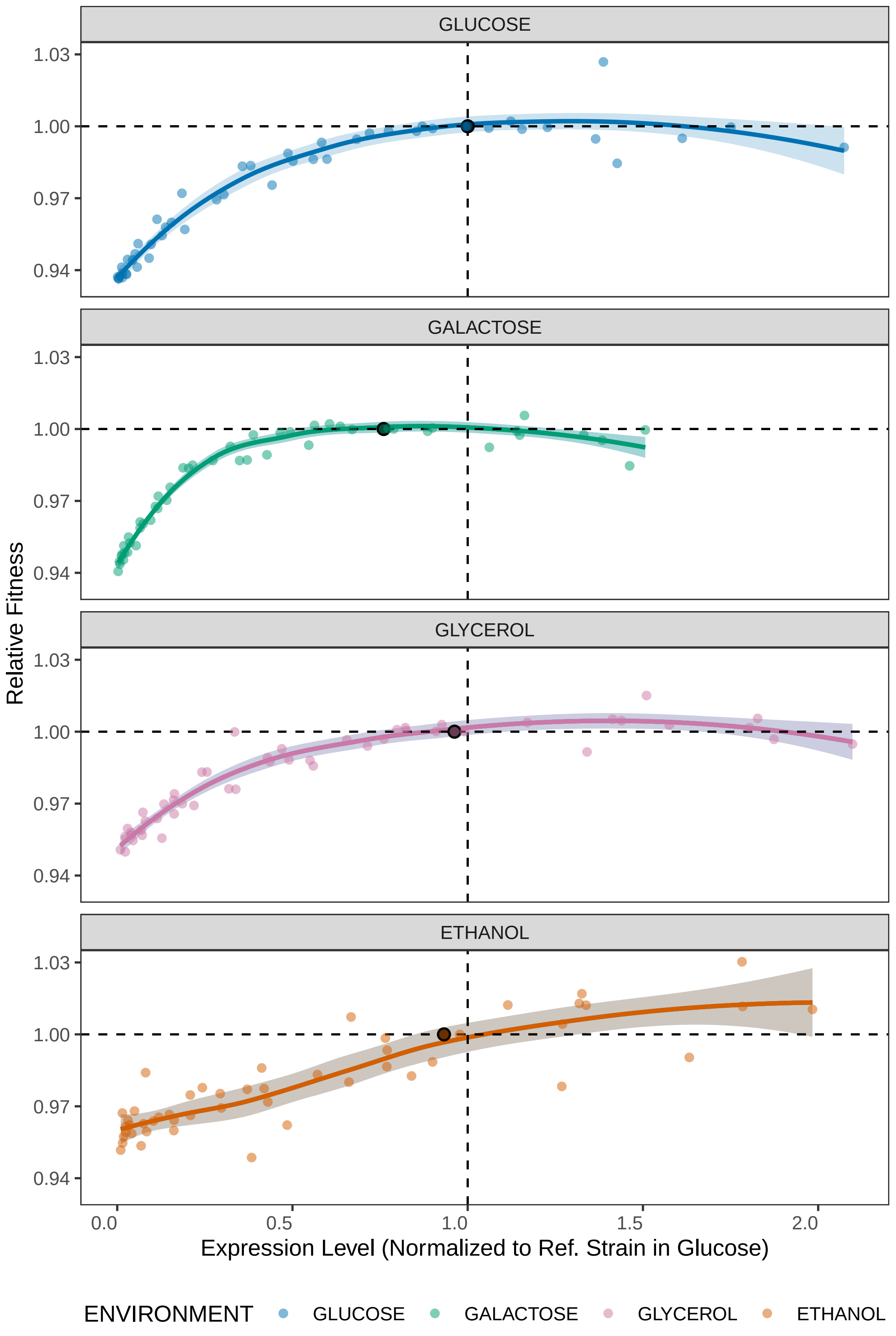

### Fig3_fitnessCurves_overlain.pdf

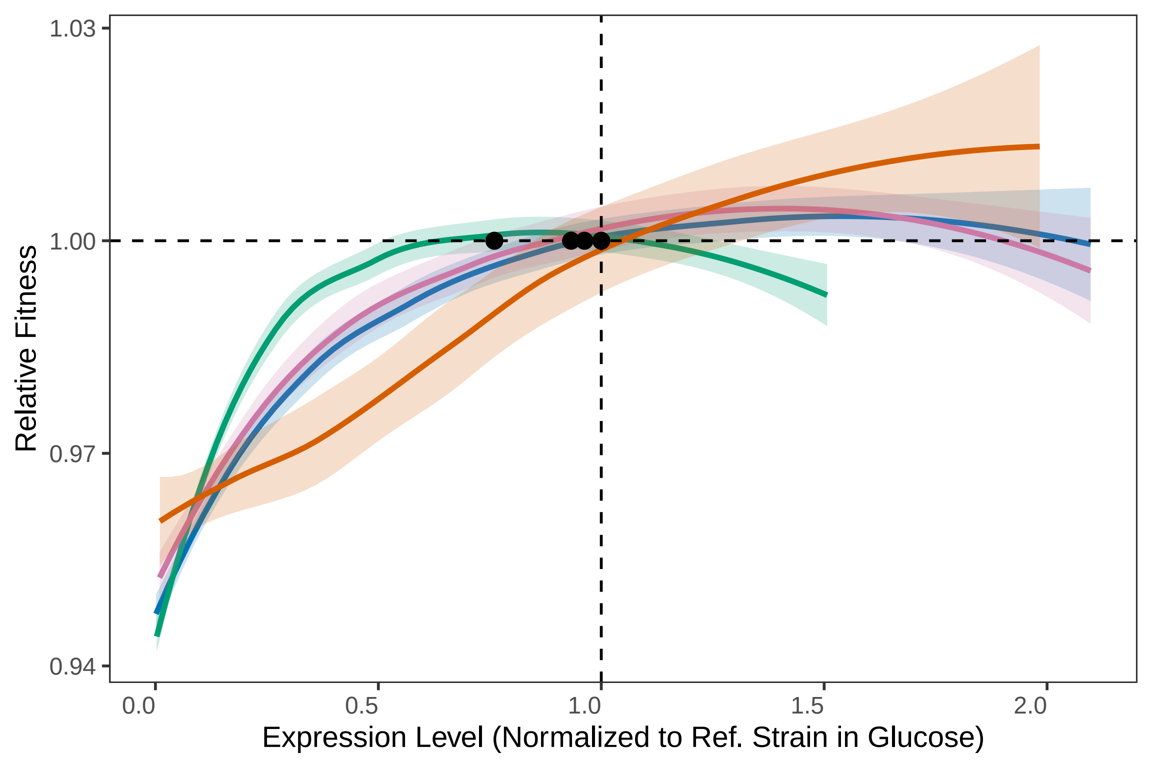

### Fig3_GALvETH.pdf

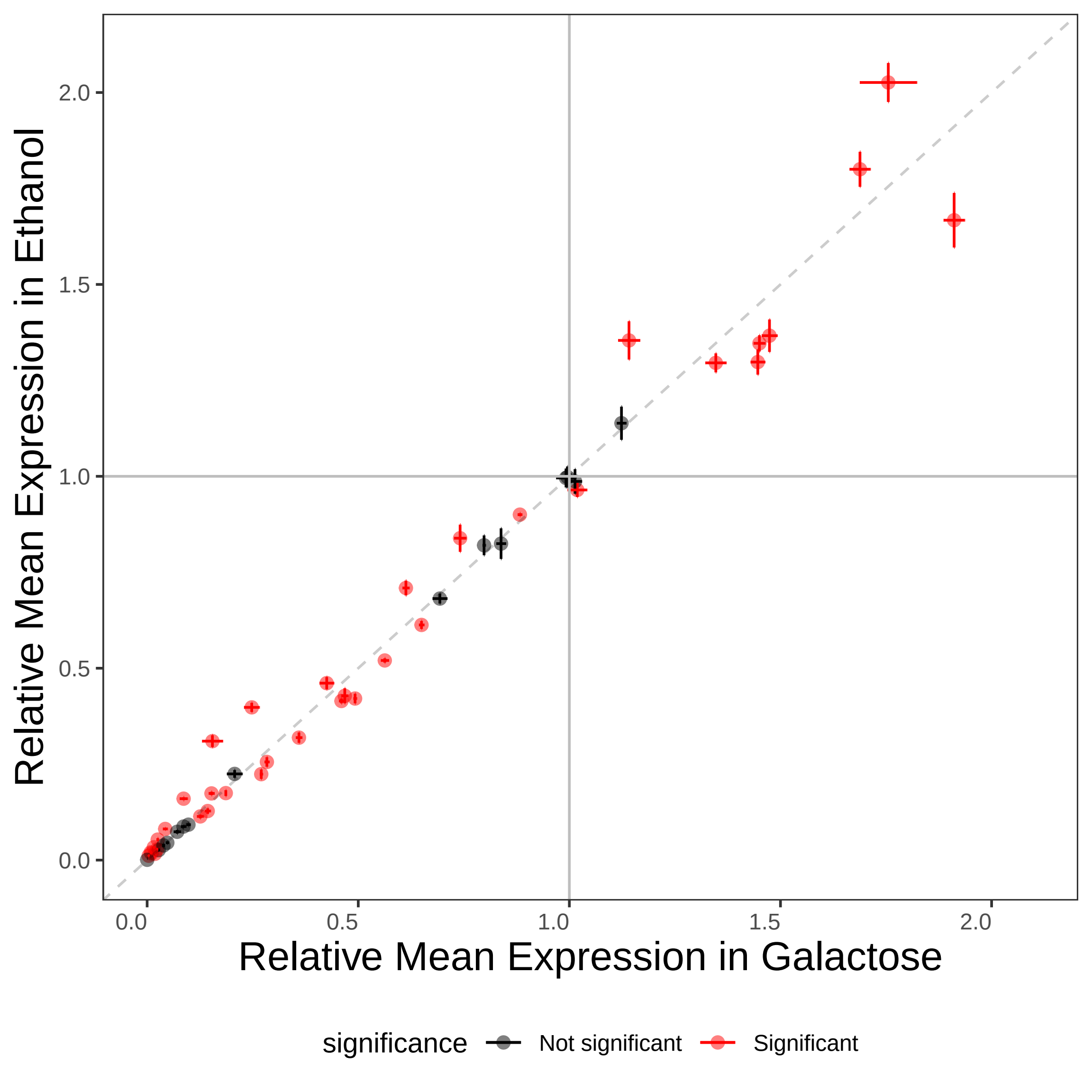

### Fig3_GALvGLY.pdf

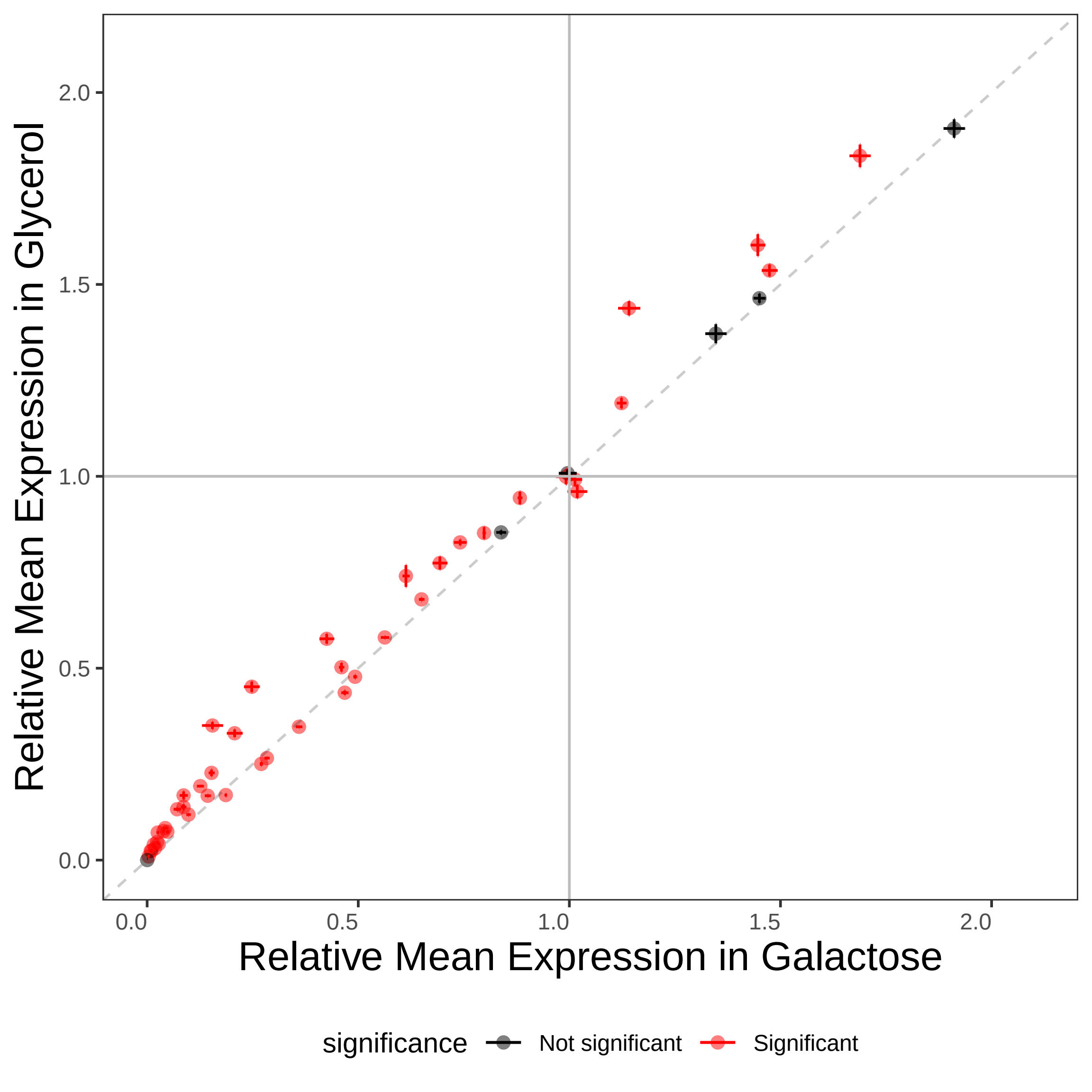

### Fig3_GLUvETH.pdf

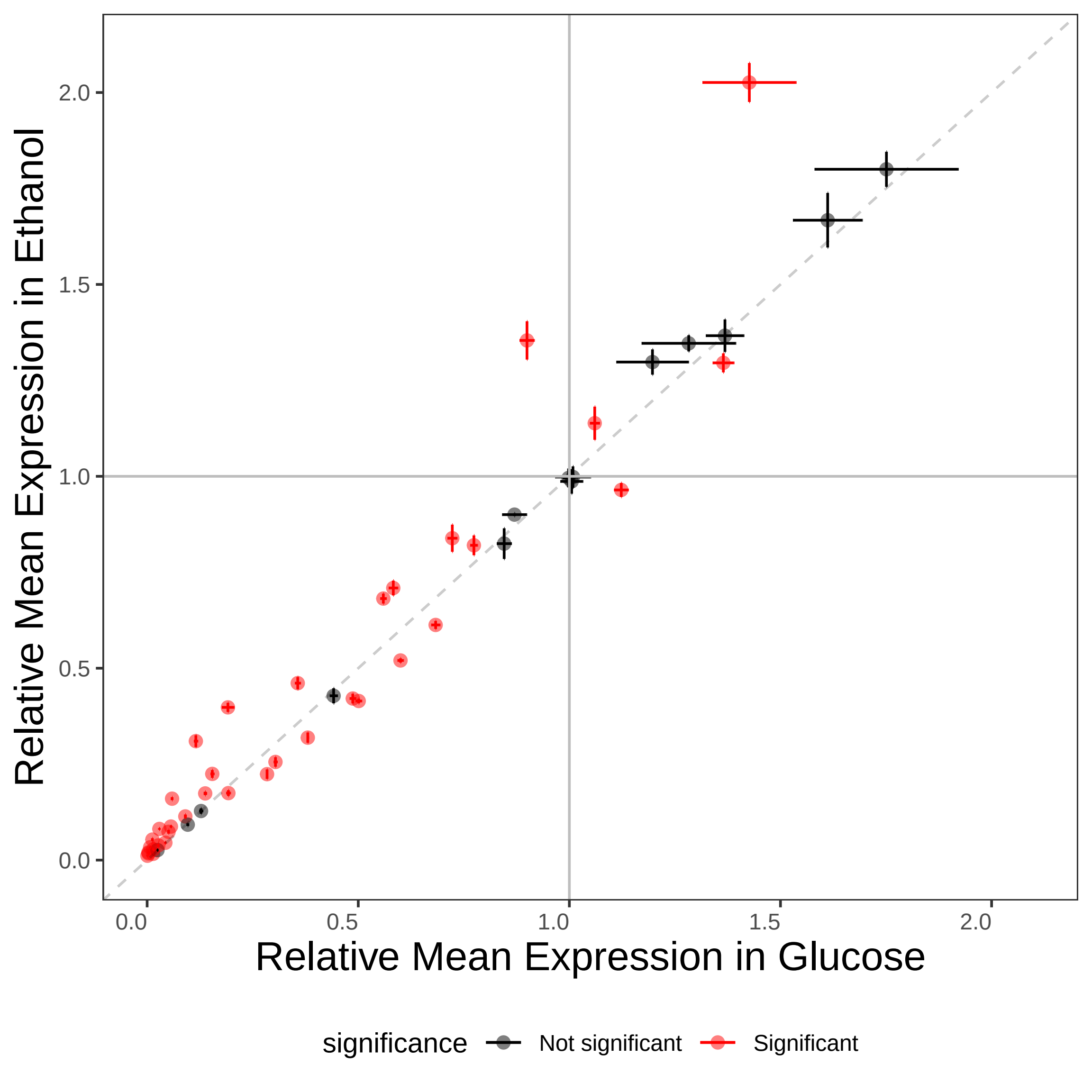

### Fig3_GLUvGAL.pdf

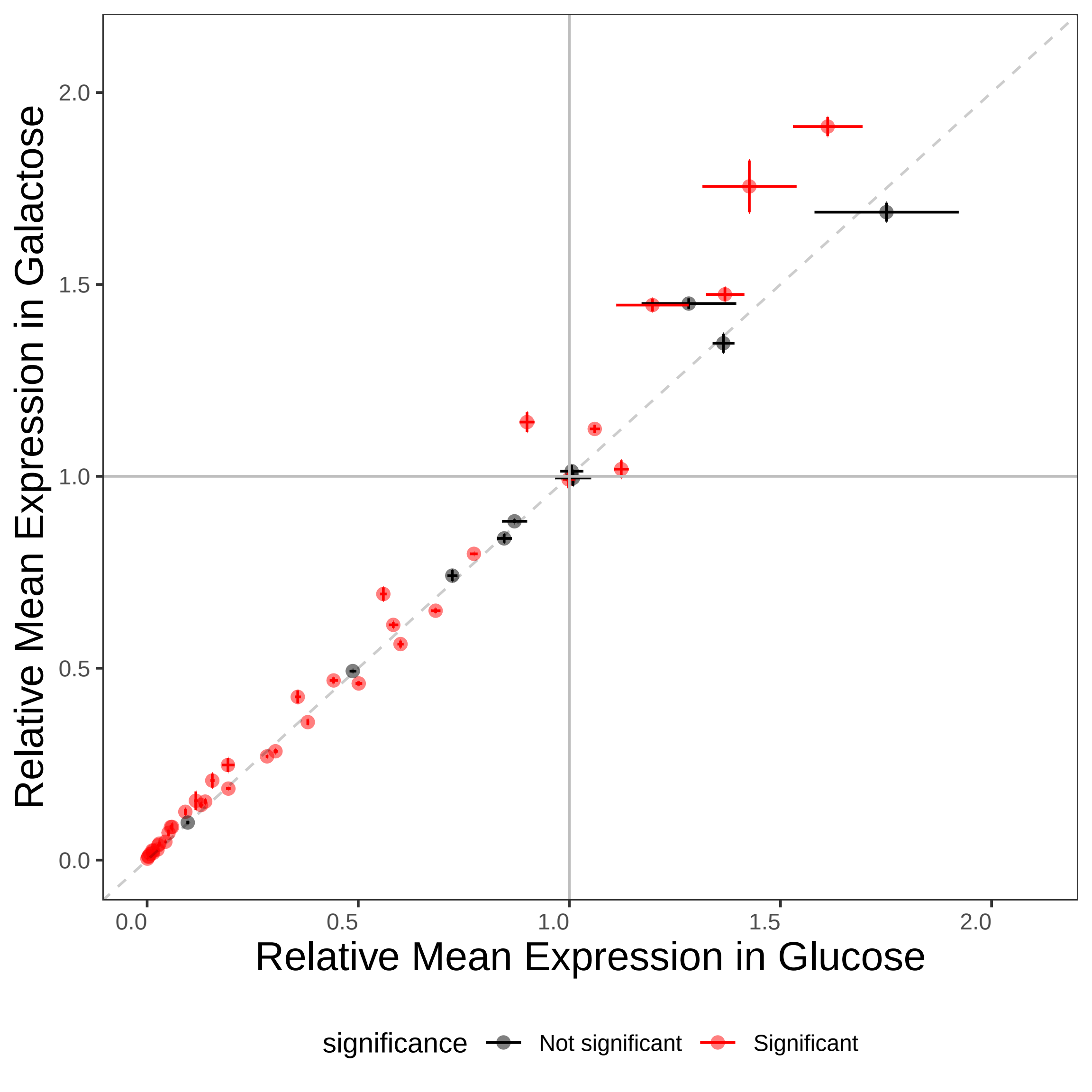

### Fig3_GLUvGLY.pdf

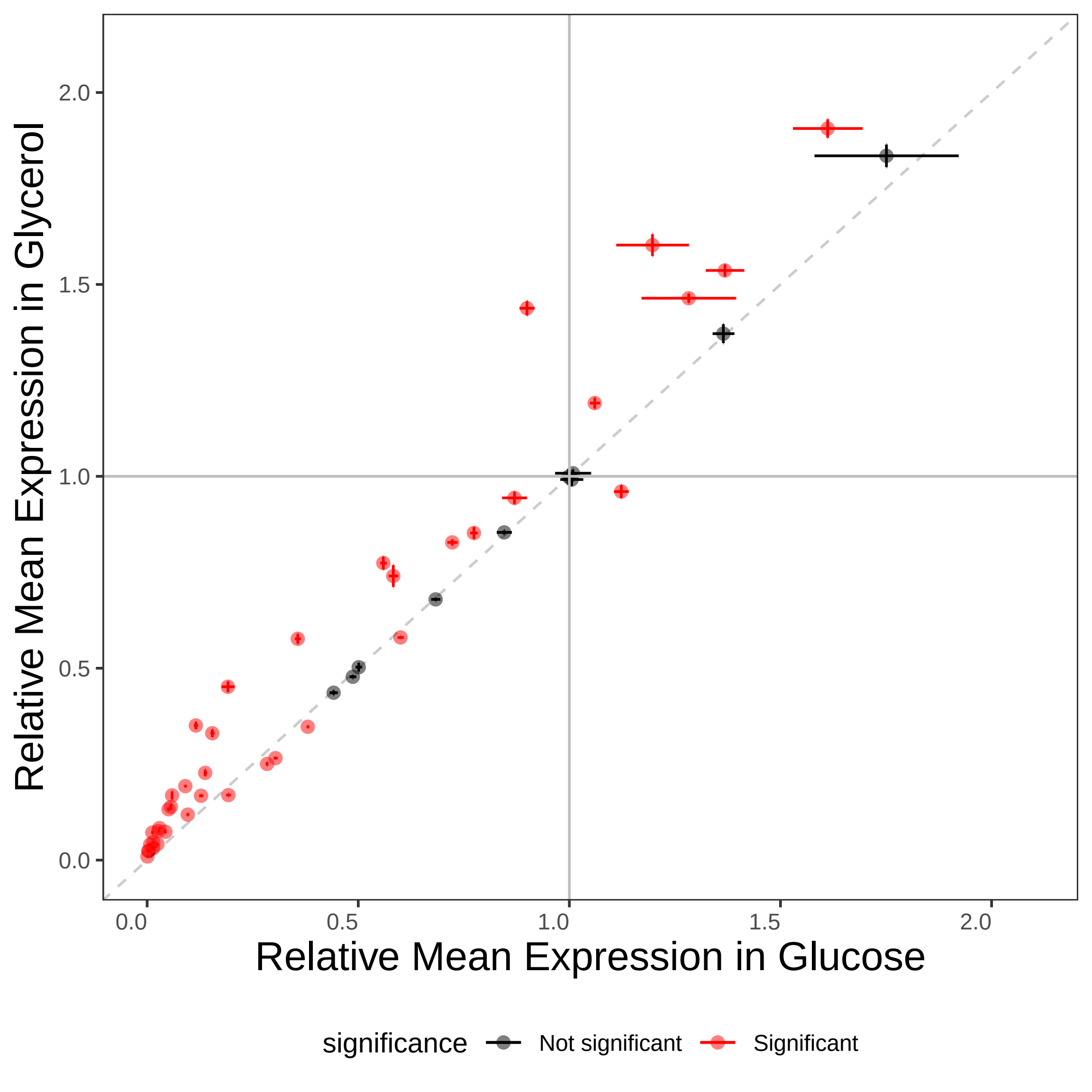

### Fig3_GLYvETH.pdf

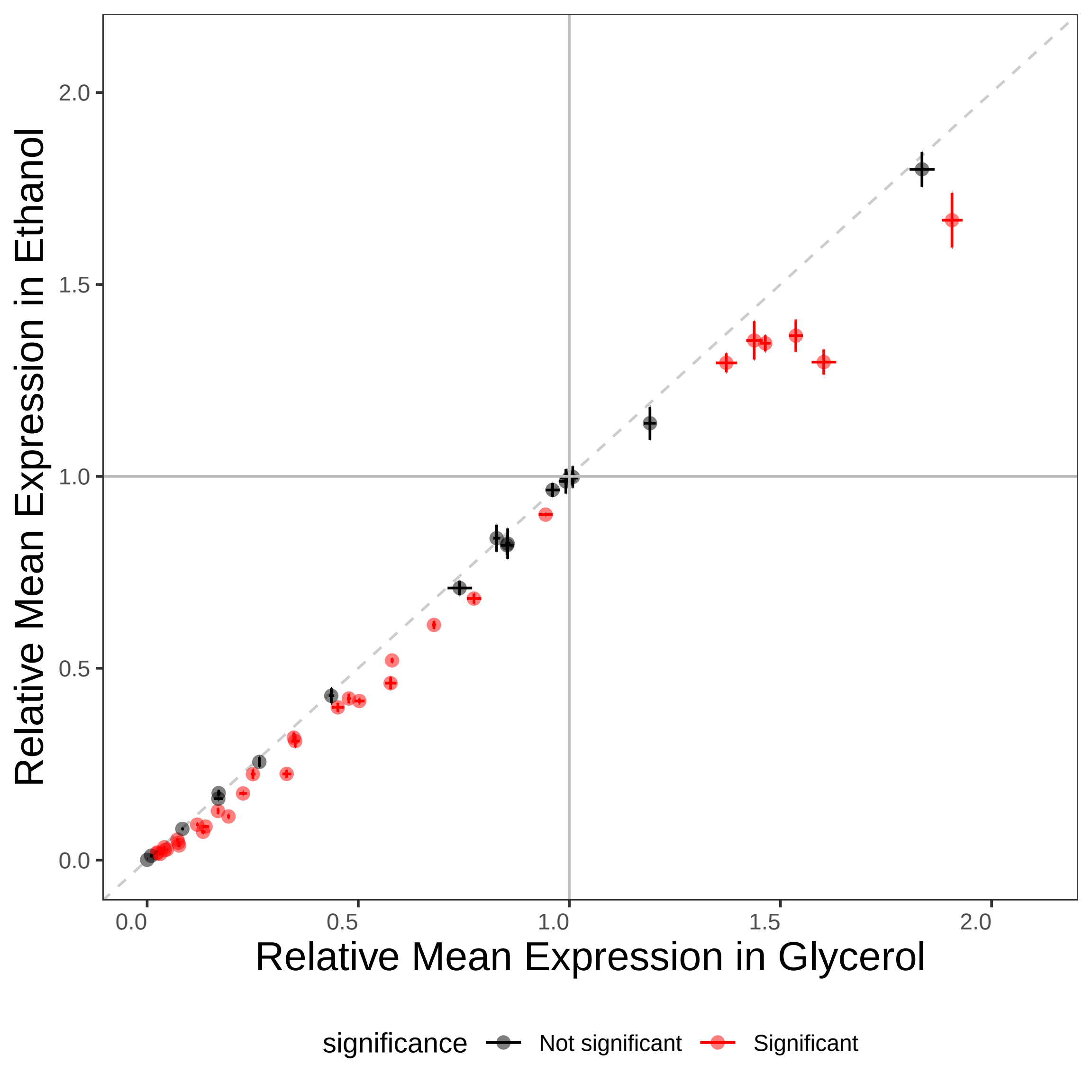

### Figure4_plot_expTFBSvTATA_linkedFacet.pdf

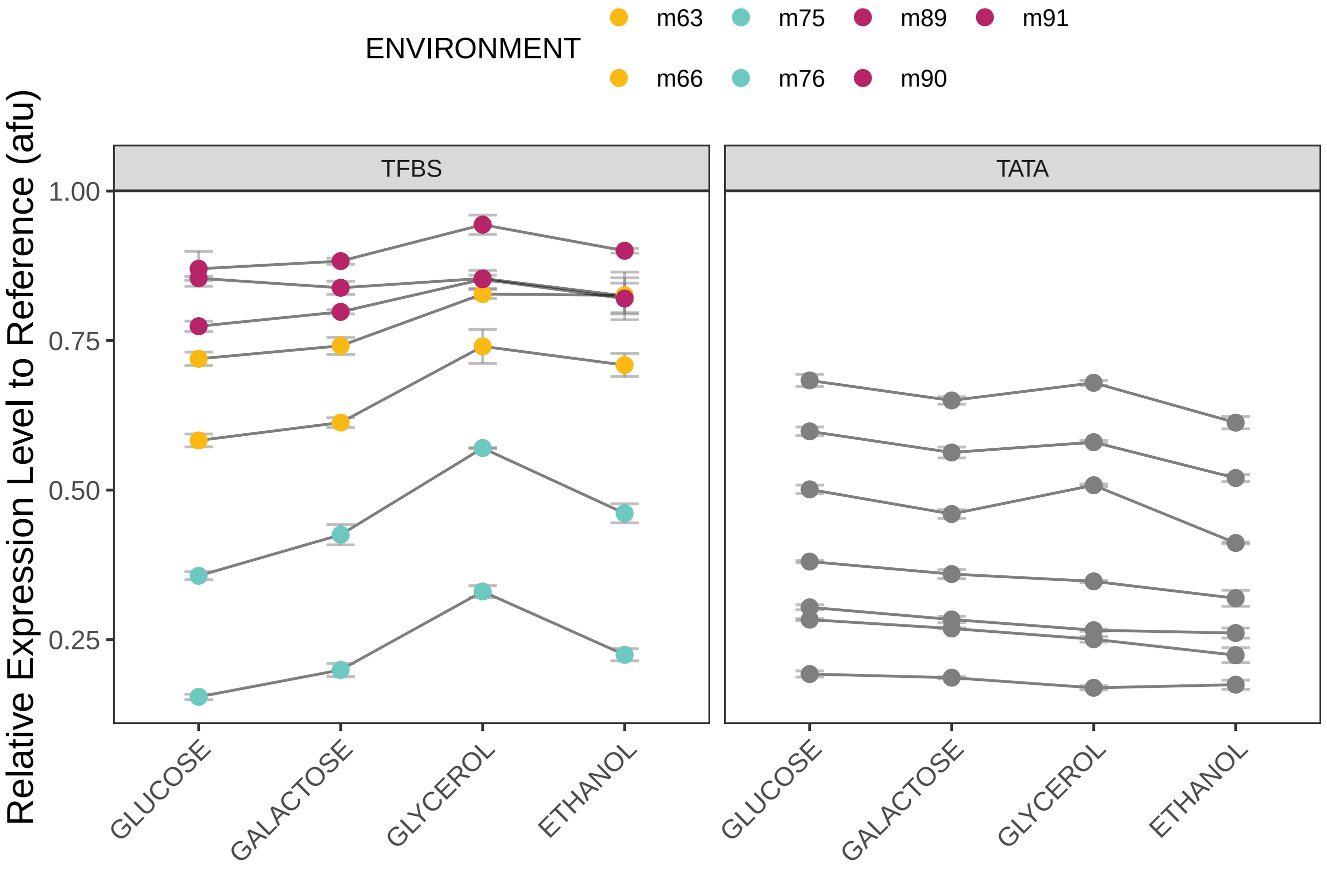

### Figure4_plot_expTFBSvTATA_variance.pdf

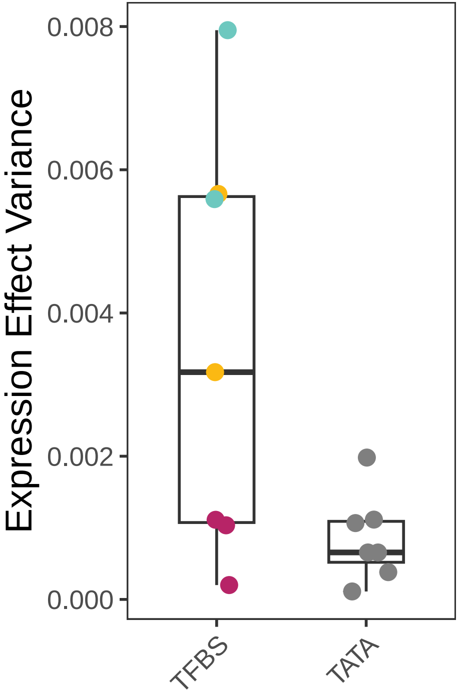

### Figure4_plot_fitTFBSvTATA_linkedFacet.pdf

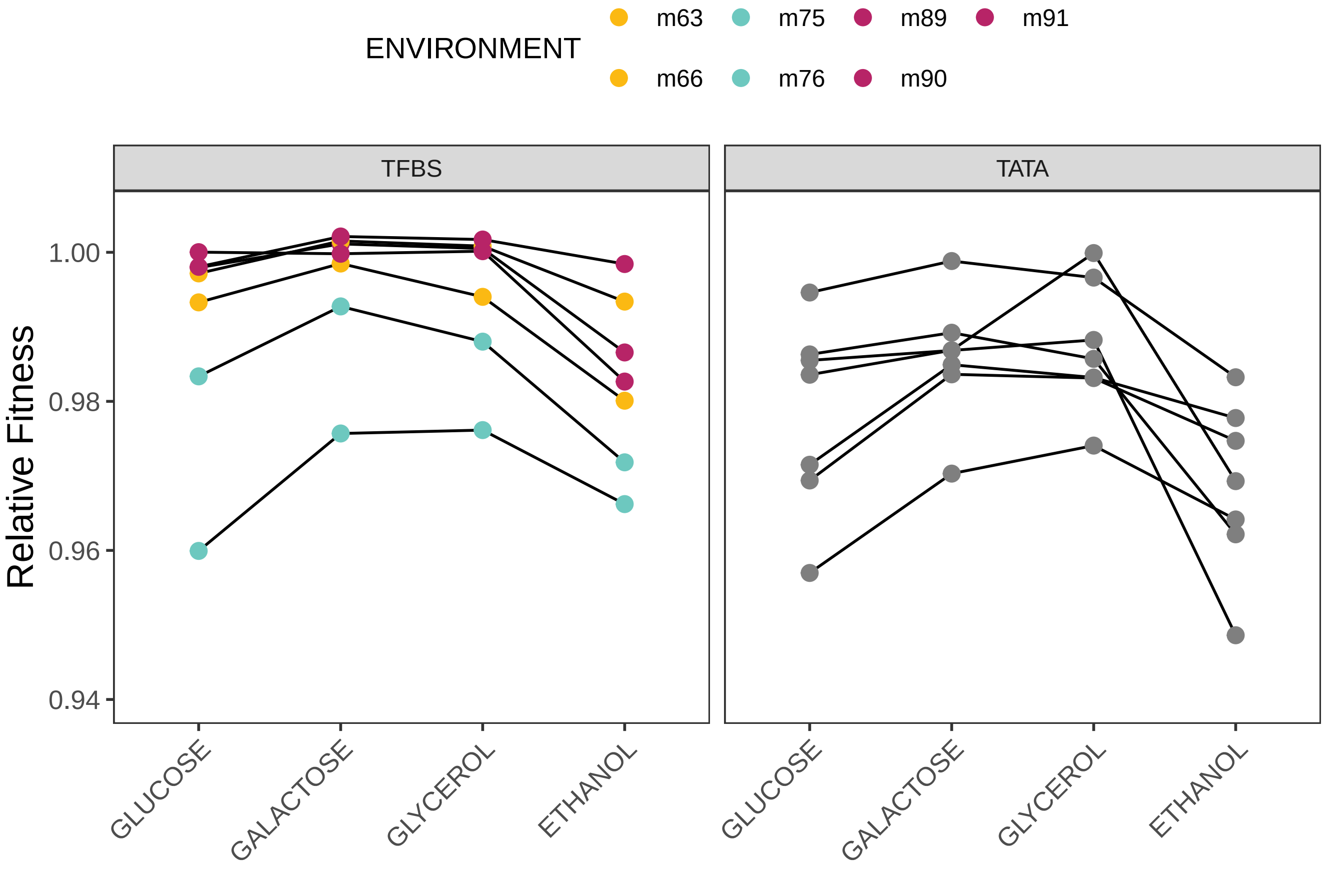

### Figure4_plot_fitTFBSvTATA_variance.pdf

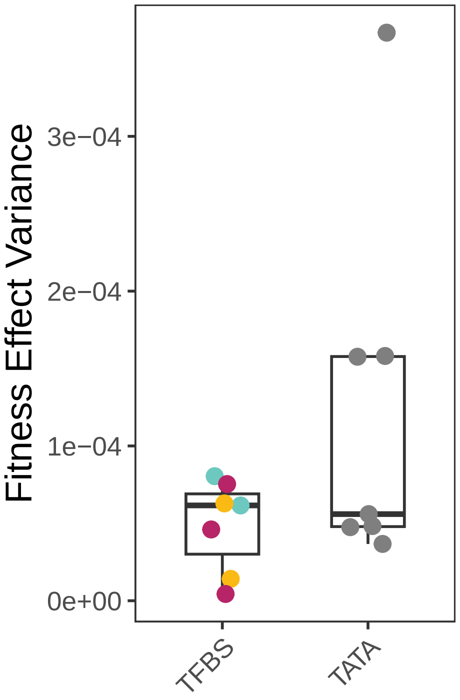
